# Whole genome sequence improvement with pedigree information and reference genotypic profiles, demonstrated in outcrossing apple

**DOI:** 10.1101/2024.08.08.607141

**Authors:** Stijn Vanderzande, Cameron Peace, Eric van de Weg

**Affiliations:** Plant Breeding, Wageningen University & Research, P.O. Box 386, NL-6700 AJ Wageningen, The Netherlands; Department of Horticulture, Washington State University, Pullman, Washington WA, USA

**Keywords:** R script, genome assembly quality, SNP array, *Malus domestica*

## Abstract

Understanding the quality of a whole genome sequence (WGS) is important for its further use. Most WGS quality evaluations are based on bioinformatic quality metrics such as the N50 score, BUSCO score, and number of contigs and scaffolds present, yet genetic information considering principles of inheritance could be used to evaluate and improve assembly and phasing. Furthermore, WGS and genome resequencing data of related individuals could provide useful information when large chromosomal segments are shared with the target individual through common ancestry. Here, we show how high-quality, phased, genome-wide genotypic information is useful to evaluate the quality of a WGS. We provide an R-tool to routinely conduct such quality evaluations. The script also provides a method to accurately determine the WGS positions of reference SNP markers, which is needed for integration of SNP array-based genotypic data sets with WGS data, and the identification and comparison of segments across WGSs that are shared by descent. Finally, we provide suggestions on how such sharing can be used to evaluate and improve new WGSs. The approach is demonstrated in apple, for which improvements in WGS quality are evident from the first collapsed WGS with many inconsistencies in genetic marker order and genotype scores, through well-assembled haploid WGSs, to incorrectly and correctly phased diploid WGSs. This study shows that homozygous regions might need extra attention in phased WGSs and that further improvements to phased WGSs can be achieved by grouping chromosomes of single parental origin into the same haplome.

## Introduction

Whole genome sequences (WGSs) are valuable resources for genomic discoveries and advancements, and thanks to decreased sequencing costs and new sequencing techniques, more WGSs are becoming available. The availability of one or more WGSs enables the detection of genetic variations such as single nucleotide polymorphisms (SNPs) within a crop species, facilitates the identification and characterization of key genes responsible for traits of interest, aids in understanding the complex genetic networks within a crop, and provides insights into the evolutionary history, genetic diversity, and population structure of a crop (Barabaschi et al. 2012; Bolger et al. 2014; Peace et al. 2019; Aranzana et al. 2019). Where the generation of the first whole genome sequences cost millions of dollars and took several years, new technologies have greatly decreased the cost and time needed to obtain a new WGS for one’s crop of interest (Jiao and Schneeberger 2017; Kersey 2019). Therefore, it is expected that more WGSs will be created for a wide range of species and individuals. However, to date many species still have no or a limited number of WGSs available, and those existing are often of unknown or limited quality.

As multiple WGSs become available, understanding the quality of and information contained within each WGS is important to understand its utility, limitations, and how it compares to other available WGSs. Although next-generation sequencing has enabled the generation of vast amounts of sequencing data, its short reads can lead to difficulties in accurately resolving repetitive regions, gene families, and complex genomic structures, resulting in fragmented or incorrect assemblies (Kersey 2019; Liao et al. 2019). For heterozygous and polyploid species, the presence of variations between homologous chromosomes and sequence similarity between homoeologous chromosomes further restrict the assembly using short reads (Michael and VanBuren 2015). Consequently, most early WGSs collapsed together the homologous copies of every chromosome into a haploid mosaic reference sequence rather than providing the variation present in the individual. Recent long-read technologies have enabled to span repetitive regions and facilitate the reconstruction of complex genomes, improving the assembly and resolving haplotypes at the chromosome level (Jiao and Schneeberger 2017; Michael and VanBuren 2020; Sharma et al. 2022). Yet long-read technologies initially showed a genotyping error rate of about 7% - 10%, relying on short-read technologies to identify and correct these errors (Koren et al. 2012). The overall quality of a WGS is mostly evaluated using various bioinformatic metrics (Gurevich et al. 2013; Whibley et al. 2021). For example, the N50 score is used to assess contiguity, measuring at which contig length half of the total assembly length is contained in contigs of that length or longer. The BUSCO score estimates the completeness of a genome assembly by searching for a set of conserved, single-copy genes. Other reported quality statistics are the coverage used, the number of scaffolds and contigs present, the genome size estimation, and repeat content quantification.

In plants, available genetic information has rarely been used to evaluate or improve the quality of a WGS. Part of the reason for this unfulfilled opportunity might be due to the relatively low marker density, inaccuracy in local marker order, and lack of published information on correctly phased genome-wide genotypic information in the case of outbreeding crops. The use of genetic information, accounting for principles of inheritance, has mostly been restricted to genetic maps composed of genome-wide genetic markers. Genetic maps are employed to scaffold *in silico* assemblies as a final step in constructing pseudomolecules for the generation of good to high-quality chromosomal-level WGSs (Ren et al. 2012; Deokar et al. 2014; Di Pierro et al. 2016; Verde et al. 2017; Bernhardsson et al. 2019; Li et al. 2019; Catchen et al. 2020). They support the correct ordering and orienting of scaffolds. The orienting relies on the presence of genetic markers that are located in two or more genetic bins within a single genomic scaffold; their relative placement in the genetic map enables the correct orientation of the scaffold. Additionally, positional information for multiple markers per scaffold helps assess the scaffolding process, facilitating the identification and resolution of potential assembly errors such as chimeric scaffolds (Drost et al. 2009; Bartholomé et al. 2015). For example, if two markers derived from a single scaffold appear to anchor to distinct regions within a linkage group or to different linkage groups, it suggests that the involved contigs have been erroneously joined during *in silico* assembly. Key factors that determine the effectiveness of a genetic map to assess a WGS’s quality are the number of true meiotic recombinations observed to create the genetic map, the distribution and density of the markers on the map, the accuracy of the genotypic data, and the size distribution of the scaffolds. Other genetic information such as parent-child relationships have been sometimes used in ‘‘trio binning” and “gamete binning” (Campoy et al. 2016; Koren et al. 2018; Shi et al. 2019; Zhang et al. 2024). Such binning requires either the parents or gametes to be sequenced to enable phasing the target WGS, which multiples the DNA sequencing cost. A more accessible and cost-efficient alternative could be reference linkage phase genotypic information from low-density genome-wide data of multiple genetically related individuals generated by SNP arrays, genotyping-by-sequencing, and/or low-coverage resequencing. Such data is often already available and its phasing information also enables the binning of contigs according to parental origin, especially when larger contigs are created using long-read sequence technologies. Yet the use of such information has been limited for WGS creation.

Apple (*Malus domestica*) with its series of released WGSs since 2010 from various institutions and of various cultivars, represents an illustrative case where WGS creation could benefit from the integration of available genome-wide genotypic information. The initially released apple WGS had some scaffolding and assembly errors due to the short-read DNA sequencing technologies employed, only a sparse SSR-SNP linkage map available at the time, and the high heterozygosity and allopolyploid-like nature of the sequenced individual, cultivar Golden Delicious (Velasco et al. 2010). Subsequent assembly versions of the ‘Golden Delicious’ WGS were improved using the Golden Delicious v2.0 based high-quality iGL genetic-physical map, yet 6% of the markers still showed inconsistencies in collinearity between genetic map and WGS (Di Pierro et al. 2016). Subsequent assemblies of the ‘Golden Delicious’ double haploid line 13 (GDDH13) and ‘Hanfu’ trihaploid line 1 (HFTH1) WGSs were improved through a complexity reduction by using di- and tri-haploids (Daccord et al. 2017; Zhang et al. 2019). The GDDH13 WGS took advantage of the iGL genetic-physical map, though only at a very late stage of its assembly, while the HFTH1 WGS also took advantage of emerging new long-read sequencing technologies, thereby becoming the most reliable apple reference WGS of that time (Skytte af Sätra et al., in prep). The further use of long-read technologies resulted in the first phased WGS of a diploid apple, of the cultivar Gala, which consisted of two haplome sequences, one for each chromosome set (Sun et al. 2020). Several other phased WGSs have been released since then for the diploid cultivars, including Honeycrisp, Fuji, and Golden Delicious (Khan et al. 2022; Su et al. 2024; Peng et al. 2024). However, no genetic information on their parents, grandparents, or other ancestors was used to ensure that these WGSs were correctly phased and that their chromosomes were ordered into two parent-specific haplomes. Such phasing and ordering would be useful for the revealing of trait-associated DNA polymorphism, checking and improving WGS quality, and revealing genetic processes such as mutation and gene conversion rates through comparisons among genome segments that are Identical By Descent (IBD). Yet such genetic information is available.

In parallel with sequence technology developments, affordable high-throughput genome-wide genotyping technologies became available, which for apple resulted in high-quality marker-based DNA profiles (i.e., genotypic information on multiple crop individuals) and high-density genetic linkage maps based on SNP array markers (Di Pierro et al. 2016; Howard et al. 2017; Vanderzande et al. 2019). To date, three SNP arrays have been commercially available in apple: the 8K, 20K, and 480K Infinium SNP arrays (Chagné et al. 2012; Bianco et al. 2014, 2016). Both the 8K and 20K apple SNP arrays have been extensively used to generate high-quality phased genome-wide DNA profiles for more than 5000 historic and modern apple cultivars and tens of full-sib families totaling many thousands more individuals (Howard et al. 2018). Several of these data sets are already publicly available and have been used for the validation and reconstruction of numerous cultivar pedigrees (Muranty et al. 2018; Vanderzande et al. 2019; Howard et al. 2021), partly due to the extensive level of data curation involving application of inheritance-based genetics principles (Vanderzande et al. 2019). These data sets also resulted in the 2^nd^ generation genetic-physical map in apple, the 15K-iGW map (Skytte af Sätra et al., in prep). To date, 15K-iGL is still the shortest published SNP map in apple while including the largest number of meioses (3172 meioses in 1586 offspring from 21 full-sib families) and markers (15,403 SNPs). Such data also helped with scaffolding and orienting contigs of haploid WGSs for individuals derived from important commercial cultivars (GDDH13) as well as the first presumedly phased WGS of a diploid cultivar (Gala) (Daccord et al. 2017; Sun et al. 2020). With the rise of multiple phased WGSs and their mutual comparison in pangemome studies (Sun et al. 2020; Wang et al. 2023), the use of genotypic data and inheritance-based genetics principles in WGS data quality control could be beneficial. For example, both ‘Gala’ and ‘Honeycrisp’ are descendants of ‘Golden Delicious’ (Evans et al. 2011; Howard et al. 2017), a cultivar on which previous diploid and haploid WGSs were based (Fig. 1). Thus, it is expected that genomic segments are shared among ‘Golden Delicious’, ‘Honeycrisp’, and ‘Gala’ and that, for those segments, the DNA sequence should be identical. Similarly, through the cultivar Delicious, ‘Gala’ is also closely related to ‘Fuji’, ‘Hanfu’ and ‘Hanfu’s haploid derivative, HFTH1 and thus these individuals are also expected to extensively share genomic segments (Fig1). Available genotypic data can identify these shared segments (Howard et al. 2021) to enable comparison of their sequences across WGSs.

**Figure 1:**
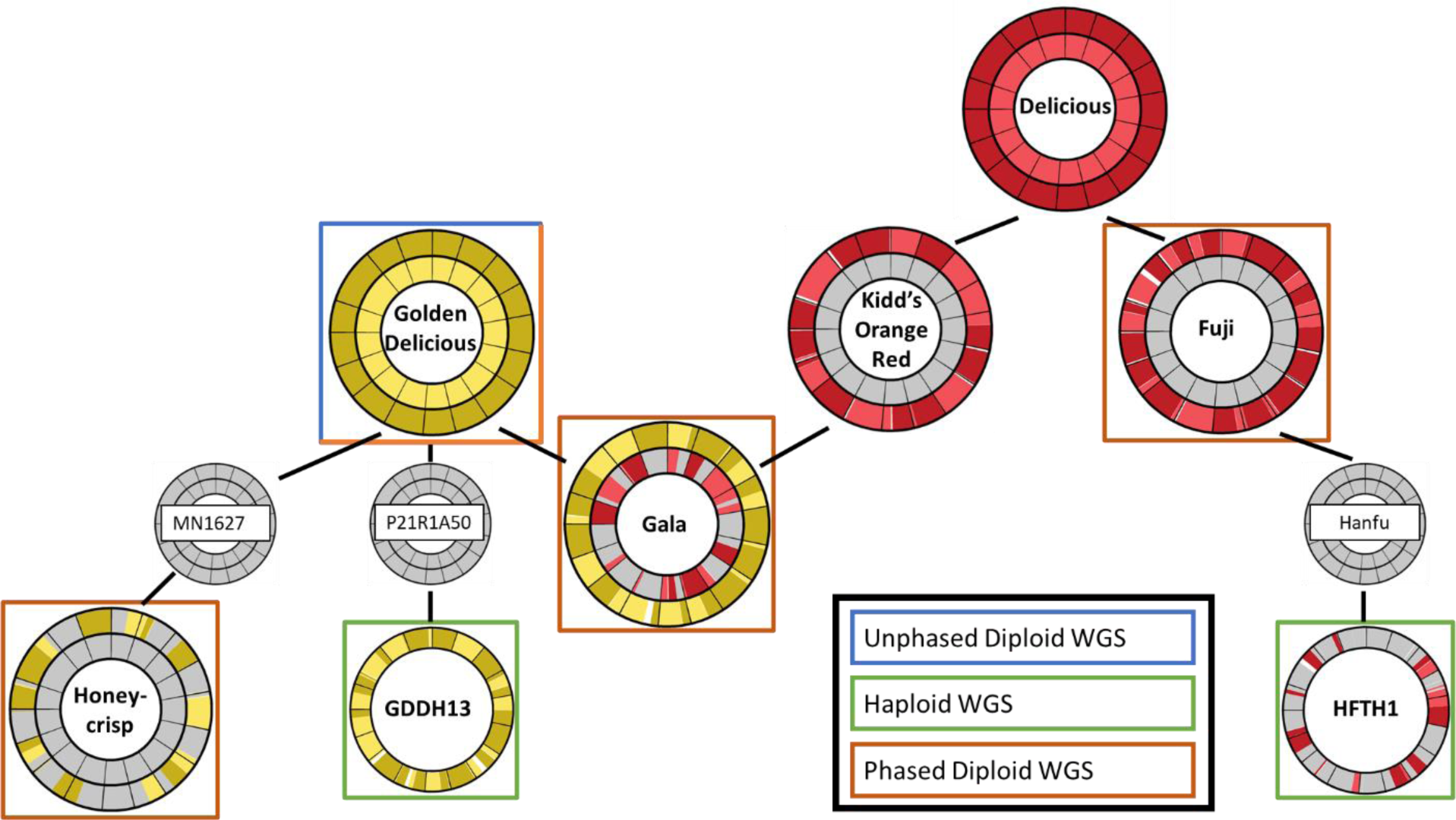
Relationships between the apple individuals whose WGS is available (in boxes). Chromosomal segments originating from Golden Delicious and Delicious are highlighted in yellows and reds, respectively, and identified from phased 20K SNP array data. The presented pedigree relationships are from published breeding records, validated with DNA markers. The origin of chromosome segments are from this study. The second parents of most individuals are not shown for simplicity (only GDDH13, GDDH13’s single parent, and HFTH1 have only one parent).

Integrating DNA sequence and assembly data of WGSs with genetic information from pedigreed and high-quality phased SNP array-based DNA profiles would enable validation or clarification of the parental origin of most to all genome segments in a WGS and the detection of shared IBD sequences across WGSs. Initial comparisons of the diploid ‘Gala’ WGS with its 8K SNP array DNA profile identified assembly issues and that the WGS was only phased correctly at a local level (Vanderzande and Peace 2023), yet no tools exist to conduct such comparisons routinely. No parental information was used for the diploid WGSs of either ‘Gala’ or ‘Honeycrisp’. Thus, it is unclear whether the later ‘Honeycrisp’ WGS improved on assembly and phasing of the earlier ‘Gala’ WGS. Furthermore, shared segments of WGSs have not been compared from an IBD perspective. Therefore, the objectives of this study were two-fold, firstly to demonstrate how use of available genetic information can enhance the quality of current and new WGSs and secondly to provide software tools to routinely evaluate the genetic quality of WGSs using genome-wide DNA profiles and visualize the outcome.

## Material and Methods

### Whole genome sequences

Five WGSs of cultivated apple (*Malus domestica*) were available at the start of this study and therefore included: Golden Delicious Genome v3.0.a1 (GDv3; (Velasco et al. 2010; Di Pierro et al. 2016)), GDDH13 v1.1 (Daccord et al. 2017), HFTH1 v1.0 (Zhang et al. 2019), phased diploid Gala v1.0 (Ga; (Sun et al. 2020)), and the phased diploid Honeycrisp v1.0 (HC;(Khan et al. 2022)). The Ga and HC WGSs each had two haplome sequences representing the two homolog sets of each individual. For the Ga WGS, these haplome sequences were labeled in the assembly as “A” or “B” and referred to hereafter as Ga_A and Ga_B, respectively. For HC, the haplome sequences were labeled in the assembly as “hap1” and “hap2”, referred to hereafter as HC_h1 and HC_h2, respectively. Each WGS and haplome sequence was downloaded from the Genome Database for Rosaceae (Jung et al. 2019). In view of readability, we did not include the version number in our WGS acronyms.

### Genome-wide genotypic data

The 15K-iGW map and 20K SNP array were used for the construction of SNP array-based DNA profiles (or SNP array profile) for a series of cultivars. The 15K-iGW’s genetic positions are based on the genetic positions of the previous iGLMap (Di Pierro et al. 2016), with genetically co-localizing markers now being ordered based on their position in the recent HFTH1 and GDDH13 WGS reference genomes (Skytte af Sätra et al., in prep), replacing their previous order from the GDv2 draft WGS. Unique genetic positions for each marker in the 15K-iGW were determined by Skytte af Sätra et al. (in prep) as follows: regression of the iGLMap genetic positions to the corresponding HFTH1 positions resulted in a set of polynomials, one per chromosome, for the creation of standardized virtual linkage maps in apple. The application of these polynomials on the 15.4K SNPs of the iGLMap resulted in the current 15K-iGW map, which holds 15.403 SNPs from the 20K SNP array, and which will be further referred to in this study as the iGW map.

Phased SNP genotypic data was obtained with the 20K SNP array (Bianco et al. 2014) as part of other studies (Allard et al. 2016; Di Pierro et al. 2016; Howard et al. 2018, 2021). A large genotypic data set was created by these studies, including SNP array data for the WGS individuals ‘Golden Delicious’, ‘Gala’, and ‘Honeycrisp’ as well as available offspring and progenitors of any WGS individual such as ‘Grimes Golden’, ‘Kidd’s Orange Red’, ‘Cox’s Orange Pippin’, ‘Delicious’, ‘Winesap’, ‘Fuji’, and ‘Ralls Janet’. SNP array data in this large data set was previously error-cleaned and phased according to Vanderzande et al. (2019) as part of other studies.

### BLAST protocol optimization for determining of marker locations in each WGS

A local BLAST data set was created for each WGS using the “makeblastdb” command (Camacho et al. 2009). In an initial round, the optimal BLAST procedure was determined using the HFTH1 WGS and genomic locations as determined for the creation of the iGW (Skytte af Sätra et al., in prep). BLAST results for both flanking sequences (i.e., a continuous sequence of at least 121 nt (=60+1+60) across both sides of the SNP) and 50 nt long probe sequences (i.e., on one side of the SNP) as provided by the SNP array designers (Bianco et al. 2014) were considered. Each of these sequences were used in two BLAST protocols against HFTH1, ‘megablast’ and ‘blastn’. Finally, when flanking sequences were used, a third BLAST protocol, blastn-short, was also performed on truncated flanking sequences of 31 nts containing only the first 15 bp on each side of the SNP. This third analysis was performed because gaps in the alignment around the target nucleotide can hamper the determination of the correct SNP location in the respective WGS. No blastn-short was run for the probes as gaps in their alignment did not hinder the accurate determination of genomic locations. All BLAST runs reported only hits with 80% identity or higher with the “-perc_identity 80” tag and output files were generated with the “-outfmt 6” tag. This resulted in four protocols: 1) megablast on flanking sequences + blastn-short, 2) blastn on flanking sequences + blastn-short, 3) megablast on probes sequences, and 4) blastn on probe sequences. The results of each protocol were analyzed by the newly developed R function ‘*blastAnalysis*’, as described below, and the resulting genomic positions were compared to those determined for the creation of the iGW (R Core Team 2018, Skytte af Sätra et al., in prep). For the protocols with flanking sequences, chosen hits for which the blastn-short still had gaps were not considered as ∼70% of these hits resulted in incorrect genome locations. The BLAST protocol that resulted in the most consistent result with those of the iGW was used in further analyses of all WGSs.

### Determination of marker locations in each WGS

As part of this study, a new function, named ‘*blastAnalysis*’, was developed in R (R Core Team 2018) to accurately determine genomic locations for each SNP and each WGS, considering information on genetic positions of segregating SNP-probes in the iGW genetic-physical map (File S1, Fig. S1, Fig. S2). The function uses results from up to two BLAST runs: a first “full BLAST” run on longer target sequences which are more likely to contain gaps in the alignment that can hamper the determination of the genomic location and a second, optional, “short BLAST” run on shorter target sequences to avoid gaps in their alignment. The function first analyzes the results from the full BLAST as described below to determine the most likely general location for each SNP. Then, if the results from a short BLAST are provided as an input, it optimizes each location in case of gaps in the alignment as described below.

The ‘*blastAnalysis*’ function filters the full BLAST results as follows: In a first step, only hits are maintained for which the alignment either contains the target SNP or ends on a nucleotide immediately flanking the target SNP (Fig. S1). For example, the probe’s final 3’-end nucleotide should be included as it is immediately flanking the target SNP. To do so, the SNP’s position in the corresponding query sequence is compared against the start and end of the alignment of the query for each hit. This position in the query sequence is provided by a user-provided “snpInfo” input file as this position can differ for each query sequence. In addition, this location can be immediately upstream or downstream of the provided sequence in case of probes. In a second step, any alignment shorter than a user-defined length is also removed (15bp was used in this study). After these filter steps, the corresponding SNP’s WGS location for each BLAST hit is determined, considering the alignment orientation and location of the SNP in the query sequence. Then, any SNP with a single BLAST hit is assigned the corresponding WGS location of that single BLAST hit. Further analyses are needed for SNPs with multiple hits, as the best hit is not necessarily the one segregating in the germplasm, especially in apple (and other crops) for which a whole (or partial) genome duplication occurred in its evolutionary history (Velasco et al. 2010) and therefore multiple hits are commonly observed.

For the remaining SNPs with multiple BLAST hits, the provided genetic map was used to determine the most likely hit using the helper function ‘*mapMatchingHit*’. This function uses the WGS locations of genetically mapped single-hit SNPs as anchor points and compares the WGS location of these anchor points to all possible WGS locations for the multiple-hit SNPs. For each multiple-hit SNP, only anchor points within a user-defined distance on the genetic map (2.5 cM in this study) are considered to evaluate its possible WGS location. To account for small local rearrangements as well as incorrect marker order within genetic bins of the genetic map (e.g., where no informative recombination is present between markers to inform their correct order), the WGS region to consider for BLAST hits is determined in three steps. First, the nearest upstream and downstream anchor points are used to determine a general WGS region (described further below). Second, up to a user-defined number (three in this study) of upstream and downstream anchor points whose location corresponded to this general WGS region are used to determine the region’s limits (described further below). Third, the region’s limits can be further extended by a user-defined distance (100,000 bp in this study). When determining the general WGS region, flanking anchor points can be assigned to different WGS chromosomes (due to assembly issues). Therefore, multiple scenarios are considered to determine this general region for the BLAST hits of multiple-hit SNPs: the assigned chromosomes of flanking anchor points are 1) consistent with each other and with the genetic map, 2) consistent with each other but not with the genetic map, 3) not consistent but one is consistent with the genetic map, and 4) not consistent and neither is consistent with the genetic map. In the first two scenarios, the flanking markers are used to define the general WGS region. In scenario 2, however, when no hits are found in the determined general WGS region, anchor points within a user-defined distance (in cM) of the target SNP are searched that match the chromosome of the genetic map and these anchor points are then used to determine the general WGS region. Prioritizing a non-target chromosome is done to account for assembly issues where a contig and all the SNPs on this contig are assigned to an incorrect chromosome. For scenario 3, priority is given to the target chromosome based on the genetic map and a more distant anchor point is searched on the side of the inconsistent anchor point to determine the general WGS region. Only when no BLAST hits are found in the resulting general WGS region on the target chromosome, anchor points on the alternative chromosome are considered to determine the general WGS region. For scenario 4, two general WGS regions, one for each chromosome, are determined and only if BLAST hits match only one of the two regions is that region considered. If no hits are found in either of the two general WGS regions, anchor points within the user-defined distance of the target SNP are searched that match the chromosome of the genetic map to determine the general WGS region. Once a general region is established, this region is further refined by a user-defined number of upstream and downstream anchor points corresponding to the general region. The start and end of this refined WGS region is determined by the lowest and highest WGS position of all considered anchor points even if the anchor point with the lowest or highest WGS position is located downstream or upstream of the target SNP according to the genetic map, respectively. This step was again done to account for possible inversions of the WGS compared to the genetic map. After an optional extension of this refined WGS region by a user-defined distance (in bp), all BLAST hits of the multiple-hit SNP within the final WGS region are considered to be consistent with the map. If only one hit is found, the SNP has its corresponding WGS location assigned. If multiple hits are retained, it is checked whether a single hit is located between the anchor points closest in the genetic map to the target SNP (as multiple SNPs on either side were used to define the target region). If so, the corresponding WGS location of that hit is used and the rest discarded. If not, all retained hits in the determined WGS region are used in subsequent rounds. If no hits are retained, all BLAST hits available at the start of this filter round are used in the next round.

Any of the remaining still-unassigned SNPs are subjected to two rounds of selection as part of the ‘b*lastAnalysis*’ function (Fig. S1). In the first round, only hits within 10% of the best identity of each SNP are retained and in the second round this subset is further filtered to only retain SNPs within 5% of the best bitscore per SNP. The 10% and 5% thresholds are chosen as multiple hits can have very close quality scores and the optimal hit is not always the one with the best score. These thresholds also enable to define relative cut-offs based on the best result for each SNP rather than applying a single cut-off value for all SNPs. In each round, if only one hit is retained for a SNP, that BLAST hit is considered to provide the most likely WGS location. Then, after each selection round, the reference map and resolved SNPs are compared to the BLAST hits of remaining unresolved SNPs as described above. After these two filter rounds, any remaining unresolved SNP is categorized into one of two groups: one group for which all remaining hits of the SNP are located on a single chromosome (single chromosome multi-hit SNPs) and another group for which the remaining hits are located on multiple chromosomes (multi-chromosome multi-hit SNPs) (Fig. S1).

In case of gaps in the alignment, the locations determined from the full BLAST results are then updated using the short BLAST results by the ‘b*lastAnalysis*’ function, if provided. For any BLAST hit where the alignment ended on the target nucleotide or either nucleotide immediately neighboring the target SNP, the genomic location could be accurately determined from this end in the alignment, even in the presence of gaps. Therefore, no short BLAST is used to update the location of these SNPs. Practically, this means that for probe sequences, where the target SNP is either at the final nucleotide of the probe’s 3’ end or immediately neighboring this nucleotide, no updates for gaps in the alignment are needed. When an update in genomic location is needed because of gaps, the short BLAST results of those SNPs are filtered to match the WGS region of the chosen full BLAST hit. If multiple short BLAST hits match the WGS region of the full BLAST hit, the short BLAST hit for which the SNP WGS location is closest to the full BLAST hit SNP WGS location is chosen.

### WGS DNA profiles versus SNP array profiles

For each WGS, a WGS-based DNA profile (WGS profile) was created similar to the SNP array profiles using R. Only SNPs with a single assigned location within the considered WGS, whose location could be accurately determined (i.e., no gaps in alignment or target SNP at end of alignment), and whose WGS location was consistent with the provided genetic map were used. For these SNPs, a new R function, ‘*retrieveSnpAllele*’, retrieved the nucleotide allele present at the SNP’s location in the respective WGS (Fig. S2). This function made use of the ‘*fastaToTabular*’ R function obtained from Github (https://github.com/lrjoshi/FastaTabular) to load the WGS into R. Then, the nucleotide was translated to the SNP array calling format where “A” and “T” nucleotides are coded as “A”, and “C” and “G” nucleotides are coded as “B”. Any SNPs that were not considered had their alleles set to missing for the WGS profile.

The WGS profiles of GDv3, Ga, and HC were then compared to the SNP array profiles of ‘Golden Delicious’, ‘Gala’, and ‘Honeycrisp’, respectively (Fig. S2). Because GDv3 is an unphased collapsed WGS, only a mosaic haploid WGS profile could be compared to the phased diploid SNP array profile. In contrast, the phased Ga and HC WGS provided two haploid WGS profiles, one for each haplome and their combined-haplome WGS profiles were compared to their SNP array profiles. No SNP array profiles were available for the ‘GDDH13’ and ‘HFTH1’ individuals to compare to their respective WGS. Instead, the SNP array profile of GDDH13’s sole grandparent ‘Golden Delicious’ was used. For ‘HFTH1’, the SNP array profile of ‘Fuji’, the only available parent of ‘Hanfu’, and thus one of the two grandparents of ‘HFTH1’, was checked for shared alleles with ‘HFTH1’. Because the HFTH1 WGS was not compared against its own SNP array profile but that of ‘Fuji’, a high proportion of reported inconsistencies was expected for genomic regions originating from the other parent of ‘Hanfu’, ‘Dongguang’. First, an R function checked for inconsistencies in alleles present between both profiles, (i.e., “*AA*” in the SNP array profile versus “*AB*”/“*BB*” for phased WGS or versus “*B*” for a single haploid WGS). Null alleles were not considered because various biological reasons for null alleles exist (e.g., additional SNPs near the 3’ end of the probe or absence of the target sequence). Instead, null alleles in the reference profile were set to missing to avoid causing inconsistencies. SNPs with inconsistencies were then reported and when inconsistencies were found, the corresponding WGS profile alleles were set to missing and reported before determining allelic origin. Then, the ‘*determineAlleleOrig*’ R function compared the alleles present in each (haplome) WGS to phased SNP data to assign maternal or paternal origin (Fig. S2). Finally, multiple R functions were used for the visualization of the parental origins of alleles in each haplome, the genetic location of SNPs with allele inconsistencies with the reference SNP array profile, regions of homozygosity based on the SNP array profile, and the proportions of SNPs that did not result in a single location in the WGS, that were located on unanchored regions, and whose location was inconsistent with the provided genetic map (Fig. S2).

### Comparison of shared segments across WGSs

The WGS profile of GDDH13 and the SNP array profiles of ‘Gala’ and ‘Honeycrisp’ were used to manually determine what genomic segments these individuals shared from their common ancestor, ‘Golden Delicious’. Similarly, the WGS profile of HFTH1 and the SNP array profile of ‘Gala’ were compared to manually determine which genomic segments they shared from their common ancestor, ‘Delicious’. In case of ‘HFTH1’, only shared segments longer than 5, 10, and 15 cM were considered to be inherited by ‘HFTH1’ from its grandparent ‘Fuji’ because genotypic data for its parent ‘Hanfu’ and its grandparent ‘Dongguang’ were missing for accurate pedigree-tracing of inherited segments. This also means that the exact delimitation of ‘Fuji’-shared haplotypes might be slightly overestimated as SNP alleles originating from ‘Dongguang’ near the transition between ‘Fuji’ and ‘Dongguang’ segments that are identical to SNP alleles of ‘Fuji’ might be incorrectly assigned as originating from ‘Fuji’. Because these shared segments originated from a common ancestor, the sequence of these segments were expected to be (almost) identical across the corresponding WGSs. As a case study, a genomic region was searched for which ‘Gala’ shared both homologs with other WGSs, one from ‘Golden Delicious’ and one from ‘Delicious’, and for which the phasing was correct for all phased WGSs. As a second case study, a region of ‘Honeycrisp’ on chromosome 7 was chosen that was homozygous based on SNP array data. For this homozygous region, the genome sequences were also expected to be the same. The segments of this region of ‘Honeycrisp’ were inherited from ‘Golden Delicious’ and ‘Northern Spy’ (Howard et al. 2017) and as these individuals are not (yet) connected by known pedigrees, this expectation on homozygosity is based on Identity By State of the SNP profiles.

MUMMER3.23 (Kurtz et al. 2004) was used to compare sequences between WGS segments, in a Linux environment created by Cygwin64 (https://www.cygwin.com/) on a Windows 10 computer. The Mummer source file was adjusted to avoid errors in the program by removing “defined” on lines 884, 894, 981, 991, 1034, and 1044 (https://github.com/bioconda/bioconda-recipes/issues/1254). The “nucmer” function of MUMMER was used for pairwise WGS segment comparisons with the following command: nucmer -maxmatch -c 500 -l 100 *WGS1-segment.fasta WGS2-segment.fasta* where “*WGS1-segment.fasta*” and “*WGS2-segment.fasta*” were the names of the fasta files containing the respective WGS sequences. For the comparison across all shared segments, these files contained the sequences for all shared segments. The “dnadiff” function of MUMMER was used to summarize the alignment results and determine the proportion of identity between shared segments and the “mummerplot” function of MUMMER was used to visualize the alignments.

## Results

### R functions

A total of 24 R functions were created to enable routine analysis of WGS assemblies and phasing using genome-wide marker DNA profiles (File S1, Table S1). A single wrapper function, ‘*phasedWgsVsSNP*’, was successfully developed that accepts BLAST outcomes, WGS assembly, and reference marker DNA profiles to analyze a WGS assembly without manual intervention required by the user. The wrapper can also only determine the marker’s location, check for inconsistencies with the genetic map, and extract alleles at the determined marker locations if no reference individual is available. Alternatively, the wrapper can run only the comparison with a reference individual and visualize the outcome, for example when marker location and WGS profile were already previously determined. This is useful because determining the marker locations in the WGS is the rate-limiting step and does not need repeating when comparing the WGS against multiple reference individuals. The type of analyses, as well as various parameters and input files can be set through a single parameter file accepted by the wrapper function (Table S2, File S2). The wrapper function contains a pipeline combining twelve other functions that can each also be used as stand-alone functions (Fig. S2, Table S1). These twelve functions accept external input files such as BLAST results as well as output from other functions used earlier in the pipeline. The pipeline enables manual adjustment of intermediate output before continuing analyses. With BLAST results available, an analysis comparing a single haploid WGS of ∼650 Mb with a single SNP array profile of ∼10K markers took ∼7.5 minutes, while the comparison of a diploid phased WGS of ∼1300 Mb with a single marker profile of ∼10K markers took ∼14 minutes using the wrapper function on a Windows 10 system with 11^th^ gen Intel processor @3.2 GHz. These two comparisons created 15 and 24 output files for haploid and diploid WGSs, respectively, which can be adjusted if needed and used with individual functions as part of a more user-involved pipeline. Example input and output data files were also generated (File S2 and File S3, respectively). Below, the results of these functions is demonstrated when applied on five genetically related WGSs in apple and integrating phased SNP marker profiles and pedigree information.

### SNP WGS mapping with BLAST & filtering with genetic information

When examining the efficiency of different BLAST approaches, use of the ‘blastn’ protocol for both flanking and probe sequences resulted in more consistent unique hits and overall consistent results compared to use of the ‘megablast’ protocol (Table 1, Table S3). The use of probe sequences resulted slightly higher number of results consistent with previously determined HFTH1 positions compared to use of flanking sequence (Table 1). Finally, for SNPs with previously undetermined locations, use of flanking sequences resulted in detection of more unique locations consistent with the WGS location of flanking SNPs. Because of the slightly higher consistency and because probes better represent the actual DNA binding conditions on the SNP array, the ‘blastn’ protocol using probe sequences were used for the results of other WGSs described hereafter.

**Table 1:** Comparison of SNP locations obtained through the ‘blastAnalysis’ function versus previously determined SNP locations in the HFTH1 WGS. Results are presented for two BLAST protocols (‘megablast’ and ‘blastn’) on both SNP flanking sequences and SNP probe sequences. For flanking sequences, a second short-blast run was incorporated to avoid gaps in the alignment (as described in Materials and Methods). Per run, the number of consistent results are indicated for three categories (SNPs with a single unique WGS location without gaps in the alignment, SNPs with multiple plausible WGS locations within the same chromosome, and SNPs for which no WGS location could be determined) as well as the total number of consistently located SNPs. Also provided is the number of SNPs for which previously no WGS location could be determined (Skytte af Sätra et al., in prep) but for which the current analyses resulted in a single unique WGS location that is consistent with the WGS location of flanking SNPs. The classifications of the individual SNPs are presented for each of the four protocol-sequence combinations in Table S3

| Category of SNP location comparison outcome | Flanking sequence |  | Probe sequence |  |
| --- | --- | --- | --- | --- |
|  | megablast | blastn | megablast | blastn |
| Matching single hit, no gaps | 14604 | 14648 | 13867 | 14652 |
| Matching multiple hits, single chromosome | 65 | 56 | 53 | 58 |
| No matching hit | 85 | 4 | 135 | 24 |
| Total SNPs with consistent WGS location | 14754 | 14708 | 14055 | 14734 |
| Previously undetermined locations consistent with WGS location of flanking SNPs | 14 | 38 | 6 | 24 |

Considering results of the ‘blastn’ analyses of probe sequences on the other WGSs, both haploid WGSs and the HC haplome WGS had the highest number of SNPs for which WGS locations were determined unambiguously (Table 2, Table S4). Those WGSs also had considerably fewer SNPs with multiple BLAST hits within the same chromosome, while more than 10% of the SNPs belonged to this category for GDv3. Furthermore, the haploid WGSs and the HC haplome WGSs had among the fewest SNPs for which no BLAST hits could be found, although the GDv3 WGS had even fewer such SNPs, while Ga_B had the most by far. The number of reference SNPs that mapped to multiple chromosomes was similar across most WGSs and low (19–102 SNPs), except for Ga_B. For SNP location comparisons between WGSs and the iGW map, almost no inconsistencies were detected for the GDv3, HFTH1 and HC WGSs (<2%), some inconsistencies for GDDH13 and Ga_A WGSs (∼2.5%), and considerably more inconsistencies were detected for Ga_B WGSs (8.2%). SNPs without a unique location in the WGS or with inconsistencies between the WGS and the iGW map were generally spread throughout the WGS (Fig. 2). However, most such SNPs for the HC_h1 WGS were located at the proximal end of linkage group 2 of the iGW map (displayed by the long grey/red peak in the innermost circle of Fig. 2E).

**Figure 2:**
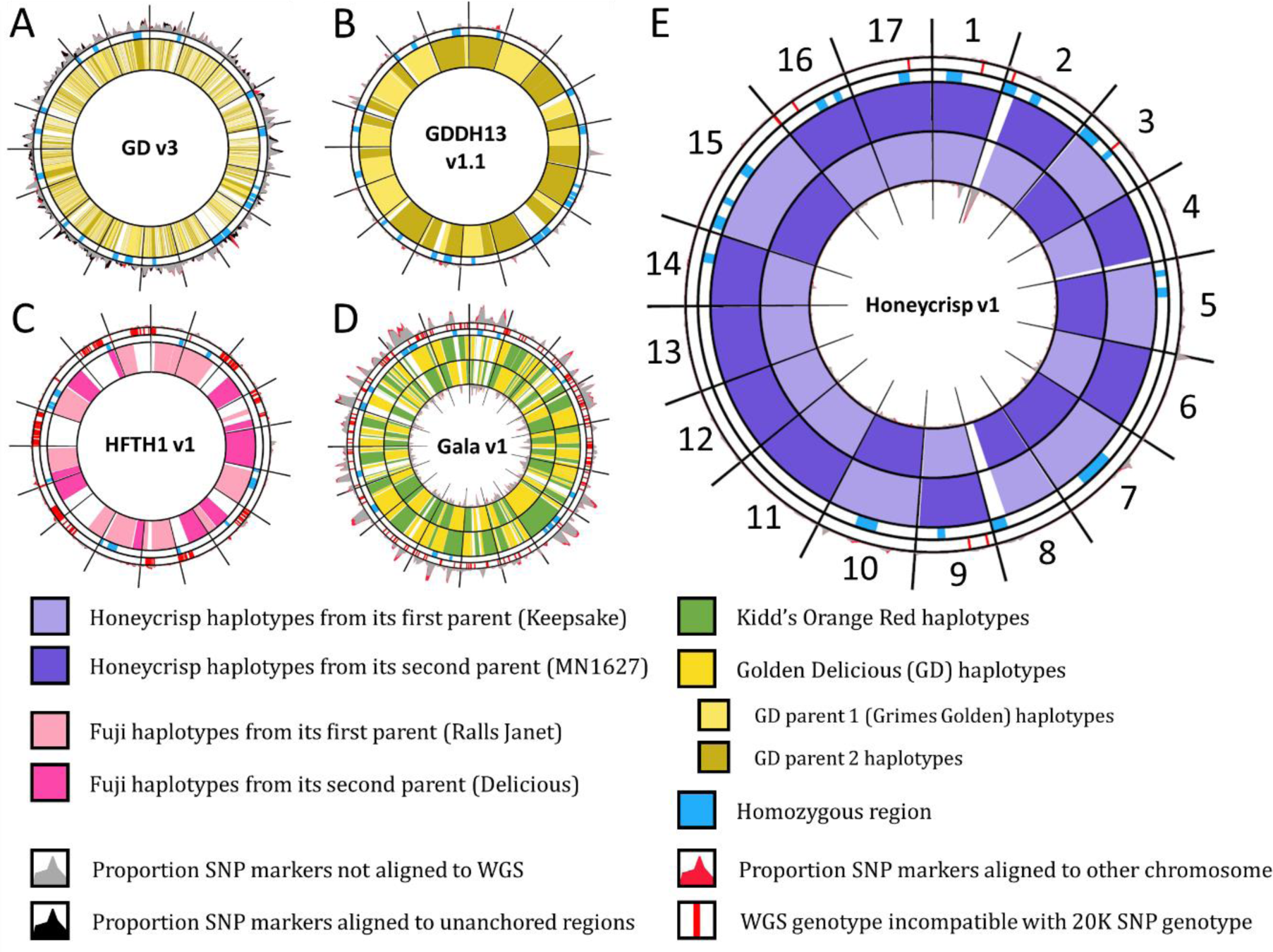
Parental origin of genome segments in the WGSs of A) GD, B) GDDH13, C) HFTH1, D) Ga, and E) HC. These parental origins are displayed in the widest circles per WGS. Both haplomes are shown for the two phased diploid WGSs, Ga and HC, with the first haplome (Ga_A and HC_h1, respectively) as the inner of the two circles and the second haplome (Ga_B and HC_h2, respectively) as the outer wide circle. For the haploid HFTH1, no parental information was available and only segments shared with grandparent ‘Fuji’ longer than 10 cM are displayed. Similarly to the circles displaying parental origin, the innermost circle of the phased WGSs represents the proportion of SNP markers per 5cM window that were either not aligned to the first haplome or had inconsistencies with the iGW genetic map when aligned to the first haplome, while the outermost circle represents the same for the second haplome. For all other WGSs, these SNPs are displayed in the outermost circle only. Of the two narrow outer circles, the inner displays homozygous regions and the outer genotype mismatches between SNP array profiles and the WGS.

**Table 2:**
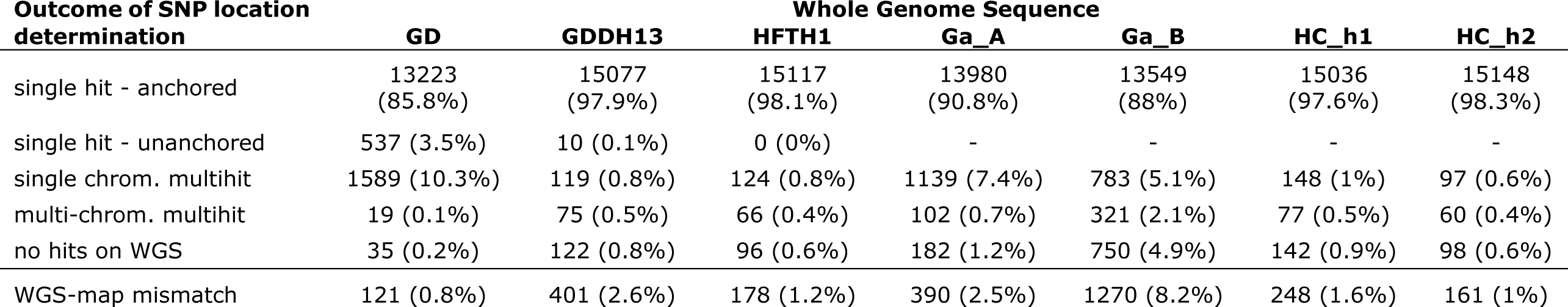
SNP location determinations within each WGS for SNPs of the iGW map. Where possible, SNPs for which a single location in the WGS could be determined were divided into those that aligned to one of the 17 chromosomes of an apple WGS (“single hit – anchored”) and those that aligned to unanchored contigs of a WGS (“single hit – unanchored”). SNPs that aligned to multiple chromosomes and for which a single alignment could not be chosen to be the most likely one were divided into those for which the multiple hits were on a single chromosome (“single-chrom. multi-hit”) and those spread over multiple chromosomes (“multi-chrom. multi-hit”). For each WGS, the number and proportion of inconsistencies between WGS location and genetic map are also provided. The classifications of the individual SNPs are presented for each of the four protocol-sequence combinations in Table S4

| Outcome of SNP location determination | Whole Genome Sequence |  |  |  |  |  |  |
| --- | --- | --- | --- | --- | --- | --- | --- |
|  | GD | GDDH13 | HFTH1 | Ga_A | Ga_B | HC_h1 | HC_h2 |
| single hit - anchored | 13223<br>(85.8%) | 15077<br>(97.9%) | 15117<br>(98.1%) | 13980<br>(90.8%) | 13549<br>(88%) | 15036<br>(97.6%) | 15148<br>(98.3%) |
| single hit - unanchored | 537 (3.5%) | 10 (0.1%) | 0 (0%) | - | - | - | - |
| single chrom. multihit | 1589 (10.3%) | 119 (0.8%) | 124 (0.8%) | 1139 (7.4%) | 783 (5.1%) | 148 (1%) | 97 (0.6%) |
| multi-chrom. multihit | 19 (0.1%) | 75 (0.5%) | 66 (0.4%) | 102 (0.7%) | 321 (2.1%) | 77 (0.5%) | 60 (0.4%) |
| no hits on WGS | 35 (0.2%) | 122 (0.8%) | 96 (0.6%) | 182 (1.2%) | 750 (4.9%) | 142 (0.9%) | 98 (0.6%) |
| WGS-map mismatch | 121 (0.8%) | 401 (2.6%) | 178 (1.2%) | 390 (2.5%) | 1270 (8.2%) | 248 (1.6%) | 161 (1%) |

### Allele origin and WGS phasing quality

For the Ga and HC WGSs, WGS genotypes could be determined for respectively 7194 and 9546 of the 9688 SNPs with reference genotypic data, while 2494 and 142 SNP genotypes, respectively, could not be used because no unique location was determined or because of a mismatch between the iGW map and WGS location for one or both haplomes of the corresponding WGS (Table S4). Of Ga’s determined WGS genotypes, 96.6% matched the ‘Gala’ marker genotype, while 3, 61 and 177 SNPs had two inconsistent alleles, one additional allele, or one missing allele in the WGS, respectively. Of the SNPs for which an additional allele was observed, 23 and 38 were observed in Ga_A and Ga_B, respectively. Another 17 SNPs for which only one allele could be extracted from the WGS data were incompatible with the 20K SNP profile (e.g., “A” allele in WGS versus “BB” for the marker genotype). For ‘Honeycrisp’, the consistency between HC WGS and SNP array genotypes was considerably higher (99.7%), with only 28 genotype mismatches observed of which 2 were due to an additional allele in the WGS (both found in HC_h2), 12 to a missing allele in the WGS, and 14 for which no alleles matched between WGS and SNP array data. No inconsistencies with the 20K SNP array profile were detected when only one allele could be extracted from the HC WGS. For the GDv3 and GDDH13 WGSs, 20 and 10 alleles respectively were inconsistent with the ‘Golden Delicious’ marker data (for example, a “*B*” allele in the WGS but an “*AA*” marker genotype). No mismatches were identified for HFTH1 WGS as no SNP data was available for HFTH1 or its diploid progenitor ‘Hanfu’, but four large genomic regions identical to ‘Fuji’ on chromosomes 1, 3, 5, and 17 were fragmented by a single mismatch with ‘Fuji’, indicating inconsistencies between WGS and SNP array data in these particular regions.

The parental origin of alleles was determined for all WGSs (Table S5). In the HC WGS, both parental homologs were represented across haplomes and each chromosomal sequence consisted of a single segment of maternal or paternal origin. However, each haplome consisted of a mixture of both maternal and paternal homologs, where HC_h1 had the expected maternal alleles for six chromosomes (3, 5, 7, 8, 10, 15) and paternal homologs for the other chromosomes and vice versa for HC_h2. In contrast, in the phased Ga WGS, each chromosome consisted of a mosaic of 3–11 segments (average 7.6) of alternating parental origin. Nevertheless, both parental alleles were still represented across both homologs. In Ga_A, the average size of a segment of single parental origin was 9 cM with 49.5% of all segments shorter than 5 cM and 20.8% being larger than 15 cM. Similarly, chromosomes of the GDDH13 consisted of a mosaic of 1–9 (average 2.7) segments of different great-grandparental origin, inherited from either parent of ‘Golden Delicious’ with 56.5% of the segments larger than 15 cM. There were only 14 (30.4%) segments shorter than 5 cM and these were mostly observed near the ends of the chromosome. Because of the haploid nature of GDDH13, only a single allele of either “maternal” or “paternal” origin was observed for each locus. For the HFTH1 WGS, 73%, 68%, and 53% of the WGS corresponded to the ‘Fuji’ genome when the threshold to assign a shared segment as originating from ‘Fuji’ was at minimum 5, 10, and 15 cM, respectively. Of these ‘Fuji’ segments in HFTH1 at the 5, 10, and 15-cM thresholds, 42%, 41%, and 38%, respectively, were consistent with alleles of ‘Rall’s Janet’ (the mother of ‘Fuji’). No small segments were observed as they were already excluded, yet many large segments were observed as was the case for GDDH13. In contrast, for GDv3 only 2 large segments (>15 cM; <0.1% of segments) were observed, with 99.4% of all segments of single parental origin smaller than 5 cM. Each chromosome of GDv3 consisted of 77–247 segments (average 119.9) and, because the WGS was collapsed, only one of either parental allele was ever observed and with both parental alleles equally represented.

In Ga_B, more issues were observed for homozygous regions of ‘Gala’, as identified by its SNP array profile, than for heterozygous regions. No unique location could be found for 69.2% of the SNPs in these homozygous regions, while only 9.6% of SNPs in heterozygous regions did not have a unique location. For other haplomes of the phased WGSs, the proportion of SNPs without unique location were 1–13.6% and 0.9– 9.3% for homozygous and heterozygous regions, respectively (Table 3). Furthermore, in Ga_B, homozygous regions contained 13% of SNPs whose WGS location did not correspond to the iGW map, while heterozygous regions only contained 1.1% of such issues. For other haplomes of the phased WGSs, the proportion of SNPs with inconsistencies between WGS location and iGW map were only 0.1–1.3% and 0.06–0.5% for homozygous and heterozygous regions, respectively (Table 3). No to moderate correlations were observed (range 0.06–0.75) between a region’s proportion of SNPs without unique location and the region’s proportion of SNPs whose WGS location did not match with the iGW map (Table 3).

**Table 3:** Number of SNPs in the phased WGS haplomes without unique location and with location inconsistent with the iGW map for homozygous and heterozygous regions, and their correlation.

| **SNP location issues observed** | **Ga_Av1** | **Ga_Bv1** | **HC_h1v1** | **HC_h2v1** |
| --- | --- | --- | --- | --- |
| *SNPs without unique location* |  |  |  |  |
| in homozygous regions | 145 (14%) | 740 (69%) | 66 (4.9%) | 13 (1%) |
| in heterozygous regions | 865 (9.3%) | 891 (9.6%) | 111 (1.2%) | 82 (0.9%) |
| *SNP locations inconsistent with map* |  |  |  |  |
| in homozygous regions | 14 (1.3%) | 142 (13.3%) | 17 (1.3%) | 2 (0.1%) |
| in heterozygous regions | 44 (0.5%) | 106 (1.1%) | 5 (0.06%) | 5 (0.06%) |
| *Correlation between a region's proportion of SNPs without unique location and SNPs with inconsistent locations* | 0.40 | 0.69 | 0.75 | 0.06 |

### Expected shared segments and sequence identity

Based on the SNP array profiles of ‘Golden Delicious’, ‘Gala’, and ‘Honeycrisp’ and the WGS profiles of GDDH13 and HFTH1, multiple shared segments were determined to be IBD among WGS individuals (Fig. 1, Fig. 3, Table S6). The total number of shared segments identified among ‘GDDH13’, ‘Gala’, and ‘Honeycrisp’ ranged from 13 to 23, covering 357 to 718 cM (Fig. 1). Of these shared segments, a total of 196 cM over 10 segments was determined to be shared among all three individuals and their shared ancestor, ‘Golden Delicious’ (Fig. 3). In the Ga WGS, 9 and 12 of these segments were in Ga_A and Ga_B, respectively, due to incorrect phasing (parent assignment) at the chromosome level. ‘Gala’ and ‘HFTH1’ shared 5 segments covering 106 cM via their common ancestor ‘Delicious’ (Fig. 3), which were further divided into 16 smaller segments because of incorrect phasing of the Ga WGS.

**Figure 3:**
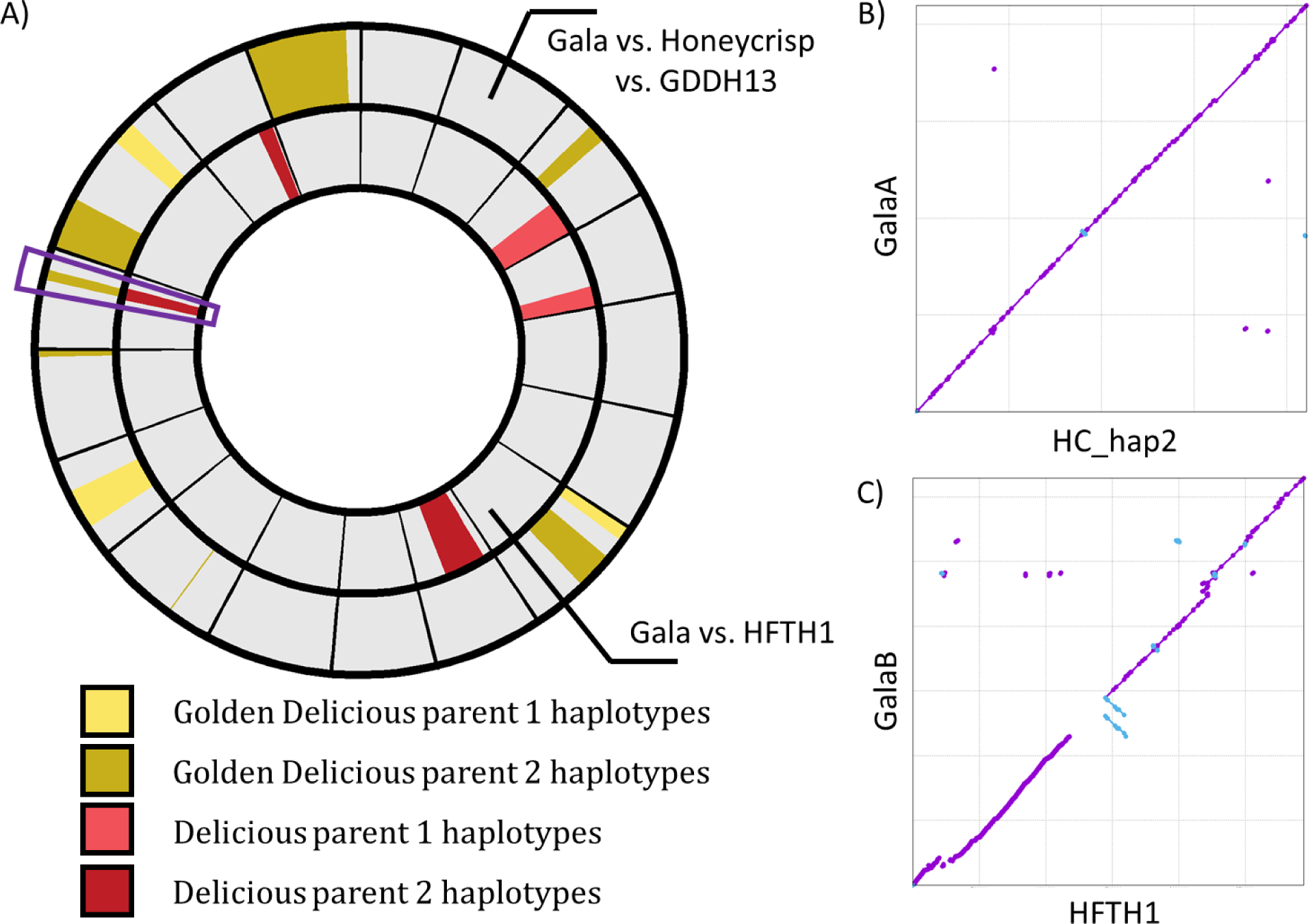
Shared segment analyses of available apple WGSs. A) The outer ring represents genomic segments shared by GDDH13, ‘Gala’, and ‘Honeycrisp’ through their common ancestor ‘Golden Delicious’ (yellow shades) and the inner ring represents genomic segments shared by HFTH1 and ‘Gala’ through their common ancestor ‘Delicious’ (red shades). The pair of segments outlined in purple (on chromosome 14) are elaborated in panels B and C. The determination of sharing is based on WGS-derived DNA profiles of GDDH13 and HFTH1 and SNP array DNA profiles of the other cultivars. B, C) Alignment of the shared segment on chromosome 14 for the Ga_A and HC_h2 haplomes (B) and the HFTH1 WGS and Ga_B haplomes (C). One region from HFTH1 appears to be missing from Ga_B while another region of HFTH1 is was detected three times of which two are in inverted orientation.

One segment on chromosome 14 was identified whose DNA profile was shared among all these WGSs (Fig. 3, Table S6). Zooming in at the sequence level revealed almost complete collinearity between HFTH1 and Ga_A. There was 99.87% to 99.94% sequence identity for the three pairwise comparisons. Alignments of HC_h2 and GDDH13 to Ga_A for this segment resulted in larger proportion of unaligned bases (2.9% and 6%, respectively) compared to alignments of HC_h2 and GDDH23 to each other (1.4% and 3.6% for HC_h2 and GDDH13, respectively). This difference indicated a part of this segment was absent in the Ga_A WGS but present in the GDDH13 and HC_h2 WGSs. Furthermore, a small part of this region was only present in the GDDH13 WGS, accounting for the higher proportion of unaligned bases in this WGS. An almost-homologous region shared between Ga_B and HFTH1 had 99.84% identity but large proportions of unaligned bases (12.9% in Ga_B and 23.7% in HFTH1) and a large region missing from Ga_B (Fig. 3C). The homologous regions on chromosome 14 within the Ga and HC WGS had a sequence identity of 98.32% and 99.95%, with 68%–74.4% and 1.13%– 1.17% unaligned bases, respectively. The two HC haplome sequences of a homozygous region in chromosome 7 of HC had 99.95% identity and 1.7–3.6% unaligned bases (Fig. S3). For this homozygous region, most of the sequence dissimilarity was observed at both edges of the segment.

## Discussion

We successfully demonstrated how WGSs can be evaluated, compared, and improved by using phased SNP array and pedigree data. We expanded how previously available high-quality genome-wide genotypic data can contribute to the evaluation of WGSs and the tools provided here enable similar analyses for new WGS assemblies in any diploid species. The ability to use previous information from highly curated and accurately ordered genome-wide genotypic data reduces redundancies, could improve cost-efficiency for creating new WGSs and could ensure these new WGSs are of the highest possible quality. The comparison of shared segments across WGSs could raise their data-quality across WGSs. For example, sequence reads across individuals could be compared for shared segments, thus increasing the coverage for these segments and improving the detection of sequencing errors. Overall, the combination and comparison of various genome-wide information is expected to improve the genetic resources available and lead to new insights in genetic phenomena like mutation and gene conversion.

The analyses shown here can complement bioinformatic parameters to evaluate the quality of genomes. For example, the results of this study highlight the quality of the HCv1 WGS regarding assembly and phasing but also indicate where improvements are possible, e.g. the composition of haplomes and their ordering according to their parental origin. The low rate of missing SNPs and inconsistencies with the genetic map support the author’s claim that the HCv1 assembly is of near completeness and highly contiguous (Khan et al. 2022). However, the top of chromosome 2 of haplome 1 did show a higher proportion of inconsistencies with the genetic map, suggesting that some improvement in the assembly is needed there. A more detailed analysis and comparison of results for that region in haplome 2 suggest that this region is missing from haplome 1 and not misassigned to another region of the haplome. SNP array-based segregation data from hundreds of Honeycrisp offspring also suggests the region is homozygous rather than having a ∼6.5cM/2Mb deletion. In addition, the informativeness of the WGSs could further be improved by ensuring that haplome 1 and haplome 2 only consisted of maternal and paternal chromosomes, respectively. Thus, even if a method has proven to provide adequate phasing, the methods developed in our study are still useful to ensure that chromosome assemblies are assigned to the correct haplomes.

Our approach can also provide more insights into the methods used for genome assembly. For example, the methods used for the assembly of the HCv1 WGS seem to be adequate at creating phased chromosome assemblies without the explicit use of additional parental sequences, marker or linkage map data information, or SNP array data. In contrast, the methods used for Ga WGS do not seem to result in well-phased assemblies as reported earlier (Vanderzande and Peace 2023). Combining the best reads and contigs per region when assembling the Ga_A haplome (Sun et al. 2020) is likely the reason for the poor chromosome-wide phasing as these best contigs are expected to represent either parental haplotype randomly. In addition, the exclusion of sequence reads once they had been assembled in the first haplome (Sun et al. 2020), would explain why larger number of SNPs do not map in the Ga_B WGS for homozygous regions (based on ‘Gala’s SNP marker calls and associated lack of segregation in its offspring). Furthermore, in the alignment by Sun et al (extended figure 5 of Sun et al. (2020)) regions of low identity in Ga_B largely coincide with the homozygous region of ‘Gala’ further indicating that regions could be missing in Ga_B for these homozygous regions. For these homozygous regions, the sequence reads are expected to be identical and were thus removed before they could be used for assembly of the Ga_B WGS. Furthermore, issues in homozygous regions are not limited to the Ga WGS as the proportion of SNPs that did not have a unique location was slightly higher for homozygous regions for most haplomes. This suggests that identical reads between homologues are not always assigned to both haplomes when assembling homozygous regions.

Furthermore, the increase of inconsistent positions for homozygous regions compared to the iGW map indicates the presence of assembly issues in homozygous regions. Thus, homozygous regions might need more attention when creating a diploid WGS and marker-based DNA profiles, such as those from SNP arrays, could identify these regions.

Future apple WGSs could benefit from comparing IBD-shared segments within and among WGSs upfront. For example, the very high number of unaligned bases between the two Ga haplomes for the chromosome 14 segment indicates a significant difference in their sequence quality, although some sequence dissimilarity would be expected as the two Ga segments originate from different ancestors. Next, a comparison of Ga segments with either the high-quality HFTH1 (Skytte af Sätra et al., in prep), GDDH13, or HC_hap1 WGS suggests that the Ga WGS may need more attention, especially the Ga_B WGS. Furthermore, the presence of a unique region in the shared segment of GDDH13 could indicate a missing part in the HC_hap1 WGS, incorrect assembly in the GDDH13 WGS, or a structural variation that occurred relatively recently. As expected, the sequence identity between the studied shared segments was high (99.89%) However, the combination of slightly lower sequence similarity and increased proportion of unaligned nucleotides compared to the studied homozygous region on chromosome 7 (99.95%) suggests that improvements in sequence similarity are possible between shared segments. In addition, high sequence similarity (99.97%) and coverage (>99.99%) observed between an Arabidopsis F1 hybrid WGS and the WGSs of its parents (Koren et al. 2018) would also suggest higher sequence similarities are possible between shared segments of parent and child. However, it can be expected that as the genetic distance between individuals increases, the similarity of their segments decreases. Furthermore, with clonally propagated crops, variations between clones can also accumulate over time. As only two segments were examined here, a more detailed comparison of shared segments is needed, both across the genome for the individuals used here, as well as across clones and more individuals of given genetic relationships.

Phased marker-based DNA profiles can identify assembly and resolve issues but the level by which depends on their marker density relative to the size of possibly misassembled samples. For instance, this study used 10K SNP and revealed more segments of maternal and paternal alleles per homolog for the Ga WGS than previously reported using 4K SNPs of a 8K SNP array (Vanderzande and Peace 2023). Nevertheless, even more switches between parental origin might be present in the Ga WGS as Sun et al. (Sun et al. 2020) reported an average of 59,428 and 47,357 contigs and scaffolds per haplome, respectively. If at least one SNP per contig is needed for proper phasing, roughly 60,000 heterozygous SNPs would be needed. In addition, checks for proper orienting would require at least two heterozygous SNP, thus further doubling the number of required SNP. Thus, SNP density needed would be dependent on contig number, distribution of the SNPs across these contigs, and on the level of heterozygosity of the genome. Data from the 480K SNP (e.g., from Muranty et al. (2018)) data might be sufficient, but would require new genotyping of ‘Gala’ and some direct relatives as well as subsequent data curation efforts. As this can be a laborious and time-consuming process, other methods such as trio-binning (Koren et al. 2018) might be better suited to improve the phasing of the Ga1 WGS as was recently demonstrated in the upcoming WA38 apple WGS (Zhang et al. 2024). These other methods would require the availability of low-depth sequence data on ‘Gala’s parents (‘Golden Delicious’ and ‘Kidd’s orange red’). Such data is available for Golden Delicious from the creation of a previous WGS (Velasco et al. 2010) but may not be available for Kidd’s orange red. Sequence data is available for the parents of Kidd’s orange red (Cox’s Orange Pippin and Delicious (Bianco et al. 2014)) but new methods would need to be created to use grandparental data in trio-binning.

Further improvements are needed for the routine comparison of the assembly of shared segments across WGSs. Unlike the comparison of SNP array profiles and WGS, the identification of shared segments across individuals with a common ancestor has not been automated yet. Such tool would facilitate the comparisons of these shared segments across WGSs. Such tool could be an upgrade from the already available the SPLoSH script (Howard et al. 2021). This script reports shared segments between two individuals but only when the shared segment is larger than the provided size and does not take into account known pedigree relationships. An upgrade to trace segments through the pedigree and ensure they are truly inherited from the shared ancestor and to consider segments of any size as long as their inheritance can be confirmed, would be needed. From there, the sequence of identified shared segments could be extracted from the WGSs and compared as was done here. Methods to create consensus sequence from shared segments are also needed. Some care is needed near the edges of the shared segments as they could only look the same based on SNP array data but actually not be shared and thus have a different sequence. This was observed in this study for the segment that was homozygous in Honeycrisp but for which the edges only looked homozygous based on SNP array data but showed differences in DNA sequences (Fig. S3). Further studies are needed to determine whether shared segments should be trimmed by a specific size to ensure DNA sequence identity. Although mutations could have occurred over generations, a consensus sequence could still be valuable. Furthermore, mutations can also occur between clones of same individual, especially those that have been around for decades yet only a single WGS is currently used. In future, variants against a consensus sequence could be used to explain differences between sports and could be used to determine lineage.

The combination of pedigree reconstruction, IBD-shared segment information, and consensus sequences for shared chromosome segments could enable cost-efficient strategies for the creation of new proxy-WGSs for pedigree-connected germplasm. For example, proxy WGSs could be obtained for founders or other key individuals like important breeding parents without the need to generate and assemble sequence data for each of them. Using the DNA profiles of connecting ancestors, the genome of other individuals within the germplasm could be described as mosaics built from the genome of these key individuals. This mosaic information could then be used to build any individual’s proxy-WGS from WGS information already available. However, the resulting proxy-WGS will have gaps for segments that contain a recombination and do not account for mutations, gene conversion, and local duplications and genome-rearrangements. For example, important breeding parents (as described in Peace et al. (Peace et al. 2014)) could be used to represent the breeding program and the availability of their WGS would enable the reconstruction of many other breeding individuals’ WGSs and may contribute to the stepwise building of WGSs of common founders. Similarly, Identity-by-State between individuals could also be used for the generation of proxy-WGSs without known pedigree connections as long shared segments are likely due to an unknown shared ancestry. Methods for IBS assessment such as the SPLosH method (Howard et al. 2021) can identify long shared segments based on DNA profiling tools. Yet, thresholds and edge-trimming would need to be established to reliably assign identical sequences to IBS regions. Recent efforts in pangenomes may result in the creation of new WGSs and other tools for efficient and informative WGS comparisons already provide sequence information for many apple individuals (Sun et al. 2020; Liao et al. 2023; Vollger et al. 2023; He et al. 2023; Wang et al. 2023; Jonkheer et al. 2024). Unfortunately, unique sequence segments were not always anchored to the core genome and therefore cannot be traced across generations to determine which other individuals carry them (Sun et al. 2020). Inclusion of crop-wild-relatives in a pan-genome, especially those that are important sources of biotic stress resistance could also be valuable, especially because some resistances are caused by the presence of additional disease resistance genes (Gao et al. 2019) that would be missed if only susceptible individuals were included. However, the presence of translocations and inversions between domesticated and crop-wild-relatives as observed in some crops (Breton et al. 2010; Wang et al. 2021; Ramsay et al. 2021) could complicate the assembly of hybrid WGSs.

The R script provided here could be a great tool for initial evaluation of unphased and phased genomes in the future, for apple and other species with available genome-wide DNA profiles. The quick runtime and availability of a wrapper function automating the analysis enable its easy integration in the creation of new WGSs. The script has already been used for the improvement of the recently published “Antonovka 172670-B” WGS (Švara et al. 2024)(Svara et al, in prep). Several R functions created can also be used to perform individual steps of the analysis that could be useful on their own, such as determining the physical position that corresponds best to the genetical position of a SNP. This is especially valuable when the best hit is not necessarily the one segregating in one’s germplasm. The breakdown of the analyses also enables users to look at intermediary results, and manually inspect any output and, if needed, adjust these output files before continuing the next steps by the same or other scripts. Accurate WGS positions of genetically mapped SNPs can be used for the integration of SNP array and (re)sequencing information, as anchor points to compare WGS sequences of a target region, and to create new genetic-physical maps for the WGS (Skytte af Sätra et al., in prep). Integration of SNP array and (re)sequencing information also ensures that previous efforts to characterize apple germplasm with SNP arrays (e.g., (Howard et al. 2018; Muranty et al. 2018; Vanderzande et al. 2019)) remain valuable as sequencing data becomes more prevalent. The scripts developed here could also provide further insights into the performance of SNPs across a wide set of WGSs. To enable such research, further work to incorporate the BLAST function as part of the software rather than requiring BLAST to be run separately would be valuable.

## Supplemental Files

Suppl Fig. S1 - Workflow to determine marker location in WGS based on genetic map

Suppl Fig. S2 - Workflow followed by wrapper R function

Suppl Fig. S3 – Alignment of both HC haplomes for homozygous region of chromosome 7

Suppl Table S1 - R functions with input and output.xlsx

Suppl Table S2 – Parameter file for wrapper function with explanation of parameters

Suppl Table S3 - Classification of multiple BLAST protocols against HFTH1 WGS based on results from the iGW

Suppl Table S4 - Classification of BLAST results and consistency with reference SNP array data for the studies WGSs

Suppl Table S5 - Parental Origin of WGS segments.xlsx

Suppl Table S6 - Shared WGS segments.xlsx

Suppl File S1 – R functions created for WGS evaluation.zip

Suppl File S2 - Demo Input Files.zip

Suppl File S3 - Demo Output Files.zip

## Supporting information

Suppl File S1 - R functions v1

Suppl File S2 - Demo Input Files

Suppl File S3 - Demo Output Files

Suppl Table S1 - R functions with input and output

Suppl Table S2 - Parameter file for wrapper function

Suppl Table S3 - Classification of multiple BLAST protocols against HFTH1

Suppl Table S4 - Classification of BLAST against various WGSs

Suppl Table S5 - Parental Origin of WGS segments

Suppl Table S6 - Shared WGS segments

## Acknowledgements

The authors thank Jonas Skytte af Sätra and Bjarne Larsen for early access to iGW information and Nick Howard for providing the most up-to-date phased reference 20K SNP array data of ‘Fuji’, ‘Gala’, ‘Golden Delicious’, and ‘Honeycrisp’. This work was partially funded by USDA’s National Institute of Food and Agriculture-Specialty Crop Research Initiative Project “RosBREED: Combining disease resistance and horticultural quality in new rosaceous cultivars” (2014-51181-22378) and the USDA National Institute of Food and Agriculture Hatch project 1014919, Crop Improvement and Sustainable Production Systems (WSU reference 00011).

