## Supplementary material for "Whole genome sequence improvement with pedigree information and reference genotypic profiles, demonstrated in outcrossing apple": Suppl. Figures 1-3

**Supplementary figures to Vanderzande et al. "Approaches for whole genome sequence improvement through the use of pedigree information and reference DNA profiles demonstrated in outcrossing apple"**

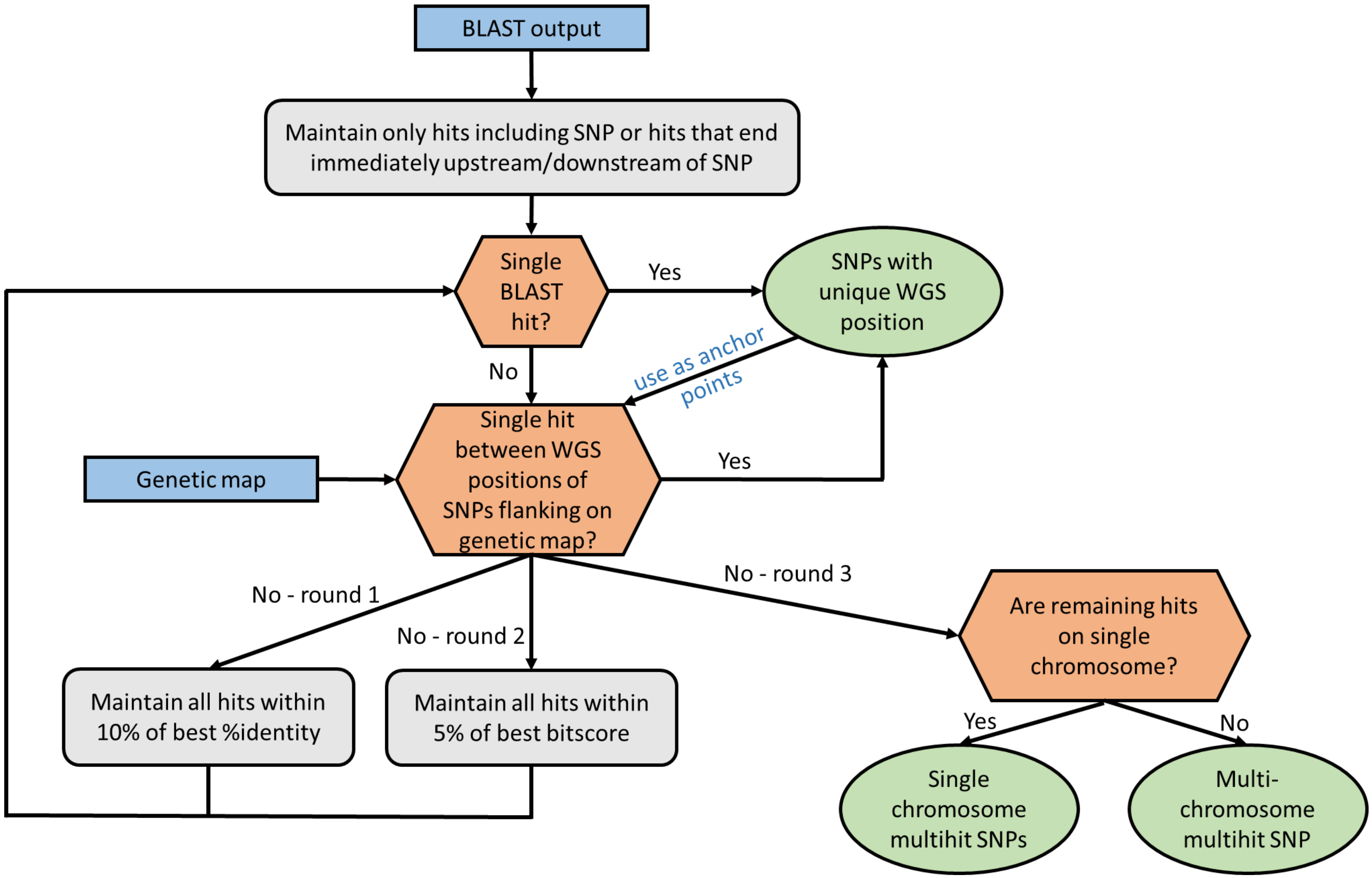


Supplementary Fig. S1: Workflow to determine the segregating locus in a whole genome sequence corresponding to SNPs of a genetic map. Blue boxes indicate input data used for the workflow while blue text indicates where generated output is used in subsequent steps of the workflow. Orange boxes indicate where SNPs are grouped into categories based on their corresponding BLAST hit results. Green boxes indicate outputs of the workflow.

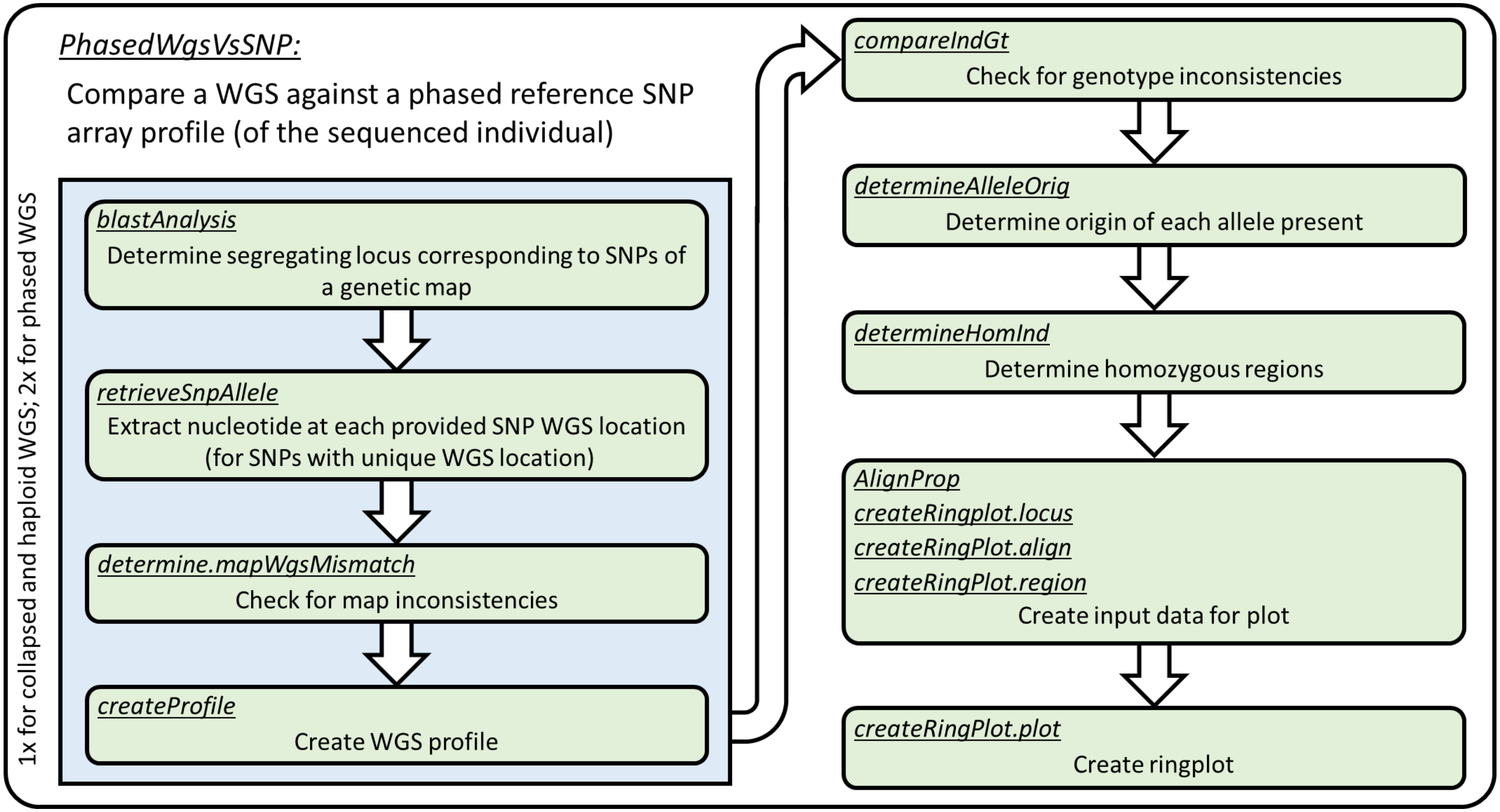


Supplementary Fig. S2: Workflow followed by the "PhasedWGSvsSNP" R function to compare reference phased SNP array DNA profiles to collapsed, haploid, and phased WGSs. For haploid and collapsed WGSs, the blue box series of steps is performed once whereas for phased diploid WGSs it is performed twice. Each green box represents a single step in the workflow, performed by one or multiple R functions. These R functions, indicated by underline and italics, provide input files for consequent steps and can also be run individually to investigate their output before continuing to the next step. For each r function, the required input files and resulting output files are listed in Supplementary Table SX.

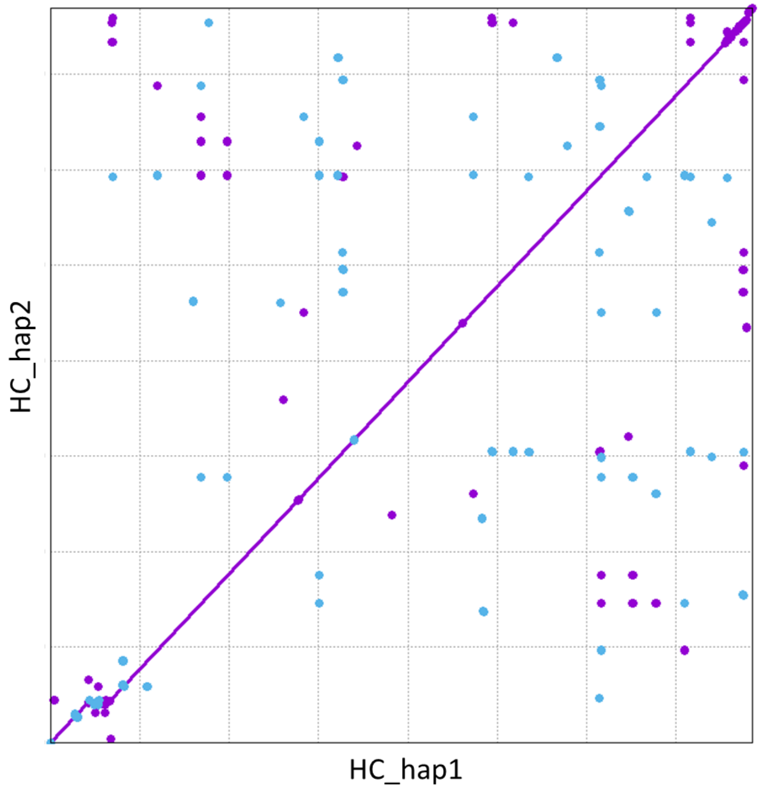


Supplementary Fig. S3: Alignment the two Honeycrisp WGS haplomes for the homozygous region of chromosome 7 as determined from 20K SNP array data. High similarity (99.95%) and low proportion of unaligned bases (1.7–3.6%) was observed except for the edges of the region. This suggest that most of this region is truly homozygous but that the size of the homozygous region is slightly overestimated when using the ‘Honeycrisp’ SNP array profile.
