## Supplementary material for "Whole genome sequence improvement with pedigree information and reference genotypic profiles, demonstrated in outcrossing apple": Suppl File S1 - R functions v1: R functions v1.R

```
###### functions created for WGS evaluation using reference genetic data
###### Developed as part of the following manuscript:
###### Vanderzande S, Peace C, van de Weg E. 
###### Approaches for whole genome sequence improvement through the use of pedigree information and reference DNA profiles demonstrated in outcrossing apple

###### Helper Functions ########

#' Determines cummulative position for each locus in a map file 
#' 
#' @param mapFile either filename if map needs to be loaded from external file or
#' name of R object. This R object is a matrix with loci as rows and at least 3 
#' columns: 1 column named "locusName" providing the name of the loci, 1 column 
#' providing linkage group info for each locus and 1 column providing positional
#' information within the linkage group
#' @param readMapFile logical; if T, an external map file with provided name 
#' will be loaded into R; if F, the provided R object will be used as map
#' @param lg string providing name of column with lg information
#' @param pos string providing name of column with positional info
#' @param chrSepSize numerical, distance in cM that should be added between between
#' two chromosomes, for example so that a line of the given size can be drawn 
#' between chromosomes for visualisation
#'
#' @return Matrix containing provided mapFile with cummulative positions added
#' @export
#'
#' @examples
addCummPos <- function(mapFile, readMapFile = F, lg = "lg", pos = "genPos", chrSepSize = 0){
  #if the mapfile needs to be loaded into R, load the correct file
  if(readMapFile){
    if(!file.exists(mapFile)){
      stop("'mapFile' file not found. Please make sure 'mapFile' file is correctly defined (case sensitive & including filename extenstion (e.g. '.csv')")
    }
    
    mapFile <- read.csv(file = mapFile, header = T, stringsAsFactors = F)
  }
  
  # check if the file includes the correct columns
  if (!all(c("locusName", lg, pos) %in% colnames(mapFile))){
    stop("Missing columns in 'mapFile'. Please make sure columns are named correctly and include 'locusName' and those defined as lg and pos parameters (case sensitive; default lg = 'lg' and pos = 'genPos')")
  }
  
  # determine what columns of the file hold the indicated lg and pos information
  lgCol <- which(colnames(mapFile) == lg)
  posCol <- which(colnames(mapFile) == pos)
  
  # for each lg, determine size (max position value) and cummulative start postion
  mapSumm <- data.frame(matrix(nrow = length(unique(mapFile[, lgCol])), ncol = 0))
  mapSumm$chrNames <- unique(mapFile[, lgCol])
  mapSumm$chrSize <- sapply(mapSumm$chrNames, function(x){return(max(mapFile[which(mapFile[,lgCol] == x), posCol]) + chrSepSize)})
  mapSumm$cummStart[1] <- 0
  for (i in 1:(nrow(mapSumm)-1)){
    mapSumm$cummStart[(i+1)] <- sum(mapSumm$chrSize[1:i])
  }
  
  #for each locus, determine cummulative start position of corresponding LG and calculate the locus' cummulative postion, then remove corresponding LG start location
  mapFile$cummStartLg <- mapSumm[match(mapFile[, lgCol], mapSumm$chrNames),]$cummStart
  mapFile$cummPos <- mapFile[ ,posCol] + mapFile$cummStartLg
  mapFile <- subset(mapFile, select=-c(cummStartLg))
  
  return(mapFile)
}

#' Function to convert a mapfile with locusName, lg, and genPos + any additional
#' info (i.e., cumm position, IBD info, ...) to a list where each LG is its own 
#' table
#' 
#' @param mapFile  either filename if map needs to be loaded from external file 
#' or name of R object. This R object is a matrix with loci as rows and at least
#' 2 #' columns: 1 column named "locusName" providing the name of the loci and 1
#' column providing linkage group info for each locus 
#' @param readMapFile logical; if T, an external map file with provided name 
#' will be loaded into R; if F, the provided R object will be used as map
#' @param lg string providing name of column with lg information
#' @return list, each element contains 1 linkage group from the map
#' @export
#'
#' @examples
mapFileToList <-function(mapFile, readMapFile = F, lg = "lg"){
  if(readMapFile){
    if(!file.exists(mapFile)){
      stop("'mapFile' file not found. Please make sure it is correctly defined (case sensitive & including filename extenstion (e.g. '.csv')")
    }
    
    mapFile <- data.frame(read.csv(file = mapFile, header = T), stringsAsFactors = F)
    
    # check if the file includes the correct columns
    if (!all(c("locusName", lg) %in% colnames(mapFile))){
      stop("Missing columns in 'mapFile'. Please make sure columns are named correctly and include 'locusName' and the one defined as lg (case sensitive; default lg = 'lg')")
    }
  }
  
  # determine what columns of the file hold the indicated lg information
  lgCol <- which(colnames(mapFile) == lg)
  
  lgNames <- unique(mapFile[,lgCol])
  mapList <- list()
  
  for (i in 1:length(lgNames)){
    mapList[[i]] <- mapFile[which(mapFile[,lgCol] == lgNames[i]),]
    names(mapList)[i] <- lgNames[i]
  }
  
  return(mapList)
}

#' Function that adds wgs chromosome and position to mapFile
#'
#' @param mapFile matrix providing genetic map where loci are rows and loci 
#' information in columns. Expects loci Names in column named "LocusName"
#' @param blastUniqHit matrix providing up to 1 whole genome location for loci of
#' the genetic map. WGS chromosome and position should be in columns named "wgsChr"
#' and "wgsPos", respectively 
#' @return list, each element contains 1 linkage group from the map 
wgsLocToMap <- function(mapFile, blastUniqHit){
  mapFile$wgsChr <- blastUniqHit[match(mapFile$locusName, blastUniqHit$locusName),]$wgsChr
  mapFile$wgsPos <- blastUniqHit[match(mapFile$locusName, blastUniqHit$locusName),]$wgsPos
  return(mapFile)
}

#' Function that returns WGS position for 3 closest loci (according to genetic 
#' map) upstream and downstream with single blast hit on, when provided, target 
#' chromosome.
#'
#' @param locusName string, gives name of locus to find flanking loci for
#' @param targetChr NULL or string providing a chromosome name. If not NULL, only
#' loci with unique BLAST hits on the provided WGS chromosome are considered
#' @param nLoci NULL or numerical, number of upstream/downstream loci to return.
#' If NULL, all loci within `dist.cM` will be returned
#' @param mapLG matrix providing genetic map, should contain columns named 
#' "locusName", "genPos", and "wgsChr" providing locus names, position in genetic
#' map, and the chromosome of the WGS to which the unique BLAST hit mapped
#' @param dist.cM distance in cM within which flankingloci with unique BLAST hit
#' should be looked for on the genetic map
#' 
#' @return List with two elements: upstream and downstream loci. Each element is
#' either a matrix of 1 to 3 rows with the corresponding loci or is NULL if no 
#' loci were found
flankingSNPs <- function(locusName, targetChr = NULL, nloci = NULL, mapLG, dist.cM){
  
  rws.locus <- which(mapLG$locusName == locusName)
  genPos.locus <- mapLG$genPos[rws.locus]
  
  upstrLoci <- mapLG[0:(rws.locus-1), ]
  upstrLoci <- upstrLoci[!is.na(upstrLoci$wgsChr), ]
  upstrLoci <- upstrLoci[upstrLoci$genPos >= (genPos.locus - dist.cM), ]
  dwnstrLoci <- mapLG[(rws.locus+1):(nrow(mapLG)+1), ]
  dwnstrLoci <- dwnstrLoci[!is.na(dwnstrLoci$wgsChr), ]
  dwnstrLoci <- dwnstrLoci[dwnstrLoci$genPos <= (genPos.locus + dist.cM), ]
  
  if (!is.null(targetChr)){
    upstrLoci <- upstrLoci[which(upstrLoci$wgsChr %in% targetChr), ]
    dwnstrLoci <- dwnstrLoci[which(dwnstrLoci$wgsChr %in% targetChr), ]
  }
  

  if (nrow(upstrLoci) > 0 & !is.null(nloci)){
    upstrLoci <- upstrLoci[max(1,(nrow(upstrLoci)-nloci+1)):nrow(upstrLoci), ]
  } else if (nrow(upstrLoci) == 0){
    upstrLoci <- NULL
  }
    
  if (nrow(dwnstrLoci) > 0 & !is.null(nloci)){
    dwnstrLoci <- dwnstrLoci[1:min(nloci,nrow(dwnstrLoci)), ]
  } else if (nrow(dwnstrLoci) == 0){
    dwnstrLoci <- NULL
  }
    
  return(list(upstrLoci,dwnstrLoci))
} 

#' Function that returns BLAST hits for a locus that are located within a window
#' on a provided target chromosome defined by the provided results from flanking
#' loci whose unique hit also mapped to the target chromosome 
#'
#' @param blastHits.locus matrix containing BLAST hit of 1 locus for which most
#' likely hit is to be determined. BLAST outformat 6 is expected with "wgsPos" 
#' added as column, providing the corresponding location of the SNP in the WGS 
#' for each BLAST hit. 
#' @param targetChr string providing the chromosome name on which BLAST hit should
#' have mapped
#' @param flankingSNPs List with two elements: upstream and downstream loci. Each
#' element is either a matrix of 1 to 3 rows with the corresponding loci or is
#' NULL if no loci were found. This list can be created with the `flankingSNPs()`
#' function
#' @param singleSide logical, indicates whether only loci on a single side (either
#' upstream or downstream) should be considered. if TRUE, a BLAST hit is still 
#' searched even if flanking loci with a single hit are only present upstream or
#' downstream
#' @param dist.bp numerical, indicating the distance in bp that can be searched
#' outside the window defined by flanking SNPs for a matching BLAST hit
#' 
#' @return matrix of BLAST hits for a locus that are located on the target chromosome
#' within a window defined by flanking markers. If no BLAST hits are found, NULL
#' is returned instead
matchingHitFlSnpSet <- function(blastHits.locus, targetChr, flankingSNPs, singleSide = F, dist.bp = 0){
  if( is.null(flankingSNPs[[1]]) & is.null(flankingSNPs[[2]]) ){
    return (NULL)
    
  } else if ( (is.null(flankingSNPs[[1]]) | is.null(flankingSNPs[[2]])) & !singleSide ){
    return (NULL)
    
  } else {
    blastHits.locus.chr <- blastHits.locus[blastHits.locus$wgsChr %in% targetChr, ]
    
    if(is.null(flankingSNPs[[1]])){
      blastHits.locus.chr.pos <- blastHits.locus.chr[which(blastHits.locus.chr$wgsPos < (max(flankingSNPs[[2]]$wgsPos) + dist.bp)), ]
      
    } else if (is.null(flankingSNPs[[2]])){
      blastHits.locus.chr.pos <- blastHits.locus.chr[which(blastHits.locus.chr$wgsPos > (min(flankingSNPs[[1]]$wgsPos) - dist.bp)), ]
      
    } else {
      blastHits.locus.chr.pos <- blastHits.locus.chr[which(blastHits.locus.chr$wgsPos > (min(flankingSNPs[[1]]$wgsPos, flankingSNPs[[2]]$wgsPos) - dist.bp)), ]
      blastHits.locus.chr.pos <- blastHits.locus.chr.pos[which(blastHits.locus.chr.pos$wgsPos < (max(flankingSNPs[[1]]$wgsPos, flankingSNPs[[2]]$wgsPos) + dist.bp)), ]
    }
    
    if (nrow(blastHits.locus.chr.pos) > 1) {
      if(is.null(flankingSNPs[[1]])){
        blastHits.locus.chr.pos2 <- blastHits.locus.chr[which(blastHits.locus.chr$wgsPos < flankingSNPs[[2]]$wgsPos[1]), ]
      } else if (is.null(flankingSNPs[[2]])){
        blastHits.locus.chr.pos2 <- blastHits.locus.chr[which(blastHits.locus.chr$wgsPos > flankingSNPs[[1]]$wgsPos[length(flankingSNPs[[1]]$wgsPos)]), ]
      } else {
        blastHits.locus.chr.pos2 <- blastHits.locus.chr[which(blastHits.locus.chr$wgsPos > min(flankingSNPs[[2]]$wgsPos[1], flankingSNPs[[1]]$wgsPos[length(flankingSNPs[[1]]$wgsPos)])), ]
        blastHits.locus.chr.pos2 <- blastHits.locus.chr.pos2[which(blastHits.locus.chr.pos2$wgsPos < max(flankingSNPs[[2]]$wgsPos[1], flankingSNPs[[1]]$wgsPos[length(flankingSNPs[[1]]$wgsPos)])), ]
      }
      if (nrow(blastHits.locus.chr.pos2) > 0) {
        blastHits.locus.chr.pos <- blastHits.locus.chr.pos2
      }
    }
    
    if (nrow(blastHits.locus.chr.pos) > 0) {
      return(blastHits.locus.chr.pos)
    } else {
      return (NULL)
    }
  }  
} 

#' Function that checks flanking loci with a unique BLAST hit and checks how their
#' BLAST hits relate, then depending on the situation, defines one or multiple 
#' WGS regions to search for BLAST hits using the `matchingHitFlSnpSet()` function
#' and returns the most likely BLAST hit for the target locus
#'
#' @param locusName string, name of the target locus
#' @param blastHits matrix containing all BLAST hits.BLAST outformat 6 is expected
#' with "wgsPos" added as column, providing the corresponding location of the SNP
#' in the WGS for each BLAST hit. 
#' @param mapList List, with each element of the list corresponding to the genetic
#' map of a single chromosome
#' @param chr.prefix string, prefix in chromosome name of WGS to the chromosome
#' name according to the genetic map. For example if the chromosome in the map is
#' called "1" and in the WGS the chromosome is called "chr1A", the prefix is "chr"
#' @param chr.suffix string, suffix in chromosome name of WGS to the chromosome
#' name according to the genetic map. For example if the chromosome in the map is
#' called "1" and in the WGS the chromosome is called "chr1A", the prefix is "A"
#' @param singleSide logical, indicates whether only loci on a single side (either
#' upstream or downstream) should be considered. if TRUE, a BLAST hit is still 
#' searched even if flanking loci with a single hit are only present upstream or
#' downstream
#' @param dist.bp numerical, indicating the distance in bp that can be searched
#' outside the window defined by flanking SNPs for a matching BLAST hit
#' @param dist.cM numerical, distance in cM within which flankingloci with unique
#' BLAST hit should be looked for on the genetic map
#' @param nloci NULL or numerical, number of upstream/downstream loci to consider
#' when determining the best BLAST hit. If NULL, all loci within `dist.cM` will
#' be returned
#' 
#' @return matrix of BLAST hits for a locus that are located within a window 
#' defined by flanking markers. If no BLAST hits are found, NULL is returned instead
mapMatchingHit <- function(locusName, blastHits, mapList, chr.prefix, chr.suffix, singleSide = F, dist.bp = 0, dist.cM = 2.5, nloci = NULL){
  
  blastHits.locus <- blastHits[blastHits$locusName == locusName, ]
  
  mapLG.locus <- names(which(sapply(mapList, function(y) locusName %in% y$locusName)))
  mapLGListNb.locus <- which(names(mapList) == mapLG.locus)
  
  if (length(mapLG.locus) == 0){
    return(NULL)
    
  } else if (length(mapLG.locus) == 1){
    
    Chr.names <- c(paste0(chr.prefix, mapLG.locus, chr.suffix), paste0(chr.prefix, "0", mapLG.locus, chr.suffix))
    flLoci <- flankingSNPs(locusName = locusName, mapLG = mapList[[mapLGListNb.locus]], dist.cM = dist.cM, nloci = 1)
    
    upstrChr <- flLoci[[1]]$wgsChr
    downstrChr <- flLoci[[2]]$wgsChr
    
    if(!is.null(upstrChr) & !is.null(downstrChr)){
      
      if (upstrChr == downstrChr){
        flLoci.trgt <- flankingSNPs(locusName = locusName, targetChr = upstrChr, mapLG = mapList[[mapLGListNb.locus]], dist.cM = dist.cM, nloci = nloci)
        targetHit <- matchingHitFlSnpSet(blastHits.locus = blastHits.locus,
                                         targetChr = upstrChr, 
                                         flankingSNPs = flLoci.trgt,
                                         singleSide = singleSide,
                                         dist.bp = dist.bp)
        
        if (is.null(targetHit) & !(upstrChr %in% Chr.names)){
          flLoci.trgt <- flankingSNPs(locusName = locusName, targetChr = Chr.names, mapLG = mapList[[mapLGListNb.locus]], dist.cM = dist.cM, nloci = nloci)
          targetHit <- matchingHitFlSnpSet(blastHits.locus = blastHits.locus,
                                           targetChr = Chr.names, 
                                           flankingSNPs = flLoci.trgt,
                                           singleSide = singleSide,
                                           dist.bp = dist.bp)  
        }
        
      } else if ((upstrChr %in% Chr.names) | (downstrChr %in% Chr.names)){
        if ((upstrChr %in% Chr.names)){altChr <- downstrChr} else {altChr <- upstrChr}
        
        flLoci.trgt <- flankingSNPs(locusName = locusName, targetChr = Chr.names, mapLG = mapList[[mapLGListNb.locus]], dist.cM = dist.cM, nloci = nloci)
        targetHit <- matchingHitFlSnpSet(blastHits.locus = blastHits.locus,
                                         targetChr = Chr.names, 
                                         flankingSNPs = flLoci.trgt,
                                         singleSide = singleSide,
                                         dist.bp = dist.bp)
        
        if (is.null(targetHit)){
          flLoci.trgt <- flankingSNPs(locusName = locusName, targetChr = altChr, mapLG = mapList[[mapLGListNb.locus]], dist.cM = dist.cM, nloci = nloci)
          targetHit <- matchingHitFlSnpSet(blastHits.locus = blastHits.locus,
                                           targetChr = altChr, 
                                           flankingSNPs = flLoci.trgt,
                                           singleSide = singleSide,
                                           dist.bp = dist.bp)
        }
        
      } else {
        flLoci.trgt1 <- flankingSNPs(locusName = locusName, targetChr = upstrChr, mapLG = mapList[[mapLGListNb.locus]], dist.cM = dist.cM, nloci = nloci)
        targetHit1 <- matchingHitFlSnpSet(blastHits.locus = blastHits.locus,
                                          targetChr = upstrChr, 
                                          flankingSNPs = flLoci.trgt1,
                                          singleSide = singleSide,
                                          dist.bp = dist.bp)
        
        flLoci.trgt2 <- flankingSNPs(locusName = locusName, targetChr = downstrChr, mapLG = mapList[[mapLGListNb.locus]], dist.cM = dist.cM, nloci = nloci)
        targetHit2 <- matchingHitFlSnpSet(blastHits.locus = blastHits.locus,
                                          targetChr = downstrChr, 
                                          flankingSNPs = flLoci.trgt2,
                                          singleSide = singleSide,
                                          dist.bp = dist.bp)
        
        targetHit <- rbind(targetHit1, targetHit2)
        
        if (is.null(targetHit)){
          flLoci.trgt <- flankingSNPs(locusName = locusName, targetChr = Chr.names, mapLG = mapList[[mapLGListNb.locus]], dist.cM = dist.cM, nloci = nloci)
          targetHit <- matchingHitFlSnpSet(blastHits.locus = blastHits.locus,
                                           targetChr = Chr.names, 
                                           flankingSNPs = flLoci.trgt,
                                           singleSide = singleSide,
                                           dist.bp = dist.bp)
          
        }
      }
      
    } else if ((!is.null(upstrChr) | !is.null(downstrChr)) & singleSide){
      
      flChr <- c(upstrChr, downstrChr)
      flLoci.trgt <- flankingSNPs(locusName = locusName, targetChr = flChr, mapLG = mapList[[mapLGListNb.locus]], dist.cM = dist.cM, nloci = nloci)
      targetHit <- matchingHitFlSnpSet(blastHits.locus = blastHits.locus,
                                       targetChr = flChr, 
                                       flankingSNPs = flLoci.trgt,
                                       singleSide = singleSide,
                                       dist.bp = dist.bp)
      
      if (is.null(targetHit) & !(flChr %in% Chr.names)){
        flLoci.trgt <- flankingSNPs(locusName = locusName, targetChr = Chr.names, mapLG = mapList[[mapLGListNb.locus]], dist.cM = dist.cM, nloci = nloci)
        targetHit <- matchingHitFlSnpSet(blastHits.locus = blastHits.locus,
                                         targetChr = Chr.names, 
                                         flankingSNPs = flLoci.trgt,
                                         singleSide = singleSide,
                                         dist.bp = dist.bp)
      }
      
    } else {
      targetHit <- NULL
    }
    
    return (targetHit)
    
  } else {
    
    stop("marker(s) found in multiple regions of linkage map. Provide unique genetic positions per marker")
  }
} 


#' Function that takes x% highest results of specific criterion/trait for specific
#' individual/locus/group
#' 
#' @param groupName string, name of individual/locus/group/... for which best
#' values need to be determined
#' @param groupColName string, column name in `datTable` that provides groupName
#' per observation
#' @param datTable data frame that holds the data
#' @param critName string, column name that holds trait/criterion for which the 
#' best x% needs to be chosen (expects positive numeric values)
#' @param critLvl numerical, minimum proportion of maximum value that needs to be
#' kept. E.g., 0.95 means all values that are at least 95% of maximum are kept 
#' (1 means only best, 0 means all results)
#' 
#' @return dataframe, subset of datTable that contains the x% best results
chooseBestbyCrit <- function(groupName, groupColName, datTable, critName, critLvl = 1){
  if(critLvl < 0 | critLvl > 1){
    warning("critlvl expected between 0 and 1 (inclusive) ")
  }
  
  groupCol <- which(colnames(datTable) == groupColName)
  critCol <- which(colnames(datTable) == critName)
  datTable.grp <- datTable[which(datTable[,groupCol] == groupName), ]
  datTable.grp.crit <- datTable.grp[which(datTable.grp[,critCol] >= critLvl*max(datTable.grp[,critCol])), ]
  return(datTable.grp.crit)
}

#' Function to convert a Fasta file to tabular format
#' Function adjusted from GitHub: https://github.com/lrjoshi/FastaTabular
#'
#' @param fastaFile external file name or R object name following the fasta format
#' @param readFasta logical, if TRUE, an external file is loaded into R with the
#' provided name, otherwise, an R object with the provided name is used
#'
#' @return matrix with two columms, first contains the sequence names provided in
#' the fasta file, the second column contains the corresponding sequences
#' @export
#'
#' @examples
fastaToTabular <- function(fastaFile, readFasta = T){
  #read fasta file
  if(readFasta){
    fastaFile <- readLines(fastaFile)
  }
  
  #find the genename location by grepping >
  location <- which((stringr::str_sub(fastaFile,1,1))==">")
  
  #start an empty vector to collect name and sequence 
  name=c()
  sequence =c()
  
  #number of genes= number of loops
  #extract name first
  for ( i in 1:length(location)){
    name_line = location[i]
    name1 = fastaFile[name_line]
    name=c(name,name1)
    #extract sequence between the names
    #the last sequence will be missed using this strategy 
    #so, we are using if condition to extract last sequence 
    start= location[i]+1
    end = location[i+1]-1
    if ( i < length (location)){
      end=end
    } else {
      end=length(fastaFile)
    }
    
    lines = start:end
    sequence1= as.character(paste(fastaFile[lines],collapse = ""))
    sequence =c(sequence,sequence1)
  }
  
  #now create table using name and sequence vector 
  data <- tibble::tibble(name,sequence)
  return(data)
}

#' Function to retrieve the sequence from a whole genome sequence given a specific 
#' chromosome, start location and end location
#'
#' @param genome matrix containing 2 columns, 1 called "name" containing the 
#' chromosome (or sequence) names, the other containing the corresponding sequences
#' @param querChr string, indicating the name of the chromosome of which a sequence
#' should be returned
#' @param querStart numerical, start position on the chromosome of the sequence
#' to be returned
#' @param querEnd numerical, last position on the chromosome of the sequence to
#' be returned
#'
#' @return string, sequence of the provided chromosome from querStart to querEnd
#' @export
#'
#' @examples
retrieveSeq <- function(genome, querChr, querStart, querEnd){
  return(substr(genome[genome$name == querChr, 2], querStart, querEnd))
}


#' Function to read mhaplotypes file created by FlexQLT which can contain multiple
#' headers not needed for the analyses here. this file contains the phased reference
#' data 
#'
#' @param mhaplotypes string, file name of the file to be read
#'
#' @return matrix, mhaplotype file only containing the needed data and correct
#' column names ("ID" for the individuals name, "VAR" indicating row type, and
#' loci names for all phased loci)
#' @export
#'
#' @examples
read.mhaplotypes <- function(mhaplotypes = "mhaplotypes.csv"){
  mhap <- read.csv(mhaplotypes, 
                   header = F, stringsAsFactors = F, check.names = F,
                   colClasses = "character", comment.char = ";")
  
  fline <- which(mhap[,1]=="ID")
  mhap.colnames <- mhap[fline, ]
  mhap <- data.frame(mhap[(fline+1):nrow(mhap), ], stringsAsFactors = F)
  colnames(mhap) <- mhap.colnames
  
  return(mhap)
}

#' function to determine the SNP position in the target sequence for a BLAST hit
#' based on the location of the SNP within provided query sequence
#'
#' @param qStart numerical, start position in query sequence that aligned to the
#' target sequence
#' @param qEnd numerical, end position in query sequence that aligned to the
#' target sequence
#' @param snpFlankSeqPos numerical, position in the query sequence where the SNP
#' is located
#' @param gStart numerical, position on the target sequence where alignment of 
#' the query starts
#' @param gEnd numerical, position on the target sequence where alignment of 
#' the query ends
#'
#' @return position of the SNP in the target sequence
#' @export
#'
#' @examples
determine.snpWgsPos <- function(qStart, qEnd, snpFlankSeqPos, gStart, gEnd){
  # the following parts avoid gaps in the alignment in specific cases
  if(snpFlankSeqPos == qStart-1){
    if(gStart < gEnd){
      snpPos <- gStart - 1
    } else if(gEnd < gStart){
      snpPos <- gStart + 1
    }  
  } else if (snpFlankSeqPos == qStart){
    snpPos <- gStart
  } else if (snpFlankSeqPos == qEnd){
    snpPos <- gEnd
  } else if (snpFlankSeqPos == qEnd + 1){
    if(gStart < gEnd){
      snpPos <- gEnd+1
    } else if(gEnd < gStart){
      snpPos <- gEnd-1
    }
  # From here, the location determination can be incorrect in case of gaps in the alignment  
  } else {
    if(gStart < gEnd){
      snpPos <- gStart + (snpFlankSeqPos - qStart)
    } else if(gEnd < gStart){
      snpPos <- gStart - (snpFlankSeqPos - qStart)
    }}
  return(snpPos)
}

#' function to find a corresponding hit in the BLAST using short sequences (to
#' avoid gaps) that matched the chosen locus from the BLAST using full sequences
#'
#' @param locusName string, name of the locus for which the right short BLAST hit
#' needs to be chosed
#' @param wgsChr string, name of the chromosome to which the chosen full BLAST
#' hit aligned
#' @param wgsPos numerical, WGS position of the locus based on the full BLAST
#' @param gStart numerical, start position of target sequence where the chosen
#' full BLAST hit aligned
#' @param gEnd numerical, end position of target sequence where the chosen full
#' BLAST hit aligned
#' @param shortBlast matrix containing the results of the BLAST using short
#' sequences. BLAST outformat 6 is expected with an added column wgsPos providing
#' the corresponding position of the SNP within the aligned region
#' 
#' @return matrix, single BLAST hit that best matches the chosen hit from the
#' full BLAST
#' @export
#'
#' @examples
determine.corrShortBlast <- function(locusName, wgsChr, wgsPos, gStart, gEnd, shortBlast){
  shortBlast.locus <- shortBlast[which(shortBlast$locusName == locusName & 
                                         shortBlast$wgsChr == wgsChr &
                                         shortBlast$gStart >= min(gStart, gEnd) &
                                         shortBlast$gStart <= max(gStart, gEnd) &
                                         shortBlast$gEnd >= min(gStart, gEnd) &
                                         shortBlast$gEnd <= max(gStart, gEnd)), ]
  if (nrow(shortBlast.locus) > 0){
    target <- min(abs(shortBlast.locus$wgsPos - wgsPos))
    shortBlast.locus <- shortBlast.locus[which(abs(shortBlast.locus$wgsPos - wgsPos) == target), ]
  }
  
  if (nrow(shortBlast.locus) == 1){
    return(shortBlast.locus)
  } else {
    return(NULL)
  }
}


###### Main Workflow Functions ########

#' Function to determine best hit per locus, based on a genetic map if available
#'
#' @param blastResults.full external file name or R matrix containing BLASTn 
#' generated output using full length flanking sequences or probes (use '-outfmt 6'
#' command to generate correct output)
#' @param readBlast.full logical; indicates whether blastResults.full needs to be
#' read from an external file. If true, blastResults.full is filename with file 
#' extension, if false, blastResults.full is R object that holds the BLAST results
#' @param blastResults.short optional parameter; `NULL`, external file name or R
#' matrix containing BLASTn generated output using shorter flanking sequences,
#' used to avoid gaps in alignments and determine SNPs position more accurately;
#' not needed for probes where the SNP is first/last nucleotide or directly 
#' upstream or downstream of probe sequence (use '-outfmt 6'  command to generate
#' correct output)
#' @param readBlast.short logical; ignored if `blastResults.short` is NULL; 
#' indicates if blastResults.short needs to be read from an external file. If true,
#' blastResults.short is filename with file extension, if false, blastResults.short
#' is R object that holds the BLAST results
#' @param chr.prefix string, prefix in chromosome name of WGS to the chromosome
#' name according to the genetic map. For example if the chromosome in the map is
#' called "1" and in the WGS the chromosome is called "chr1A", the prefix is "chr"
#' @param chr.suffix string, suffix in chromosome name of WGS to the chromosome
#' name according to the genetic map. For example if the chromosome in the map is
#' called "1" and in the WGS the chromosome is called "chr1A", the prefix is "A"
#' @param snpInfo external csv file name or R matrix containing information on SNP
#' location within BLASTed sequences. Contains at least 2 columns and has a third
#' when `blastResults.short` is not NULL: 1. Column called "snpName" that has names
#' of the SNPs/sequences that were BLASTed; 2. calumns called "flSeqSnpPos.full"
#' containing the (numerical) position of each SNP within the sequences used for 
#' `blastResults.full`; 3. column called "flSeqSnpPos.short" containing the 
#' (numerical) position of each SNP within the sequences used for `blastResults.short`.
#' Positions can be outside of BLASTed sequence (e.g., in case of probes)
#' @param readSnpInfo logical; readSnpInfo indicates if snpInfo needs to be read
#' from an external file. If true, snpInfo is filename with file extension, if 
#' false, snpInfo is R object that holds the needed snpInfo
#' @param mapFile optional; `NULL`, external file name or R object containing the
#' genetic map for (some of) the BLASTed loci, must contain at least 3 columns:
#' 1. column called "locusName" which has locus names as used for the BLAST; 
#' 2. column called "lg" which has the corresponding linkage group (or chromosome)
#' of each locus; 3. column called "genPos" which has the corresponding genetic 
#' position (in cM) of each locus in the map
#' @param readMapFile logical; ignored if `mapFile` is NULL; indicates whether
#' mapFile needs to be read from an external file. If true, mapFile is filename
#' with file extension, if false, mapFile is R object that holds the needed snpInfo
#' @param lenght.hit numerical, indicating minimum length of a full BLAST hit to
#' be considered for WGS position determination
#' @param dist.bp numerical, indicating the distance in bp that can be searched
#' outside the window defined by flanking SNPs for a matching BLAST hit
#' @param dist.cm numerical, distance in cM within which flanking loci with unique
#' BLAST hit should be looked for on the genetic map
#' @param nLoci NULL or numerical, number of upstream/downstream loci to consider
#' when determining the best BLAST hit. If NULL, all loci within `dist.cM` will
#' be returned
#' @param writeOutput logical, indicates whether results need to be written to 
#' external files
#' @param outfolder `NULL` or string, if not NULL, name of the folder in which
#' results should be saved. Will be created if it doesn't exist 
#' @param outpath string, prefix that output files will have
#' @param returnOutput logical, indicates whether results should be returned (i.e.,
#' results could also be only written to external files)
#'
#' @return List of three matrices containing 1. SNPs with unique hit in WGS and
#' their position; 2. SNPs with non-unique hits within single chromosome and their
#' position; 3. SNPs with non-unique hits across multiple chromosomes
#' @export
#'
#' @examples
blastAnalysis <- function(blastResults.full = "snpblast_full.out",
                          readBlast.full = T,
                          blastResults.short = NULL,
                          readBlast.short = T,
                          chr.prefix,
                          chr.suffix,
                          snpInfo = "snpinfo.csv",
                          readSnpInfo = T,
                          mapFile = NULL,
                          readMapFile = T,
                          length.hit = 15,
                          dist.bp = 0,
                          dist.cM = 2.5,
                          nloci = NULL,
                          writeOutput = T,
                          outfolder = NULL,
                          outpath = "snpLoc_refGenome",
                          returnOutput = T) {
  
  # Read BLAST output and add headers
  print("Step 1: Reading Data")
  
  if(readBlast.full){
    if(!file.exists(blastResults.full)){
      stop("'blastResults.full' file not found. Please make sure 'blastResults.full' file is correctly defined (case sensitive & including filename extenstion (e.g. '.out' or '.csv')")
    }
    blastResults.full <- data.frame(read.table(file = blastResults.full), stringsAsFactors = F)
  }
  
  if(ncol(blastResults.full) != 12){
    stop("Unexpected Number of Columns in Provided 'blastOuput' file. Please check it was generated using the '-outfmt 6' command and contains the following columns 'querry name', 'subject name', 'proportion identitiy', 'lenght of Match', 'number mismatches', 'number gaps', 'querry Start', 'querry End', 'subject Start', 'subject gEnd', 'eValue', and 'bitScore'")
  }
  
  colnames(blastResults.full) <- c("locusName", "wgsChr", "pIdent","lenMatch", "mismatch", "gap", "qStart", "qEnd", "gStart", "gEnd", "eVal", "bitScore")
  
  if (!is.null(blastResults.short)){
    if(readBlast.short){
      if(!file.exists(blastResults.short)){
        stop("'blastResults.short' file not found. Please make sure 'blastResults.short' file is correctly defined (case sensitive & including filename extenstion (e.g. '.out' or '.csv')")
      }
    
      blastResults.short <- data.frame(read.table(file = blastResults.short), stringsAsFactors = F)
    }
  
    if(ncol(blastResults.short) != 12){
      stop("Unexpected Number of Columns in Provided 'blastOuput' file. Please check it was generated using the '-outfmt 6' command and contains the following columns 'querry name', 'subject name', 'proportion identitiy', 'lenght of Match', 'number mismatches', 'number gaps', 'querry Start', 'querry End', 'subject Start', 'subject gEnd', 'eValue', and 'bitScore'")
    } 
  
    colnames(blastResults.short) <- c("locusName", "wgsChr", "pIdent","lenMatch", "mismatch", "gap", "qStart", "qEnd", "gStart", "gEnd", "eVal", "bitScore")
  }
  
  if(readSnpInfo){
    if(!file.exists(snpInfo)){
      stop("'snpInfo' file not found. Please make sure 'snpInfo' file is correctly defined (case sensitive & including filename extenstion (e.g. '.out' or '.csv')")
    }
    # Read SNP info file which provides SNP location within
    snpInfo <-  data.frame(read.csv(file = snpInfo, header = T), stringsAsFactors = F)
  }
  
  if(is.null(blastResults.short)){
    if(!all(c("snpName", "flSeqSnpPos.full") %in% colnames(snpInfo))){
      stop("Missing columns in 'snpInfo'. Please make sure columns are named correctly and include 'snpName', 'flSeqSnpPos.full' (case sensitive)")
    }
  } else {
    if(!all(c("snpName", "flSeqSnpPos.full","flSeqSnpPos.short") %in% colnames(snpInfo))){
      stop("Missing columns in 'snpInfo'. Please make sure columns are named correctly and include 'snpName', 'flSeqSnpPos.full', 'flSeqSnpPos.short' (case sensitive)")
    }
  }
  
  if (!is.null(mapFile)){
    
    if(readMapFile){
      if(!file.exists(mapFile)){
        stop("'mapFile' file not found. Please make sure 'mapFile' file is correctly defined (case sensitive & including filename extenstion (e.g. '.out' or '.csv')")
      }
      
      mapFile <- data.frame(read.csv(file = mapFile, header = T), stringsAsFactors = F)
    }
    
    if(!all(c("locusName", "lg", "genPos") %in% colnames(mapFile))){
      stop("Missing columns in 'mapFile'. Please make sure columns are named correctly and include 'locusName', 'lg', and 'genPos' (case sensitive) ")
    }
  }
    
  # Add SNP's position in flanking sequence to full BLAST results.
  blastResults.full$flSeqSnpPos <- snpInfo[match(blastResults.full$locusName, snpInfo$snpName),]$flSeqSnpPos.full
  
  #add position of the SNP in the genome
  blastResults.full$wgsPos <- mapply(determine.snpWgsPos,
                                     qStart = blastResults.full$qStart,
                                     qEnd = blastResults.full$qEnd,
                                     snpFlankSeqPos = blastResults.full$flSeqSnpPos, 
                                     gStart = blastResults.full$gStart,
                                     gEnd = blastResults.full$gEnd)
  
  # if blast hit ends/starts at SNP location within sequence or just before/after that location, position can always be accurately be determined. Therefore, gaps have no influence on position determination and are set to 0
  blastResults.full$gap[which(blastResults.full$flSeqSnpPos == (blastResults.full$qStart - 1) | 
                                blastResults.full$flSeqSnpPos == blastResults.full$qStart |
                                blastResults.full$flSeqSnpPos == blastResults.full$qEnd |
                                blastResults.full$flSeqSnpPos == (blastResults.full$qEnd + 1))] <- 0
  
  # Remove any results where the matched seq does not cover right up to the SNP + miminum 15bp long
  blastResults.full <- subset(blastResults.full, qStart <= (flSeqSnpPos + 1), drop = F)
  blastResults.full <- subset(blastResults.full, qEnd >= (flSeqSnpPos - 1), drop = F)
  blastResults.full <- subset(blastResults.full, lenMatch >= length.hit, drop = F)
  
  print("Step 2: Taking Unique Hits")
  
  # Count for each locus/SNP how many hits were found
  blastResults.full$nHits <- dplyr::add_count(blastResults.full, locusName)$n
  
  # Find all SNPs with just one hit and save them
  blastResults.chosenHit <- blastResults.full[blastResults.full$nHits == 1, ,drop = F]
  blastResults.remainHit <- blastResults.full[blastResults.full$nHits > 1, ,drop = F]
  
  # If a map is available, use info to check if a single hit matches the map
  if (!is.null(mapFile)){
    print("Step 3: Checking Multi-Hit SNPs for Hits Consistent with Genetic Map")
    
    mapList <- mapFileToList(mapFile)
    mapList <- lapply(mapList, wgsLocToMap, blastUniqHit = blastResults.chosenHit)
    
    mapList.matchingWgs <- pbapply::pblapply(unique(blastResults.remainHit$locusName), 
                                              mapMatchingHit, 
                                              blastHits = blastResults.remainHit, 
                                              mapList = mapList, 
                                              chr.prefix = chr.prefix, 
                                              chr.suffix = chr.suffix,
                                              singleSide = F,
                                              dist.bp = dist.bp,
                                              dist.cM = dist.cM,
                                              nloci = nloci)
    
    mapList.matchingWgs <- mapList.matchingWgs[!sapply(mapList.matchingWgs,is.null)]
    foundhits <- sapply(mapList.matchingWgs, function(x){return(unique(x$locusName))})
    
    blastResults.chosenHit <- rbind(blastResults.chosenHit, do.call(rbind, mapList.matchingWgs[sapply(mapList.matchingWgs, function(x){nrow(x) == 1})]))
    blastResults.remainHit <- rbind(blastResults.remainHit[!(blastResults.remainHit$locusName %in% foundhits), ], do.call(rbind, mapList.matchingWgs[sapply(mapList.matchingWgs, function(x){nrow(x) > 1})]))
    
  } else {
    print("Step 3 Skipped - No Genetic Map")
  }
  
  print("Step 4: Keep Hits within 10% of Highest %Identity per Locus")
  
  blastResults.remainHit <- pbapply::pblapply(unique(blastResults.remainHit$locusName), chooseBestbyCrit, groupColName = "locusName", datTable = blastResults.remainHit, critName = "pIdent", critLvl = 0.9)
  blastResults.remainHit <- do.call(rbind, blastResults.remainHit)
  blastResults.remainHit$nHits <- dplyr::add_count(blastResults.remainHit, locusName)$n
  
  blastResults.chosenHit <- rbind(blastResults.chosenHit, blastResults.remainHit[blastResults.remainHit$nHits == 1, ,drop = F])
  blastResults.remainHit <- blastResults.remainHit[blastResults.remainHit$nHits > 1, ,drop = F]
  
  if (!is.null(mapFile)){
    print("Step 5: Checking Remaining Multi-Hit SNPs for Hits Consistent with Genetic Map")
    
    mapList <- lapply(mapList, wgsLocToMap, blastUniqHit = blastResults.chosenHit)
    
    mapList.matchingWgs <- pbapply::pblapply(unique(blastResults.remainHit$locusName), 
                                              mapMatchingHit, 
                                              blastHits = blastResults.remainHit, 
                                              mapList = mapList, 
                                              chr.prefix = chr.prefix, 
                                              chr.suffix = chr.suffix,
                                              singleSide = F,
                                              dist.bp = dist.bp,
                                              dist.cM = dist.cM,
                                              nloci = nloci)
    
    mapList.matchingWgs <- mapList.matchingWgs[!sapply(mapList.matchingWgs,is.null)]
    foundhits <- sapply(mapList.matchingWgs, function(x){return(unique(x$locusName))})
    
    blastResults.chosenHit <- rbind(blastResults.chosenHit, do.call(rbind, mapList.matchingWgs[sapply(mapList.matchingWgs, function(x){nrow(x) == 1})]))
    blastResults.remainHit <- rbind(blastResults.remainHit[!(blastResults.remainHit$locusName %in% foundhits), ], do.call(rbind, mapList.matchingWgs[sapply(mapList.matchingWgs, function(x){nrow(x) > 1})]))
    
  } else {
    print("Step 5 Skipped - No Genetic Map")
  }
  
  print("Step 6: Keep Hits within 5% of Highest Bitscore per Locus")
  
  blastResults.remainHit <- pbapply::pblapply(unique(blastResults.remainHit$locusName), chooseBestbyCrit, groupColName = "locusName", datTable = blastResults.remainHit, critName = "bitScore", critLvl = 0.95)
  blastResults.remainHit <- do.call(rbind, blastResults.remainHit)
  blastResults.remainHit$nHits <- dplyr::add_count(blastResults.remainHit, locusName)$n
  
  blastResults.chosenHit <- rbind(blastResults.chosenHit, blastResults.remainHit[blastResults.remainHit$nHits == 1, ,drop = F])
  blastResults.remainHit <- blastResults.remainHit[blastResults.remainHit$nHits > 1, ,drop = F]
  
  if (!is.null(mapFile)){
    print("Step 7: Checking Remaining Multi-Hit SNPs for Hits Consistent with Genetic Map")
    
    mapList.matchingWgs <- pbapply::pblapply(unique(blastResults.remainHit$locusName), 
                                              mapMatchingHit, 
                                              blastHits = blastResults.remainHit, 
                                              mapList = mapList, 
                                              chr.prefix = chr.prefix, 
                                              chr.suffix = chr.suffix,
                                              singleSide = T,
                                              dist.bp = dist.bp,
                                              dist.cM = dist.cM,
                                              nloci = nloci)
    
    mapList.matchingWgs <- mapList.matchingWgs[!sapply(mapList.matchingWgs,is.null)]
    foundhits <- sapply(mapList.matchingWgs, function(x){return(unique(x$locusName))})
    
    blastResults.chosenHit <- rbind(blastResults.chosenHit, do.call(rbind, mapList.matchingWgs[sapply(mapList.matchingWgs, function(x){nrow(x) == 1})]))
    blastResults.remainHit <- rbind(blastResults.remainHit[!(blastResults.remainHit$locusName %in% foundhits), ], do.call(rbind, mapList.matchingWgs[sapply(mapList.matchingWgs, function(x){nrow(x) > 1})]))
    
  } else {
    print("Step 7 Skipped - No Genetic Map")
  }
  
  if (!is.null(blastResults.short)){
    
    blastResults.all <- rbind(blastResults.chosenHit, blastResults.remainHit)
    
    if (nrow(blastResults.all[blastResults.all$gap > 0, ,drop = F]) == 0){
      
      print("Steps 8-9 Skipped - No adjustments needed for gaps in alignment") 
      
    } else {
      
      print("Step 8: Load Results of Short Fragment BLAST")
      #For all SNPs with single hit that had a gap in their alignment (=> SNP position probably wrong), look at the BLAST results using only few bp flanking seq and find the hit that (almost) matches the location of the unique hit with gap
      
      blastResults.short$flSeqSnpPos <- snpInfo[match(blastResults.short$locusName, snpInfo$snpName),]$flSeqSnpPos.short
      blastResults.short <- subset(blastResults.short, qStart <= (flSeqSnpPos + 1), drop = F)
      blastResults.short <- subset(blastResults.short, qEnd >= (flSeqSnpPos - 1), drop = F)
      
      #add position of the SNP in the genome
      blastResults.short$wgsPos <- mapply(determine.snpWgsPos,
                                          qStart = blastResults.short$qStart,
                                          qEnd = blastResults.short$qEnd,
                                          snpFlankSeqPos = blastResults.short$flSeqSnpPos,
                                          gStart = blastResults.short$gStart,
                                          gEnd = blastResults.short$gEnd)
      
      blastResults.short$gap[which(blastResults.short$flSeqSnpPos == (blastResults.short$qStart - 1) | 
                                     blastResults.short$flSeqSnpPos == blastResults.short$qStart |
                                     blastResults.short$flSeqSnpPos == blastResults.short$qEnd |
                                     blastResults.short$flSeqSnpPos == (blastResults.short$qEnd + 1)  )] <- 0
      
      
      print("Step 9: Updating SNPs with Gaps in Alignment")
      #Filter unique SNPs for those with(out) gap
      blastResults.all.gap <- blastResults.all[blastResults.all$gap > 0, ,drop = F]
      blastResults.all.noGap <- blastResults.all[blastResults.all$gap == 0, ,drop = F]
      
      blastResults.gap.short <- pbapply::pbmapply(determine.corrShortBlast,
                                                   locusName = blastResults.all.gap$locusName, 
                                                   wgsChr = blastResults.all.gap$wgsChr,
                                                   wgsPos = blastResults.all.gap$wgsPos,
                                                   gStart = blastResults.all.gap$gStart,
                                                   gEnd = blastResults.all.gap$gEnd,
                                                   MoreArgs = list(shortBlast = blastResults.short))
      
      blastResults.all.gapFixed <- t(sapply(1:nrow(blastResults.all.gap),function(x){if(!is.null(blastResults.gap.short[[x]])){return(blastResults.gap.short[[x]])}else{return(blastResults.all.gap[x,1:14])}}))
      blastResults.all.gapFixed <- lapply(1:nrow(blastResults.all.gap),function(x){if(!is.null(blastResults.gap.short[[x]])){return(blastResults.gap.short[[x]])}else{return(blastResults.all.gap[x,1:14])}})
      blastResults.all.gapFixed <- do.call(rbind, blastResults.all.gapFixed)
      blastResults.all.gapFixed <- rbind(blastResults.all.noGap[,1:14], blastResults.all.gapFixed)
      blastResults.all <- blastResults.all.gapFixed[order(factor(blastResults.all.gapFixed$locusName, levels = unique(blastResults.all$locusName))),]
      
      blastResults.all$nHits <- dplyr::add_count(blastResults.all, locusName)$n
      blastResults.chosenHit <- blastResults.all[which(blastResults.all$nHits == 1), ]
      blastResults.remainHit <- blastResults.all[which(blastResults.all$nHits > 1), ]
    }
    
  } else {
    print("Steps 8-9 Skipped - No Short Segment BLAST Results Provided")
    
  }
  
  print("Step 10: Writing Results into Files")
  
  if(nrow(blastResults.remainHit) > 0){
    multiHitSingChr <- data.frame()
    multiHitMultiChr <- data.frame()
    
    for (i in 1:length(unique(blastResults.remainHit$locusName))){
      locusResults <- blastResults.remainHit[blastResults.remainHit$locusName == unique(blastResults.remainHit$locusName)[i], ]
      
      if (length(unique(locusResults$wgsChr)) == 1){
        multiHitSingChr <- rbind(multiHitSingChr,locusResults)
        
      } else {
        multiHitMultiChr <- rbind(multiHitMultiChr, locusResults)
        
      }
    }
  } else {
    multiHitSingChr <- data.frame(matrix(NA, nrow = 0, ncol = ncol(blastResults.remainHit)))
    multiHitMultiChr <- data.frame(matrix(NA, nrow = 0, ncol = ncol(blastResults.remainHit)))
    colnames(multiHitSingChr) <- colnames(blastResults.remainHit)
    colnames(multiHitSingChr) <- colnames(blastResults.remainHit)
    
  }
  
  if (writeOutput){
    if (!is.null(outfolder)){
      if (!file.exists(outfolder)){
        dir.create(outfolder)
      }
      outpath <- paste0(outfolder,"/",outpath)
    } 
      
    write.csv(blastResults.chosenHit, file = paste0(outpath, "_UniqueHits.csv"), row.names = F)
    write.csv(multiHitSingChr, file = paste0(outpath, "_MulitHits_SingleChr.csv"), row.names = F)
    write.csv(multiHitMultiChr, file = paste0(outpath, "_MulitHits_MultiChr.csv"), row.names = F)
  }
  
  if(returnOutput){
    return(list(uniqueHits = blastResults.chosenHit, 
                singleChrHits = multiHitSingChr,
                multiChrHits = multiHitMultiChr))
  }
}


#' Returns SNP allele in the given WGS for a set of SNPs with provided WGSlocation
#'
#' @param wgs external file name or R object name following the fasta format
#' @param readWgs logical, if TRUE, an external file with the provided name is
#' loaded as wgs file into R , otherwise, an R object with the provided name is used
#' @param snpPos external file name or R object containing at least three columns;
#' 1. column named "locusName" with the SNP names for which the allele should be
#' retrieved; 2. column named "wgsChr" which holds the chromosome/sequence name
#' as used in the WGS file on which the SNPs are located; 3. columns named "wgsPos"
#' which holds the positions within the chromosome/sequence where the SNPs are
#' located. If `skipgap` is TRUE, a fourth column called 'gap' is required which
#' indicates how many gaps were present in the alignment (i.e. the exact position
#' of the SNP is not certain). Output from `blastAnalysis` can be used
#' @param readSnpPos logical, if TRUE, an external file with the #' provided name
#' is loaded into R as snpPos file, otherwise, an R object with the provided name
#' is used
#' @param skipGap logical, if TRUE, any SNPs with gaps in their alignment will be
#' skipped and their assigned allele will be set to missing ("-")
#' @param convertABfmt logical, if TRUE, alleles will be converted to the AB format
#' used by SNP arrays (i.e., "A" and "T" are coded as "A" and "C" and "G" are
#' coded as "B")
#' @param writeOutput logical, indicates whether results need to be written to 
#' external files
#' @param outfolder `NULL` or string, if not NULL, name of the folder in which
#' results should be saved. Will be created if it doesn't exist 
#' @param outpath string, prefix that output files will have
#' @param returnOutput logical, indicates whether results should be returned (i.e.,
#' results could also be only written to external files)
#'
#' @return matrix of two columns containing the names of the SNPs and their
#' respecitve WGS alleles (based on the provided WGS positions)
#' @export
#'
#' @examples
retrieveSnpAllele <- function(wgs = "refWGS.fasta",
                              readWgs = T,
                              snpPos = "snpPos.csv",
                              readSnpPos = F,
                              skipGap = T,
                              convertABfmt = T,
                              writeOutput = T,
                              outfolder = NULL,
                              outpath = "snpAllele_refGenome",
                              returnOutput = T) {
  
  if(readWgs){
    if(!file.exists(wgs)){
      stop("'wgs' file not found. Please make sure 'wgs' file is correctly defined (case sensitive & including filename extenstion (e.g. '.fasta')")
    }
  }
  
  if(readSnpPos){
    if(!file.exists(snpPos)){
      stop("'snpPos' file not found. Please make sure 'snpPos' file is correctly defined (case sensitive & including filename extenstion (e.g. '.csv')")
    }
    
    snpPos <- read.csv(file = snpPos, header = T, stringsAsFactors = F)
  }
  
  if(!all(c("locusName", "wgsChr", "wgsPos") %in% colnames(snpPos))){
    stop("Missing columns in 'snpPos' file. Please make sure columns are named correctly and include 'locusName', 'wgsChr', and 'wgsPos' (case sensitive) ")
  }
  
  if(skipGap){
    if(!("gap" %in% colnames(snpPos))){
      stop("Gap info missing from 'snpPos' file. Please make sure column with gap info called 'gap' is present (case sensitive) or skipGap is set to 'F'")
    }
    snpPos <- snpPos[snpPos$gap == 0, , drop=F]
  }
  
  print("Step 1: Loading WGS - this can take a while")
  
  refGenome <- fastaToTabular(wgs, readFasta = readWgs)
  refGenome <- as.data.frame(refGenome, stringsAsFactors = F)
  ## remove ">" from sequence names
  refGenome[,1] <- gsub(">","",refGenome[,1])
  
  print("Step 2: Retrieving SNP Alleles")
  
  snpPos <- subset(snpPos, select=c(locusName, wgsChr, wgsPos))
  
  snpPos$wgsAllele <- mapply(retrieveSeq, querChr = snpPos$wgsChr, querStart = snpPos$wgsPos, querEnd = snpPos$wgsPos, MoreArgs = list(genome = refGenome))
  
  if (convertABfmt){
    snpPos$wgsAllele <- sapply(snpPos$wgsAllele, convertNtToSNPar <- function(nt){
      if (nt %in% c("a","A","t","T", "w", "W")){
        return("A")
      } else if (nt %in% c("c","C","g","G", "s", "S")){
        return("B")
      } else {
        return("-")
      }})
  }
  
  if(writeOutput){
    if (!is.null(outfolder)){
      if (!file.exists(outfolder)){
        dir.create(outfolder)
      }
      outpath <- paste0(outfolder,"/",outpath)
    } 
    
    write.csv(snpPos, file = paste0(outpath, "_wgsAlleles.csv"), row.names = F)
  }
  
  if (returnOutput){
    return(snpPos)
  }
}


#' Function that compares physical positions in a WGS to the genetic map and reports
#' any inconsistencies between them
#'
#' @param mapWgsFile external file name or R object name that contains the
#' positions of both WGS and genetic map. Must contain at least 5 columns: 1. column
#' named "locusName" containing the names of the loci, 2. column named "wgsChr"
#' containing the assigned chromosome in the WGS of each locus, 3. column named
#' "wgsPos" containing the assigned posiion in the wGS (in bp) of each locus,
#' 4. column named "lg" containing the linkage group in the genetic map of each
#' locus, 5. column named "genPos" containing the genetic position in the genetic
#' map of each locus
#' @param readMapWGS logical, if TRUE, an external file with the provided name is
#' loaded as `mapWGSFile` into R , otherwise, an R object with the provided name
#' is used
#' @param chr.prefix string, prefix in chromosome name of WGS to the chromosome
#' name according to the genetic map. For example if the chromosome in the map is
#' called "1" and in the WGS the chromosome is called "chr1A", the prefix is "chr"
#' @param chr.suffix string, suffix in chromosome name of WGS to the chromosome
#' name according to the genetic map. For example if the chromosome in the map is
#' called "1" and in the WGS the chromosome is called "chr1A", the prefix is "A"
#' @param dist.bp numerical, indicating the distance in bp that can be searched
#' outside the window defined by flanking SNPs for a matching BLAST hit
#' @param dist.cm numerical, distance in cM within which flankingloci with unique
#' BLAST hit should be looked for on the genetic map
#' @param nloci NULL or numerical, number of upstream/downstream loci to consider
#' when checking consistency with genetic map. If NULL, all loci within `dist.cM`
#' will be returned
#' @param writeOutput logical, indicates whether results need to be written to 
#' external files
#' @param outfolder `NULL` or string, if not NULL, name of the folder in which
#' results should be saved. Will be created if it doesn't exist 
#' @param outpath string, prefix that output files will have
#' @param returnOutput logical, indicates whether results should be returned (i.e.,
#' results could also be only written to external files)
#'
#' @return matrix containing a list of SNP names that are inconsistent with the
#' genetic map and their chromosome and bp position in the WGS
#' @export
#'
#' @examples
determine.mapWgsMismatch <- function(mapWgsFile = "mapWgsFile.csv",
                                     readMapWGS = T,
                                     chr.prefix,
                                     chr.suffix,
                                     dist.bp = 0,
                                     dist.cM = 2.5,
                                     nloci = NULL,
                                     writeOutput = T,
                                     outfolder = NULL,
                                     outpath = "refGenome",
                                     returnOutput = F){
  if(readMapWGS){
    if(!file.exists(mapWgsFile)){
      stop("'wgsAlleles' file not found. Please make sure 'wgsAlleles' file is correctly defined (case sensitive & including filename extenstion (e.g. '.csv')")
    }
    mapWgsFile <- read.csv(file = mapWgsFile, header = T, stringsAsFactors = F)
  }
  
  if (!all(c("locusName", "wgsChr", "wgsPos", "lg", "genPos") %in% colnames(mapWgsFile))){
    stop("Missing columns in 'mapWgsFile' file. Please make sure columns are named correctly and include 'locusName', 'wgsChr', 'wgsPos', 'lg', and 'genPos' (case sensitive)")
  }
  
  mismatchLg <- mapWgsFile[mapply(function(x, y){!(x %in% c(paste0(chr.prefix, y, chr.suffix),
                                                            paste0(chr.prefix, "0", y, chr.suffix))) & !is.na(x)},
                                  x = mapWgsFile$wgsChr, y = mapWgsFile$lg), ]
  
  mapWgsFile[which(mapWgsFile$locusName %in% mismatchLg$locusName), ]$wgsChr <- NA
  mapWgsFile.noNA <- mapWgsFile[which(!is.na(mapWgsFile$wgsChr)), ]
  mapWgsFile.noNA$posMismatch <- pbapply::pbmapply(function(locusName, wgsChr, wgsPos, dist.bp, mapFile){
    flSnps <- flankingSNPs(locusName = locusName, targetChr = wgsChr, mapLG = mapFile, dist.cM = dist.cM, nloci = nloci)
    if (!is.null(flSnps[[1]]) & !is.null(flSnps[[2]])){
      if (wgsPos > (min(flSnps[[1]]$wgsPos, flSnps[[2]]$wgsPos) - dist.bp) &
          wgsPos < (max(flSnps[[1]]$wgsPos, flSnps[[2]]$wgsPos) + dist.bp)){
        return(F)
      } else {
        return(T)
      }
    } else if (!is.null(flSnps[[1]])){
      if (wgsPos > (min(flSnps[[1]]$wgsPos) - dist.bp)){
        return(F)
      } else {
        return(T)
      }  
    } else if (!is.null(flSnps[[2]])){
      if (wgsPos < (max(flSnps[[2]]$wgsPos) + dist.bp)){
        return(F)
      } else {
        return(T)
      }
    } else {
      return(F)
    } 
  },
  locusName = mapWgsFile.noNA$locusName,
  wgsChr = mapWgsFile.noNA$wgsChr,
  wgsPos = mapWgsFile.noNA$wgsPos,
  MoreArgs = list(dist.bp = dist.bp, mapFile = mapWgsFile.noNA))
  mismatchPos <- mapWgsFile.noNA[which(mapWgsFile.noNA$posMismatch), !colnames(mapWgsFile.noNA) == "posMismatch"]
  
  if(writeOutput){
    if (!is.null(outfolder)){
      if (!file.exists(outfolder)){
        dir.create(outfolder)
      }
      outpath <- paste0(outfolder,"/",outpath)
    } 
    
    
    write.csv(rbind(mismatchLg, mismatchPos), file = paste0(outpath, "_mapMismatch.csv"), row.names = F)
  }
  
  if (returnOutput){
    return(rbind(mismatchLg, mismatchPos))
  }
}


#' Creates a DNA profiles from the WGS alleles based on the genetic map, possibly
#' settinh alleles to missing for markers whose location is not consistent with
#' the genetic map.
#'
#' @param wgsAlleles external file name or R object name. Matrix of two columns
#' 1. column named "locusName" containing the names of the loci and 2. column
#' named "wgsAllele" containing the respective WGS alleles. Output from `retrieveSnpAllele`
#' can be used.
#' @param mapFile either filename if map needs to be loaded from external file or
#' name of R object. Matrix with loci as rows and at least 3 
#' columns: 1 column named "locusName" providing the name of the loci, 1 column 
#' providing linkage group info for each locus and 1 column providing positional
#' information within the linkage group
#' @param mapMismatch NULL, external file name, or R object name. if not NULL,
#' matrix with at least 1 column named "locusName" containing the names of loci
#' that are inconsistent with the genetic map. Output from `determine.mapWgsMismatch`
#' can be used.
#' @param readAlleles logical; if T, an external file with provided name will be
#' loaded into R as `wgsAlleles` matrix; if F, the provided R object will be used
#' as `wgsAlleles` matrix
#' @param readMapFile logical; if T, an external map file with provided name 
#' will be loaded into R; if F, the provided R object will be used as map
#' @param readMismatch logical; if T, an external file with provided name will be
#' loaded into R as `mapMismatch` matrix; if F, the provided R object will be used
#' as `mapMismatch` matrix
#' @param naString string used for missing alleles, by default "-" is used
#' @param writeOutput logical, indicates whether results need to be written to 
#' external files
#' @param outfolder `NULL` or string, if not NULL, name of the folder in which
#' results should be saved. Will be created if it doesn't exist 
#' @param outpath string, prefix that output files will have
#' @param returnOutput logical, indicates whether results should be returned (i.e.,
#' results could also be only written to external files)
#'
#' @return matrix containing a WGS DNA profile based on the genetic map
#' @export 
#'
#' @examples
createProfile <- function(wgsAlleles = "wgsAlleles.csv",
                          mapFile = "mapFile.csv",
                          mapMismatch = NULL,
                          readAlleles = F,
                          readMapFile = F,
                          readMismatch = F,
                          naString = "-",
                          writeOutput = T,
                          outfolder = NULL,
                          outpath = "snpProf_refGenome",
                          returnOutput = F) {
  
  if(readAlleles){
    if(!file.exists(wgsAlleles)){
      stop("'wgsAlleles' file not found. Please make sure 'wgsAlleles' file is correctly defined (case sensitive & including filename extenstion (e.g. '.csv')")
    }
    
    wgsAlleles <- read.csv(file = wgsAlleles, header = T, stringsAsFactors = F)
  }  
  
  if (!all(c("locusName", "wgsAllele") %in% colnames(wgsAlleles))){
    stop("Missing columns in 'wgsAllelesFile' file. Please make sure columns are named correctly and include 'locusName', and 'wgsAllele' (case sensitive)")
  }
  
  if(readMapFile){
    if(!file.exists(mapFile)){
      stop("'mapFile' file not found. Please make sure 'mapFile' file is correctly defined (case sensitive & including filename extenstion (e.g. '.csv')")
    }
    
    mapFile <- read.csv(file = mapFile, header = T, stringsAsFactors = F)
  }  
  
  if (!all(c("locusName", "lg", "genPos") %in% colnames(mapFile))){
    stop("Missing columns in 'mapFile'. Please make sure columns are named correctly and include 'locusName', 'lg', and 'genPos' (case sensitive)")
  }
  
  if(!is.null(mapMismatch)){
    if(readMismatch){
      if(!file.exists(MapMismatch)){
        stop("'mapMismatch' file not found. Please make sure 'mapMismatch' file is correctly defined (case sensitive & including filename extenstion (e.g. '.csv')")
      }
      
      mapMismatch <- read.csv(file = mapMismatch, header = T, stringsAsFactors = F)
    }  
    
    if (!("locusName" %in% colnames(mapMismatch))){
      stop("Missing column in 'mapMismatch'. Please make sure columns are named correctly and include 'locusName' (case sensitive)")
    }
  }
  
  mapFile$wgsAllele <- wgsAlleles[match(mapFile$locusName, wgsAlleles$locusName),]$wgsAllele
  
  if(!is.null(mapMismatch)){
    mapFile[which(mapFile$locusName %in% mapMismatch$locusName), ]$wgsAllele <- NA
  }
  
  profile <- data.frame(t(mapFile$wgsAllele), stringsAsFactors = F)
  colnames(profile) <- mapFile$locusName
  
  if(writeOutput){
    profile[is.na(profile)] <- naString
    
    if (!is.null(outfolder)){
      if (!file.exists(outfolder)){
        dir.create(outfolder)
      }
      outpath <- paste0(outfolder,"/",outpath)
    } 
    
    write.csv(profile, file = paste0(outpath, "_dnaProfile.csv"), row.names = F)
  }
  
  if (returnOutput){
    profile[profile == naString] <- NA
    return(profile)
  }
}

#' Compares haplotypes or genotypes between two individuals
#'
#' @param ind1.hap1 vector of strings representing the alleles of the first
#' haplotype of individual 1 or `NULL`. If `NULL`, a vector is expected under
#' ind1.gt. Expected to be the same size and order of loci as ind1.hap2/ind2.hap1/
#' ind2.hap2/ind2.gt
#' @param ind1.hap2 optional vector of strings representing the alleles of the
#' second haplotype of individual 1 (when not haploid)  or `NULL`. Expected to be
#' the same size and order of loci as ind1.hap1/ind2.hap1/ind2.hap2/ind2.gt
#' @param ind2.hap1 vector of strings representing the alleles of the first
#' haplotype of individual 2 or `NULL`. If `NULL`, a vector is expected under
#' ind2.gt. Expected to be the same size and order of loci as ind1.hap1/ind1.hap2/
#' ind1.gt/ind2.hap2
#' @param ind2.hap2 optional vector of strings representing the alleles of the
#' second haplotype of individual 2 (when not haploid)  or `NULL`. Expected to be
#' the same size and order of loci as ind1.hap1/ind1.hap2/ind1.gt/ind2.hap1
#' @param ind1.gt vector of strings representing the genotypes of individual 1 or
#' `NULL`. If `NULL`, a vector is expected under ind1.hap1. Both alleles of each
#' genotype should be separated by the string used for parameter `allelSep`. 
#' Expected to be the same size and order of loci as ind2.hap1/ind2.hap2/ind2.gt
#' @param ind2.gt vector of strings representing the genotypes of individual 2 or
#' `NULL`. If `NULL`, a vector is expected under ind1.hap1. Both alleles of each
#' genotype should be separated by the string used for parameter `allelSep`.
#' Expected to be the same size and order of loci as ind1.hap1/ind1.hap2/ind1.gt
#' @param locusNames vector containing the names of each locus for which alleles
#' or genotypes are provided under ind1.hap1/ind1.hap2/ind2.hap1/ind2.hap2/ind1.gt/
#' ind2.gt
#' @param allelSep string used as separator between two alleles in the genotypes
#' provided under ind1.gt or ind2.gt. Not used if alleles are provided in
#' ind1.hap1/ind1.hap2/ind2.hap1/ind2.hap2/
#' @param alleleNA string used as missing/unknown allele.
#' @param mismatch.code numerical, number given to loci for which there is a 
#' mismatch in genotypes between individual 1 and individual 2. Used for visualisation
#' @param match.code numerical, number given to loci for which there is a 
#' match in genotypes between individual 1 and individual 2. Used for visualisation
#' @param writeOutput logical, indicates whether results need to be written to 
#' external files
#' @param outfolder `NULL` or string, if not NULL, name of the folder in which
#' results should be saved. Will be created if it doesn't exist 
#' @param outpath string, prefix that output files will have
#' @param returnOutput logical, indicates whether results should be returned (i.e.,
#' results could also be only written to external files)
#' @param reportMismatchOnly logical, indicates whether only loci with a genotype
#' mismatch should be reported (when TRUE). If False, the outcome for all loci 
#' (including those whose genotype matched) will be reported
#'
#' @return list of two elelements: 1. numerical providing the proportion of matching
#' loci, 2. a matrix containing either the outcome for all loci (genotype match or
#' mismatch) or only reporting loci with a genotype mismatch
#' @export
#'
#' @examples
compareIndGt <- function(ind1.hap1 = NULL, ind1.hap2 = NULL, ind2.hap1 = NULL, ind2.hap2 = NULL, # for phased data / only hap1 for haploid data
                         ind1.gt = NULL, ind2.gt = NULL, # for unphased data
                         locusNames,
                         allelSep = " ",
                         alleleNA = "-",
                         mismatch.code = 0, match.code = 1,  # can be changed for visualisation
                         writeOutput = T,
                         outfolder = NULL,
                         outpath = "snpMatch_refGenome",
                         returnOutput = F,
                         reportMismatchOnly = T){
  
  if (mismatch.code == match.code){
    stop("match and mismatch code are identical, mismatches can't be identified")
  }
  
  if (all(is.null(ind1.hap1), is.null(ind1.gt))){
    stop("No genetic info provided for first individual, provide at least one genotypic data point (ind1.hap1 and ind1.hap2 for phased data, ind1.hap1 for haploid data, or ind1.gt for unphased data)")
  }
  
  if (all(!is.null(ind1.hap1), !is.null(ind1.gt))){
    stop("Both phased and unphased data provided for ind 1, provide ind1.hap1 and ind1.hap2 for phased data, ind1.hap1 for haploid data, or ind1.gt for unphased data and have the others for ind1 as NULL")
  }
  
  if (all(is.null(ind2.hap1), is.null(ind2.gt))){
    stop("No genetic info provided for first individual, provide at least one genotypic data point (ind2.hap1 and ind2.hap2 for phased data, ind2.hap1 for haploid data, or ind2.gt for unphased data)")
  }
  
  if (all(!is.null(ind2.hap1), !is.null(ind2.gt))){
    stop("Both phased and unphased data provided for ind 2, provide ind2.hap1 and ind2.hap2 for phased data, ind2.hap1 for haploid data, or ind2.gt for unphased data and have the others for ind2 as NULL")
  }
  
  if(!is.null(ind1.gt)){
    ind1.alleles <- data.frame(t(sapply(ind1.gt, function(x){
      if(is.na(x)){
        return(c(NA, NA))
      } else if(length(str_split_1(x, allelSep)) != 2){
        stop("non-diploid genotype score(s) for ind 1. Ensure all genotypes scores have exactly 2 alleles (can be missing) separated by the 'alleleSep' character")
      } else {
        return(str_split_1(x, allelSep))
      }})), stringsAsFactors = F)
    ind1.hap1 <- ind1.alleles[,1]
    ind1.hap2 <- ind1.alleles[,2]
  }
  if (!is.na(alleleNA)){
  ind1.hap1[ind1.hap1 == alleleNA] <- NA
  }
  if(!is.null(ind1.hap2) & !is.na(alleleNA)){ind1.hap2[ind1.hap2 == alleleNA] <- NA}
  
  if(!is.null(ind2.gt)){
    ind2.alleles <- data.frame(t(sapply(ind2.gt, function(x){
      if(is.na(x)){
        return(c(NA, NA))
      } else if(length(str_split_1(x, allelSep)) != 2){
        stop("non-diploid genotype score(s) for ind 2. Ensure all genotypes scores have exactly 2 alleles (can be missing) separated by the 'alleleSep' character")
      } else {
        return(str_split_1(x, allelSep))
      }})), stringsAsFactors = F)
    ind2.hap1 <- ind2.alleles[,1]
    ind2.hap2 <- ind2.alleles[,2]
  }
  if(!is.na(alleleNA)){
    ind2.hap1[ind2.hap1 == alleleNA] <- NA
  }
  if(!is.null(ind2.hap2) & !is.na(alleleNA)){ind2.hap2[ind2.hap2 == alleleNA] <- NA}
  
  compCase <- matrix(c(c(T,T,T,T),
                       c(T,T,T,F),
                       c(T,F,T,T),
                       c(T,F,T,F)), ncol = 4)
  
  case <- which(colSums(compCase == c(!is.null(ind1.hap1),
                                      !is.null(ind1.hap2),
                                      !is.null(ind2.hap1),
                                      !is.null(ind2.hap2))) == nrow(compCase))
  
  if(case == 1){
    if(length(unique(c(length(ind1.hap1), length(ind1.hap2), length(ind2.hap1), length(ind2.hap2)))) != 1){
      stop("Genotypic profiles are not of equal length. Please ensure they cover the same number of loci")
    }
    
    if(length(ind1.hap1) != length(locusNames)){
      stop("locusNames does not cover identical numbers of loci as genotypic profiles. Please ensure the number of alleles in the genotypic profiles match the number of SNP alleles in locusNames")
    }
    
    matchingLoci <- mapply(function(hap1.1, hap1.2, hap2.1, hap2.2){
      if(all(is.na(c(hap1.1, hap1.2))) | 
         all(is.na(c(hap2.1, hap2.2))) |
         (any(is.na(c(hap1.1, hap1.2))) & any(is.na(c(hap2.1, hap2.2))))){
        return(NA)
      } else if (any(is.na(c(hap1.1, hap1.2))) | any(is.na(c(hap2.1, hap2.2)))){
        if(any(na.exclude(c(hap1.1, hap1.2)) %in% na.exclude(c(hap2.1, hap2.2)))){
          return(NA)
        } else {
          return(mismatch.code)
        }
      } else if (paste0(hap1.1,hap1.2) == paste0(hap2.1,hap2.2) |
                 paste0(hap1.1,hap1.2) == paste0(hap2.2,hap2.1)){
        return(match.code)
      } else {
        return(mismatch.code)}},
      hap1.1 = ind1.hap1,
      hap1.2 = ind1.hap2,
      hap2.1 = ind2.hap1,
      hap2.2 = ind2.hap2)
  } else if (case == 2){
    if(length(unique(c(length(ind1.hap1), length(ind1.hap2), length(ind2.hap1)))) != 1){
      stop("Genotypic profiles are not of equal length. Please ensure they cover the same number of loci")
    }
    
    if(length(ind1.hap1) != length(locusNames)){
      stop("locusNames does not cover identical numbers of loci as genotypic profiles. Please ensure the number of alleles in the genotypic profiles match the number of SNP alleles in locusNames")
    }
    
    matchingLoci <- mapply(function(hap1.1, hap1.2, hap2.1){
      if(any(is.na(c(hap1.1, hap1.2, hap2.1)))){
        return(NA)
      } else if (any(c(hap1.1, hap1.2) %in% hap2.1)) {
        return(match.code)
      } else {
        return(mismatch.code)}},
      hap1.1 = ind1.hap1,
      hap1.2 = ind1.hap2,
      hap2.1 = ind2.hap1)
    
  } else if (case == 3){
    if(length(unique(c(length(ind1.hap1), length(ind2.hap1), length(ind2.hap2)))) != 1){
      stop("Genotypic profiles are not of equal length. Please ensure they cover the same number of loci")
    }
    
    if(length(ind1.hap1) != length(locusNames)){
      stop("locusNames does not cover identical numbers of loci as genotypic profiles. Please ensure the number of alleles in the genotypic profiles match the number of SNP alleles in locusNames")
    }
    
    matchingLoci <- mapply(function(hap1.1, hap2.1, hap2.2){
      if(any(is.na(c(hap1.1, hap2.1, hap2.2)))){
        return(NA)
      } else if (hap1.1 %in% c(hap2.1, hap2.2)) {
        return(match.code)
      } else {
        return(mismatch.code)}},
      hap1.1 = ind1.hap1,
      hap2.1 = ind2.hap1,
      hap2.2 = ind2.hap2)
    
  } else if (case == 4){
    if(length(ind1.hap1) != length(ind2.hap1)){
      stop("Two haplotype profiles are not of equal length. Please ensure they cover the same number of loci")
    }
    
    if(length(ind1.hap1) != length(locusNames)){
      stop("locusNames does not cover identical numbers of loci as haplotype profiles. Please ensure the number of alleles in the haplotype profiles match the number of SNP alleles in locusNames")
    }
    
    matchingLoci <- mapply(function(hap1.1, hap2.1){
      if(any(is.na(c(hap1.1, hap2.1)))){
        return(NA)
      } else if (hap1.1 ==  hap2.1) {
        return(match.code)
      } else {
        return(mismatch.code)}},
      hap1.1 = ind1.hap1,
      hap2.1 = ind2.hap1)
    
  }
  
  matchingProp <- matchingLoci[!is.na(matchingLoci)]
  matchingProp <- round(100*length(matchingProp[matchingProp == match.code])/length(matchingProp), digits = 1)
  
  locusMatch <- data.frame(cbind(locusNames, matchingLoci), stringsAsFactors = F)
  colnames(locusMatch) <- c("locusName", "gtMatch")
  if(reportMismatchOnly){
    locusMatch <- locusMatch[which(locusMatch$gtMatch == mismatch.code), ,drop=F]
  }
  matchList <- list(matchProp = matchingProp, locusmatch = locusMatch)
  
  if(writeOutput){
    if (!is.null(outfolder)){
      if (!file.exists(outfolder)){
        dir.create(outfolder)
      }
      outpath <- paste0(outfolder,"/",outpath)
    } 
    
    suppressWarnings(lapply(matchList, write.table, file = paste0(outpath, "_IdentityReport.csv"), row.names = F, append=TRUE, sep = ",", col.names = T))
  }
  
  if(returnOutput){
    return(matchList)
  }
}


#' function to determine whether allele originates from one or other haplotype
#'
#' @param ind1.hap1 vector of strings representing the alleles of the first
#' haplotype to be compared against. Expected to be the same size and order of 
#' loci as ind1.hap2/ind2.hap and genetic map
#' @param ind1.hap2 vector of strings representing the alleles of the second
#' haplotype to be compared against. Expected to be the same size and order of 
#' loci as ind1.hap1/ind2.hap and genetic map
#' @param ind2.hap vector of strings representing the alleles to determine origin
#' for. Expected to be the same size and order of loci as ind1.hap1/ind1.hap2
#' @param ind1.hap1.code numerical, number given to loci for which ind2.hap uniquely
#' matches ind1.hap1. Used for visualisation
#' @param ind1.hap2.code numerical, number given to loci for which ind2.hap uniquely
#' matches ind1.hap2. Used for visualisation
#' @param mapFile either filename if map needs to be loaded from external file or
#' name of R object. Matrix with loci as rows and at least 3 
#' columns: 1 column named "locusName" providing the name of the loci, 1 column 
#' providing linkage group info for each locus and 1 column providing positional
#' information within the linkage group
#' @param readMapFile logical; if T, an external map file with provided name 
#' will be loaded into R; if F, the provided R object will be used as map
#' @param perRegion logical; if T, results are reported per region for which all
#' loci have same origin (i.e. first and last locus and their position), otherwise,
#' the outcome for each individual locus will be reported
#' @param writeOutput logical, indicates whether results need to be written to 
#' external files
#' @param outfolder `NULL` or string, if not NULL, name of the folder in which
#' results should be saved. Will be created if it doesn't exist 
#' @param outpath string, prefix that output files will have
#' @param returnOutput logical, indicates whether results should be returned (i.e.,
#' results could also be only written to external files)
#'
#' @return a matrix containing either the outcome for all loci or only reporting
#' the origin of loci per region of single origin.
#' @export
#'
#' @examples
determineAlleleOrig <- function(ind1.hap1, ind1.hap2, ind2.hap,
                                ind1.hap1.code = 0, ind1.hap2.code = 1, misOrigin.code = -1,
                                alleleNA = "-",
                                mapFile = "mapFile.csv",
                                readMapFile = T,
                                perRegion = F,
                                writeOutput = T,
                                outfolder = NULL,
                                outpath = "snpMatch_refGenome",
                                returnOutput = F){
  
  if (readMapFile){
    if(!file.exists(mapFile)){
      stop("'mapFile' file not found. Please make sure 'mapFile' file is correctly defined (case sensitive & including filename extenstion (e.g. '.csv')")
    }
    mapFile <- read.csv(file = mapFile, header = T, stringsAsFactors = F)
  }
  
  if (!all(c("locusName", "lg", "genPos") %in% colnames(mapFile))){
    stop("Missing columns in 'mapFile'. Please make sure columns are named correctly and include 'locusName', 'lg', and 'genPos' (case sensitive)")
  }
  
  if (length(unique(c(nrow(ind1.hap1),nrow(ind1.hap2),nrow(ind2.hap)))) != 1){
    stop("Three haplotype profiles are not of equal length. Please ensure they cover the same number of loci")
  }
  
  if(length(ind1.hap1) != nrow(mapFile)){
    stop("The genetic map does not cover identical number of loci as genotype profiles. Please ensure the number of alleles in the genotype profiles match the number of SNP alleles in the genetic map")
  }
  
  if (!is.na(alleleNA)){
    ind1.hap1[ind1.hap1 == alleleNA] <- NA
    ind1.hap2[ind1.hap2 == alleleNA] <- NA
    ind2.hap[ind2.hap == alleleNA] <- NA
  }
  
  alleleOrig <- mapply(function(hap1.1 = NA, hap1.2 = NA, hap2 = NA){
    if(any(is.na(c(hap1.1, hap1.2, hap2)))){
      return(NA)
    } else if(hap1.1 == hap1.2) {
      if(hap1.1 == hap2){
        return(NA)
      } else {
        return(misOrigin.code)
      }
    } else if(hap1.1 == hap2) {
      return(ind1.hap1.code)
    } else if(hap1.2 == hap2) {
      return(ind1.hap2.code)
    } else {
      return(misOrigin.code)}},
    hap1.1 = ind1.hap1, hap1.2 = ind1.hap2, hap2 = ind2.hap)
  alleleOrig <- data.frame(cbind(mapFile, alleleOrig), stringsAsFactors = F)
  colnames(alleleOrig) <- c(colnames(mapFile), "state")
  
  if(perRegion){
    alleleOrig <- na.omit(alleleOrig)
    alleleOrig <- mapFileToList(alleleOrig)
    
    alleleOrig <- lapply(alleleOrig,
                      function(x){if (nrow(x) > 1){
                        x$prevAlleleOrig <- c("-", x$state[1:(length(x$state)-1)])
                        x$nextAlleleOrig <- c(x$state[2:length(x$state)], "-")
                        alleleOrigState.start <- x[x$state != x$prevAlleleOrig, , drop=F]
                        alleleOrigState.end <- x[x$state != x$nextAlleleOrig, , drop = F]
                        alleleOrigSumm <- data.frame(cbind(alleleOrigState.start$locusName, alleleOrigState.end$locusName, alleleOrigState.start$lg, alleleOrigState.start$genPos, alleleOrigState.end$genPos, alleleOrigState.start$state), stringsAsFactors = F)
                        colnames(alleleOrigSumm) <- c("locusName.start", "locusName.end", "lg", "pos.start", "pos.end", "state")
                        return(alleleOrigSumm)
                      } else {
                        alleleOrigSumm <- data.frame(cbind(x$locusName, x$locusName, x$lg, x$genPos, x$genPos, x$state), stringsAsFactors = F)
                        colnames(alleleOrigSumm) <- c("locusName.start", "locusName.end", "lg", "pos.start", "pos.end", "state")
                        return(alleleOrigSumm) 
                      }})
    
    alleleOrigSumm <- do.call(rbind, alleleOrig)
    
    if(writeOutput){
      if (!is.null(outfolder)){
        if (!file.exists(outfolder)){
          dir.create(outfolder)
        }
        outpath <- paste0(outfolder,"/",outpath)
      } 
      
      write.csv(alleleOrigSumm, file = paste0(outpath, "_alleleOrigSumm.csv"), row.names = F)
    }
    
    if(returnOutput){
      return(alleleOrigSumm)
    }
    
  } else {
    if(writeOutput){
      if (!is.null(outfolder)){
        if (!file.exists(outfolder)){
          dir.create(outfolder)
        }
        outpath <- paste0(outfolder,"/",outpath)
      } 
      
      write.csv(alleleOrig, file = paste0(outpath, "_alleleOrig.csv"), row.names = F)
    }
    
    if(returnOutput){
      return(alleleOrig)
    }
  }
}


#' Determines homozygous regions based on the two provided haplotypes
#'
#' @param hap1 vector of strings representing the alleles of the first haplotype.
#' Expected to be the same size and order of loci as hap2 and genetic map
#' @param hap2 vector of strings representing the alleles of the second haplotype.
#' Expected to be the same size and order of loci as hap1 and genetic map
#' @param MissVal Determines how to deal with missing values. if "skip", no state
#' of homozygosity is given to loci with missing alleles, if "exclude", these loci
#' are considered as heterozygous loci, if "include", these loci are considered
#' homozygous
#' @param hom.code numerical different from 0, number given to homozygous loci.
#' Used for visualisation
#' @param mapFile either filename if map needs to be loaded from external file or
#' name of R object. Matrix with loci as rows and at least 3 
#' columns: 1 column named "locusName" providing the name of the loci, 1 column 
#' providing linkage group info for each locus and 1 column providing positional
#' information within the linkage group
#' @param readMapFile logical; if T, an external map file with provided name 
#' will be loaded into R; if F, the provided R object will be used as map
#' @param perRegion logical; if T, results are reported per region for which all
#' loci are either homozygous or heterozygous (i.e. first and last locus and their
#' position), otherwise, the outcome for each individual locus will be reported
#' @param threshold numerical, used when homozygosity is reported per region,
#' minimum length for a region to be reported (e.g., 5 (cM))
#' @param writeOutput logical, indicates whether results need to be written to 
#' external files
#' @param outfolder `NULL` or string, if not NULL, name of the folder in which
#' results should be saved. Will be created if it doesn't exist 
#' @param outpath string, prefix that output files will have
#' @param returnOutput logical, indicates whether results should be returned (i.e.,
#' results could also be only written to external files)
#'
#' @return a matrix containing either the outcome for all loci or only reporting
#' the homozygous regions (start and end locus) larger than the provided threshold.
#' @export
#'
#' @examples
determineHomInd <- function(hap1, hap2,
                            MissVal = c("skip","exclude", "include"),
                            alleleNA = "-",
                            hom.code = 1,
                            mapFile = "mapFile.csv",
                            readMapFile = T,
                            perRegion = F,
                            threshold = 0,
                            writeOutput = T,
                            outfolder = NULL,
                            outpath = "snpMatch_refGenome",
                            returnOutput = F) {
  
  if(readMapFile){
    if(!file.exists(mapFile)){
      stop("'mapFile' file not found. Please make sure 'mapFile' file is correctly defined (case sensitive & including filename extenstion (e.g. '.csv')")
    }
    
    mapFile <- read.csv(file = mapFile, header = T, stringsAsFactors = F)
  }
  
  if (!all(c("locusName", "lg", "genPos") %in% colnames(mapFile))){
    stop("Missing columns in 'mapFile'. Please make sure columns are named correctly and include 'locusName', 'lg', and 'genPos' (case sensitive) ")
  }
  
  if (length(unique(c(length(hap1),length(hap2)))) != 1){
    stop("Two haplotype profiles are not of equal length. Please ensure they cover the same number of loci")
  }
  
  if(length(hap1) != nrow(mapFile)){
    stop("The genetic map does not cover identical number of loci as genotype profiles. Please ensure the number of alleles in the genotype profiles match the number of SNP alleles in the genetic map")
  }
  
  if (!is.na(alleleNA)){
    hap1[hap1 == alleleNA] <- NA
    hap2[hap2 == alleleNA] <- NA 
  }
  
  MissVal <- match.arg(MissVal)
  
  hom <- mapply(function(Allele1, Allele2){
    if (is.na(Allele1) |
        is.na(Allele2)){
      switch(MissVal,
             skip = return(NA),
             exclude = return("0"),
             include = return(hom.code))
    } else if (Allele1 == Allele2) {
      return(hom.code)
    } else {
      return("0")
    }},
    Allele1 = hap1, Allele2 = hap2)
  
  hom <- data.frame(cbind(mapFile, hom), stringsAsFactors = F)
  colnames(hom) <- c(colnames(mapFile), "state")
  
  if(perRegion){
    hom <- na.omit(hom)
    homList <- mapFileToList(hom)
    
    homList <- lapply(homList,
                      function(x){if (nrow(x) > 1){
                        x$prevHom <- c("-", x$state[1:(length(x$state)-1)])
                        x$nextHom <- c(x$state[2:length(x$state)], "-")
                        homState.start <- x[x$state != x$prevHom, , drop=F]
                        homState.end <- x[x$state != x$nextHom, , drop = F]
                        homSumm <- data.frame(cbind(homState.start$locusName, homState.end$locusName, homState.start$lg, homState.start$genPos, homState.end$genPos, homState.start$state), stringsAsFactors = F)
                        colnames(homSumm) <- c("locusName.start", "locusName.end", "lg", "pos.start", "pos.end", "state")
                        homSumm <- homSumm[homSumm$state == hom.code, ,drop=F]
                        homSumm <- homSumm[(as.numeric(homSumm$pos.end) - as.numeric(homSumm$pos.start)) >= threshold, ,drop=F]
                        return(homSumm)
                      } else {
                        homSumm <- cbind(x$locusName, x$locusName, x$lg, x$genPos, x$genPos, x$state)
                        colnames(homSumm) <- c("locusName.start", "locusName.end", "lg", "pos.start", "pos.end", "state")
                        homSumm <- homSumm[homSumm$state == hom.code, ,drop=F]
                        homSumm <- homSumm[(as.numeric(homSumm$pos.end) - as.numeric(homSumm$pos.start)) >= threshold, ,drop=F]
                        return(homSumm) 
                      }})
    
    homSumm <- do.call(rbind, homList)
    
    if(writeOutput){
      if (!is.null(outfolder)){
        if (!file.exists(outfolder)){
          dir.create(outfolder)
        }
        outpath <- paste0(outfolder,"/",outpath)
      } 
      
      write.csv(homSumm, file = paste0(outpath, "_homSumm.csv"), row.names = F)
    }
    if(returnOutput){
      return(homSumm)
    }
    
  } else {
    if(writeOutput){
      if (!is.null(outfolder)){
        if (!file.exists(outfolder)){
          dir.create(outfolder)
        }
        outpath <- paste0(outfolder,"/",outpath)
      } 
      
      write.csv(hom, file = paste0(outpath, "_hom.csv"), row.names = F)
    }
    if(returnOutput){
      return(hom)
    }
  }
}


#' Defines, per user-defined window, proportion of loci that did not have a unique
#' BLAST hit, are located on an unanchored sequence, or whose location is
#' inconsistent with the genetic map. 
#'
#' @param mapWgsFile external file name or R object name that contains the
#' positions of both WGS and genetic map. Must contain at least 5 columns: 1. column
#' named "locusName" containing the names of the loci, 2. column named "wgsChr"
#' containing the assigned chromosome in the WGS of each locus, 3. column named
#' "wgsPos" containing the assigned posiion in the wGS (in bp) of each locus,
#' 4. column named "lg" containing the linkage group in the genetic map of each
#' locus, 5. column named "genPos" containing the genetic position in the genetic
#' map of each locus
#' @param mismatchLoci external file name, or R object name. Matrix with at least
#' 1 column containing the names of loci that are inconsistent with the genetic 
#' map. Output from `determine.mapWgsMismatch` can be used.
#' @param readMapWGS logical; if T, an external map file with provided name 
#' will be loaded into R; if F, the provided R object will be used as mapWGS file
#' @param readMismatch logical; if T, an external file with provided name will be
#' loaded into R as `mismatchLoci` matrix; if F, the provided R object will be used
#' as `mapMismatch` matrix
#' @param chr.prefix string, prefix in chromosome name of WGS to the chromosome
#' name according to the genetic map. For example if the chromosome in the map is
#' called "1" and in the WGS the chromosome is called "chr1A", the prefix is "chr"
#' @param chr.suffix string, suffix in chromosome name of WGS to the chromosome
#' name according to the genetic map. For example if the chromosome in the map is
#' called "1" and in the WGS the chromosome is called "chr1A", the prefix is "A"
#' @param chrSepSize numerical, distance in cM that should be added between between
#' two chromosomes, for example so that a line of the given size can be drawn 
#' between chromosomes for visualisation
#' @param window numerical, size of interval over which the proportions should be calculated. 
#' @param steps numerical, distance between two consecutive intervals.To avoid 
#' overlap between intervals, `steps` should be equal to `window` (or larger)
#' @param writeOutput logical, indicates whether results need to be written to 
#' external files
#' @param outfolder `NULL` or string, if not NULL, name of the folder in which
#' results should be saved. Will be created if it doesn't exist 
#' @param outpath string, prefix that output files will have
#' @param returnOutput logical, indicates whether results should be returned (i.e.,
#' results could also be only written to external files)
#'
#' @return matrix containing the proportions of loci that did not have a unique
#' BLAST hit, are located on an unanchored sequence, or whose location is
#' inconsistent with the genetic map. 
#' @export
#'
#' @examples
AlignProp <- function(mapWgsFile = mapWgsFile,
                      mismatchLoci = mismatchLoci,
                      readMapWGS = F,
                      readMismatch = F,
                      chr.prefix,
                      chr.suffix,
                      chrSepSize = 0,
                      window = 3,
                      steps = 2,
                      writeOutput = T,
                      outfolder = NULL,
                      outpath = "snpMatch_refGenome",
                      returnOutput = F){
  if(readMapWGS){
    if(!file.exists(mapWgsFile)){
      stop("'mapWgsFile' file not found. Please make sure 'mapWgsFile' file is correctly defined (case sensitive & including filename extenstion (e.g. '.csv')")
    }
    
    mapWgsFile <- read.csv(file = mapWgsFile, header = T, stringsAsFactors = F)
  }
  
  if (!all(c("locusName", "lg", "genPos", "wgsChr", "wgsPos") %in% colnames(mapWgsFile))){
    stop("Missing columns in 'mapFile'. Please make sure columns are named correctly and include 'locusName', 'lg', and 'genPos' (case sensitive) ")
  }
  
  if(readMismatch){
    if(!file.exists(mismatchLoci)){
      stop("'mapMismatch' file not found. Please make sure 'mapMismatch' file is correctly defined (case sensitive & including filename extenstion (e.g. '.csv')")
    }
    
    mismatchLoci <- read.csv(file = mismatchLoci, header = T, stringsAsFactors = F)
  }  
  
  mapFileCumm <- addCummPos(mapWgsFile, chrSepSize = chrSepSize)
  chrNames <- unique(c(paste0(chr.prefix,mapFileCumm$lg,chr.suffix),
                       paste0(chr.prefix,"0",mapFileCumm$lg,chr.suffix),
                       NA))
  
  mapFileCumm$unanchored <- sapply(mapFileCumm$wgsChr, function(x){
    if (!(x %in% chrNames)){
      return(1)
    } else {
      return(NA)
    }})
  unanchoredLoci <- mapFileCumm[mapFileCumm$unanchored == 1, ]$locusName
  mapFileCumm$unaligned <- sapply(mapFileCumm$wgsChr, function(x){
    if(is.na(x)){
      return(1)
    } else {
      return(NA)
    }})
  mapFileCumm$mapMismatch <- sapply(mapFileCumm$locusName, function(x){
    if(x %in% mismatchLoci &
       !(x %in% unanchoredLoci)){
      return(1)
    } else {
      return(NA)
    }})
  
  mapListCumm <- mapFileToList(mapFileCumm)
  
  mapListAlignSumm <- lapply(mapListCumm, function(x){
    AlignSumm <- data.frame(matrix(nrow = 0, ncol = 6))
    colnames(AlignSumm) <- c("lg", "genPos", "cummPos", "unalignedProp", "unanchoredProp", "mapMismatchProp")
    currPos <- min(x$genPos)
    currPosCumm <- min(x$cummPos)
    while (currPos < max(x$genPos)) {
      x.temp <- subset(x, genPos >= (currPos - window/2) & genPos <= (currPos + window/2)) 
      if (nrow(x.temp) > 0){
        unanchoredProp <- round(100*nrow(x.temp[which(x.temp$unanchored == 1), ,drop = F])/nrow(x.temp),0)
        unalignedProp <- round(100*nrow(x.temp[which(x.temp$unaligned == 1), ,drop = F])/nrow(x.temp),0)
        mapMismatchProp <- round(100*nrow(x.temp[which(x.temp$mapMismatch == 1), ,drop = F])/nrow(x.temp),0)
      } else {
        unanchoredProp <- 0
        unalignedProp <- 0
        mapMismatchProp <- 0
      }
      AlignSumm <- rbind(AlignSumm, c(unique(x$lg), currPos, currPosCumm, unalignedProp, unanchoredProp, mapMismatchProp))
      AlignSumm <- data.frame(AlignSumm, stringsAsFactors = F)
      colnames(AlignSumm) <- c("lg", "genPos", "cummPos", "unalignedProp", "unanchoredProp", "mapMismatchProp")  
      currPos <- currPos + steps
      currPosCumm <- currPosCumm + steps
      
    }
    return(AlignSumm)})
  AlignSumm <- do.call(rbind, mapListAlignSumm)
  
  if(writeOutput){
    if (!is.null(outfolder)){
      if (!file.exists(outfolder)){
        dir.create(outfolder)
      }
      outpath <- paste0(outfolder,"/",outpath)
    } 
    
    write.csv(AlignSumm, file = paste0(outpath, "_AlignSumm.csv"), row.names = F)
  }
  
  if(returnOutput){
    return(AlignSumm)  
  }
}


#' Creates the input file for the circular visualisation of regions describing 
#' the origin of alleles or homozygosity. Also creates chromosome separators and
#' ring borders
#'
#' @param plotList list of lists. Each element describes a ring in the circular 
#' plot and consists of 1. a matrix named "regionDef" that defines which regions
#' of the corresponding ring should be plotted and how (what color) and 2. a 
#' vector of two elements named "ringPos" that defines the width of the ring. The
#' "regionDef" matrix has at least 3 columns: 1. a colunm named "state" containing
#' a numerical code should be plotted in (e.g., regions with haplotypes of maternal
#' origin have one code asigned, haplotypes of paternal origin have another code),
#' 2. a column named "locusName.start" indicating at which locui the regions start,
#' and 3. a column named "locusName.end" indicating at which loci the regions end.
#' The "ringPos" vector contains x-coordinates (distances from the center of the
#' circle) between which the regions listed in the corresponding matrix should
#' be displayed (i.e., the width of the ring)
#' @param xmin numerical, optional, indicates where chromosome separators should
#' start within ringplot (how close to the center of the circle). Will only be used
#' if value is closer to the center of the circle than smallest ring provided in
#' plotlist. Useful, for example, when other function(s) will create additional 
#' rings that are closer to the center and those circles need chromosome separators
#' as well.
#' @param xmax numerical, optional, indicates where chromosome separators should
#' end within ringplot (how far from the center of the circle). Will only be used
#' if value is further from the center of the circle than largest ring provided
#' in plotlist. Useful, for example, when other function(s) will create additional 
#' rings that are further from the center and those circles need chromosome 
#' separators as well.
#' @param mapFile either filename if map needs to be loaded from external file or
#' name of R object. Matrix with loci as rows and at least 3 
#' columns: 1 column named "locusName" providing the name of the loci, 1 column 
#' providing linkage group info for each locus and 1 column providing positional
#' information within the linkage group
#' @param readMapFile logical; if T, an external map file with provided name 
#' will be loaded into R; if F, the provided R object will be used as map
#' @param chrSepSize numerical, distance in cM that should be added between between
#' two chromosomes when caluclating the cummulative position across the genetic map
#' for each locus, for example so that a line of the given size can be drawn between
#' chromosomes for visualisation.
#' @param ringSepsize numerical, thickness of the separator (border) between rings
#' @param sep.code numerical different from 0, number given to chromosome and ring
#' separators in the output matrix. Used for determining their color during 
#' visualisation
#' @param writeOutput logical, indicates whether results need to be written to 
#' external files
#' @param outfolder `NULL` or string, if not NULL, name of the folder in which
#' results should be saved. Will be created if it doesn't exist 
#' @param outpath string, prefix that output files will have
#' @param returnOutput logical, indicates whether results should be returned (i.e.,
#' results could also be only written to external files)
#'
#'
#' @return matrix containing the dimensions of the regions to be plotted with 
#' respective code used for coloring as well as separators for each chromosome
#' and ring. Can be used as input for `createRingPlot.plot` function
#' @export
#'
#' @examples
createRingPlot.region <- function(plotList,
                                  xmin = NA,
                                  xmax = NA,
                                  mapFile = "mapFile.csv",
                                  readMapFile = T,
                                  chrSepSize = 0,
                                  ringSepsize = 0.05,
                                  sep.code = 99,
                                  writeOutput = T,
                                  outfolder = NULL,
                                  outpath = "snpMatch_refGenome",
                                  returnOutput = F){
  if(readMapFile){
    if(!file.exists(mapFile)){
      stop("'mapFile' file not found. Please make sure 'mapFile' file is correctly defined (case sensitive & including filename extenstion (e.g. '.csv')")
    }
    
    mapFile <- read.csv(file = mapFile, header = T, stringsAsFactors = F)
  }
  
  if (!all(c("locusName", "lg", "genPos") %in% colnames(mapFile))){
    stop("Missing columns in 'mapFile'. Please make sure columns are named correctly and include 'locusName', 'lg', and 'genPos' (case sensitive) ")
  }
  
  mapFileCumm <- addCummPos(mapFile = mapFile, chrSepSize = chrSepSize)
  plotFile <- data.frame()
  
  for(i in 1:length(plotList)){
    plotFile.temp <- subset(plotList[[i]]$regionDef, select = c(state, locusName.start, locusName.end))
    plotFile.temp$state <- plotFile.temp$state
    plotFile.temp$ymin <- mapFileCumm[match(plotList[[i]]$regionDef$locusName.start, mapFileCumm$locusName),]$cummPos
    plotFile.temp$ymax <- mapFileCumm[match(plotList[[i]]$regionDef$locusName.end, mapFileCumm$locusName),]$cummPos
    plotFile.temp$xmin <- plotList[[i]]$ringPos[1]
    plotFile.temp$xmax <- plotList[[i]]$ringPos[2]
    plotFile.temp <- subset(plotFile.temp, select = c(state, ymin, ymax, xmin, xmax))
    plotFile <- rbind(plotFile, plotFile.temp)
  }
  
  xrange <- unique(c(plotFile$xmin,plotFile$xmax))
  xrange <- xrange[order(xrange)]
  ymin <- 0
  ymax <- max(mapFileCumm$cummPos)+chrSepSize
  
  for (i in 1:length(unique(mapFileCumm$lg))){
    plotFile <- rbind(plotFile, c(sep.code,
                                  max(mapFileCumm[mapFileCumm$lg == unique(mapFileCumm$lg)[i], ]$cummPos),
                                  max(mapFileCumm[mapFileCumm$lg == unique(mapFileCumm$lg)[i], ]$cummPos)+chrSepSize,
                                  min(xrange, xmin, na.rm = T), max(xrange, xmax, na.rm = T)))
  }
  for (i in 1:length(xrange)){
    plotFile <- rbind(plotFile, c(sep.code,
                                  ymin, ymax,
                                  xrange[i] - (ringSepsize/2),
                                  xrange[i] + (ringSepsize/2)))
  }
  
  if(writeOutput){
    if (!is.null(outfolder)){
      if (!file.exists(outfolder)){
        dir.create(outfolder)
      }
      outpath <- paste0(outfolder,"/",outpath)
    } 
    
    write.csv(plotFile, file = paste0(outpath, "_RegionPlotDef.csv"), row.names = F)
  }
  
  return(plotFile)
}

#' Creates the input file for the circular visualisation of proportion of loci
#' that do not have a unique WGS position assigned, that are aligned to unanchored
#' regions, and that are inconsistent with the genetic map
#'
#' @param AlignErrFile either filename if AlignErrFile needs to be loaded from
#' external file or name of R object. Matrix with genetic regions as rows and at
#' least 4 columns: 1. column named "cummPos" with the cummulative position in the
#' genetic map of each region, 2. column named "unalignedProp" with the proportions
#' (0-100) of loci that do not have a unique WGS position assigned for each region,
#' 3. column named "unanchoredProp" with the proportions (0-100) of loci that are
#' aligned to unanchored regions, and 4. column named "mapMismatchProp" with the
#' proportion (0-100) of loci that are inconsistent with the genetic map. Output
#' from the function `AlignProp` can be used
#' @param readAlignErrFile logical; if T, an external AlignErrFile with provided
#' name will be loaded into R; if F, the provided R object will be used as 
#' AlignErrFile
#' @param xmin numerical, x-value (distance from the circle center) where 0% should
#' be in the plot. Can be higher than xmax for "inverse" plot (i.e. peaks of high
#' proportions pointing inwards)
#' @param xmax numerical, x-value (distance from the circle center) where 100% 
#' should be in the plot. Can be lower than xmin for "inverse" plot (i.e. peaks 
#' of high proportions pointing inwards)
#'
#' @return matrix containing the proportions of loci that did not have a unique
#' BLAST hit, are located on an unanchored sequence, or whose location is
#' inconsistent with the genetic map, adjusted for the plotting area. Can be used
#' as input for `createRingPlot.plot` function
#' @export
#'
#' @examples
createRingPlot.align <- function(AlignErrFile = "AlignErrFile.csv",
                                 readAlignErrFile = T,
                                 xmin,
                                 xmax){
  xrange <- xmax-xmin
  AlignErrFile$unalign.start <- xmin

  AlignErrFile$unalign.end <- AlignErrFile$unalign.start + xrange*AlignErrFile$unalignedProp/100
  AlignErrFile$unanch.end <- AlignErrFile$unalign.end + xrange*AlignErrFile$unanchoredProp/100
  AlignErrFile$misalign.end <- AlignErrFile$unanch.end + xrange*AlignErrFile$mapMismatchProp/100
  
  alignPlotFile <- subset(AlignErrFile, select = c(cummPos, unalign.start, unalign.end, unanch.end, misalign.end))
  return(alignPlotFile)
}

#' Creates the input file for the circular visualisation of single loci
#'
#' @param locusNames either filename if locusNames needs to be loaded from
#' external file or name of R object. Matrix with loci names that need to be plotted
#' with at least 2 columns: 1. column named "locusName" containing the names of 
#' the loci to be plotted and 2. column with named defined under `locus,code` that
#' contains the numerical codes that indicate how loci should be colored
#' @param locus.code string, name of columns that contains the numerical codes 
#' that indicate how loci should be colored
#' @param mapFile either filename if map needs to be loaded from external file or
#' name of R object. Matrix with loci as rows and at least 3 
#' columns: 1 column named "locusName" providing the name of the loci, 1 column 
#' providing linkage group info for each locus and 1 column providing positional
#' information within the linkage group
#' @param readlocusNames logical; if T, an external locusNames file with provided
#' name will be loaded into R; if F, the provided R object will be used as locusNames
#' file
#' @param readMapFile logical; if T, an external map file with provided name 
#' will be loaded into R; if F, the provided R object will be used as map
#' @param xmin numerical, x-value (distance from the circle center) where ring
#' with plotted loci should start
#' @param xmax numerical, x-value (distance from the circle center) where ring
#' with plotted loci should end
#' @param locusWidth numerical, width that a single locus should have in the plot
#' (in cM if map is in cM; a single (SNP) locus normally has a width of zero and
#' would not be visible, therefore a thicker width should be provided by the user)
#' @param chrSepSize numerical, distance in cM that should be added between between
#' two chromosomes when caluclating the cummulative position across the genetic map
#' for each locus, for example so that a line of the given size can be drawn between
#' chromosomes for visualisation.
#' @param ringSepsize numerical, thickness of the separator (border) at each side
#' of the ring diplaying the loci
#' @param sep.code numerical different from 0, number given to chromosome and ring
#' separators in the output matrix. Used for determining their color during 
#' visualisation
#'
#' @return matrix containing the dimensions of the loci to be plotted with 
#' respective code used for coloring as well as separators for each chromosome
#' and ring. Can be used as input for `createRingPlot.plot` function
#' @export
#'
#' @examples
createRingplot.locus <- function(locusNames = "locusNames.csv",
                                 locus.code = colname.code,
                                 mapFile = "mapFile.csv",
                                 readlocusNames = T,
                                 readMapFile = T,
                                 xmin,
                                 xmax,
                                 locusWidth = 2,
                                 chrSepSize = 0,
                                 ringSepsize = 0.05,
                                 sep.code = 99){
  
  if(readlocusNames){
    if(!file.exists(locusNames)){
      stop("'locusNames' file not found. Please make sure 'locusNames' file is correctly defined (case sensitive & including filename extenstion (e.g. '.csv')")
    }
    
    locusNames <- read.csv(file = locusNames, header = T, stringsAsFactors = F)
  }
  
  if (!all(c("locusName",locus.code) %in% colnames(locusNames))){
    stop("Missing columns in 'locusNames'. Please make sure columns are named correctly and include 'locusName' and the name defined uner 'locus.code' (case sensitive) ")
  }
  
  if(readMapFile){
    if(!file.exists(mapFile)){
      stop("'mapFile' file not found. Please make sure 'mapFile' file is correctly defined (case sensitive & including filename extenstion (e.g. '.csv')")
    }
    
    mapFile <- read.csv(file = mapFile, header = T, stringsAsFactors = F)
  }
  
  if (!all(c("locusName", "lg", "genPos") %in% colnames(mapFile))){
    stop("Missing columns in 'mapFile'. Please make sure columns are named correctly and include 'locusName', 'lg', and 'genPos' (case sensitive) ")
  }
  
  mapFileCumm <- addCummPos(mapFile = mapFile, chrSepSize = chrSepSize)
  ymin <- 0
  ymax <- max(mapFileCumm$cummPos) + chrSepSize
  
  if(nrow(locusNames)>0){
    plotFile <- locusNames
    plotFile$state <- locusNames[,which(colnames(locusNames) == locus.code)]
    plotFile$ymin <- mapFileCumm[match(locusNames$locusName, mapFileCumm$locusName),]$cummPos - locusWidth/2
    plotFile$ymax <- mapFileCumm[match(locusNames$locusName, mapFileCumm$locusName),]$cummPos + locusWidth/2
    plotFile$xmin <- xmin
    plotFile$xmax <- xmax
    plotFile <- subset(plotFile, select = c(state, ymin, ymax, xmin, xmax))
    plotFile <- rbind(plotFile, c(sep.code,
                                  ymin, ymax,
                                  xmin - ringSepsize/2,
                                  xmin + ringSepsize/2))
    plotFile <- rbind(plotFile, c(sep.code,
                                  ymin, ymax,
                                  xmax - ringSepsize/2,
                                  xmax + ringSepsize/2))
  } else {
    plotFile <- data.frame(matrix(, nrow = 0, ncol = 5))
    plotFile <- rbind(plotFile, c(sep.code,
                                  ymin, ymax,
                                  xmin - ringSepsize/2,
                                  xmin + ringSepsize/2))
    plotFile <- rbind(plotFile, c(sep.code,
                                  ymin, ymax,
                                  xmax - ringSepsize/2,
                                  xmax + ringSepsize/2))
    colnames(plotFile) <- c("state", "ymin", "ymax", "xmin", "xmax")
  }
  
  return(plotFile)
}

#' Function that uses various input files to create a circular plot representing
#' the allele origins of alleles in the WGSs as well as genotype mismatches, regions
#' of homozygosity, and BLAST alignment summary per region
#'
#' @param regionTable either filename if regionTable needs to be loaded from
#' external file or name of R object. Matrix containing at least 5 columns: 1. 
#' column named "state", containing a numerical code representing the state of a
#' region (e.g., maternal allel, paternal allele, homozygous, genotype mismatch).
#' "state" column determines the color in the plot; 2. column named "ymin" providing
#' start cumulative position in the genetic map of each region; 3. column named
#' "ymax" providing end cumulative position in the genetic map of each region;
#' 4. column named "xmin" indicating how far from the circle center the displaying
#' of each region starts; 5. column named "xmax" indicating how far from the circle
#' center the displaying of each region ends. Togeteher with "xmin", it determines
#' the width of the region. Output from `createRingPlot.region` or 
#' `createRingplot.locus` can be used.
#' @param colorDef  either filename if regionTable needs to be loaded from external
#' file or name of R object. Matrix containing at least 3 columns: 1. column named
#' "state" containing the numerical codes representing the state of the region as
#' used in the `regionTable` file; 2. column named "color", containing color codes
#' that indicate what color each "state" code should be displayed in; 3. column
#' named "legend" which contains description of each "state"/color.
#' @param missAlignTable.hap1 either filename if regionTable needs to be loaded 
#' from external file or name of R object. Matrix containing the info to plot per
#' region in the genetic map the proportion of loci that do not have a unique 
#' location in the first haplome, that align to unanchored regions in the first
#' haplome, and that align to regions in the first haplome which are inconsistent
#' to the  genetic map. This matrix has at least 4 columns: 1. column named 
#' "unalign.start" indicating for each window in the genetic map how far from the
#' circle's center the coloring should start representing the proportion of loci
#' without a unique alignment; 2. column named "unalign.end" indicating for each 
#' window in the genetic map how far from the circle's center the coloring should 
#' end representing the proportion of loci without a unique alignment. This is 
#' also where the coloring starts representing the proportion of loci that were 
#' aligned to unachored regions; 3 column named "unanch.end" indicating for each
#' window in the genetic map how far from the circle's center the coloring should
#' end representing the proportion of loci that were aligned to unachored regions.
#' This is also where the coloring starts representing the proportion of loci that
#' aligned to region inconsistent with the genetic map; 4. column named 
#' "misalign.end" indicating for each window in the genetic map how far from the
#' circle's center the coloring should end representing the proportion of loci 
#' that aligned to region inconsistent with the genetic map. Output from 
#' `createRingPlot.align` can be used
#' @param missAlignTable.hap2 either filename if regionTable needs to be loaded 
#' from external file or name of R object. Matrix similar to `missAlignTable.hap1`
#' but containing the information for the second haplome instead. Output from 
#' `createRingPlot.align` can be used
#' @param readRegionT logical; if T, an external regionTable with provided name 
#' will be loaded into R; if F, the provided R object will be used as regionTable
#' @param readColorDef logical; if T, an external colorDef file with provided name 
#' will be loaded into R; if F, the provided R object will be used as colorDef
#' @param readMissAlignT.hap1 logical; if T, an external MissAlignT.hap1 file
#' with provided name will be loaded into R; if F, the provided R object will be
#' used as MissAlignT.hap1
#' @param readMissAlignT.hap2 logical; if T, an external MissAlignT.hap2 file 
#' with provided name will be loaded into R; if F, the provided R object will be
#' used as MissAlignT.hap2
#' @param unalignCol color code representing the color used for the proportion of
#' loci without unique location per window in the genetic map
#' @param unanchCol color code representing the color used for the proportion of
#' loci aligning to unanchored regions per window in the genetic map
#' @param misalignCol color code representing the color used for the proportion of
#' loci aligning to regions inconsistent with the genetic map per window in the
#' genetic map
#' @param  logical, indicates whether results need to be written to 
#' external files
#' @param outfolder `NULL` or string, if not NULL, name of the folder in which
#' results should be saved. Will be created if it doesn't exist 
#' @param outpath string, prefix that output files will have
#' @param plot.width numerical, indicating the width of the plot that will be 
#' generated
#' @param plot.units unit of `plot.width, can be "in", "cm", "mm", or "px"
#' @param plot.dpi numerical indicating plot resolution
#' @param returnOutput logical, indicates whether results should be returned (i.e.,
#' results could also be only written to external files)
#' 
#' @return circular plot representing the allele origins of alleles in the WGSs
#' as well as genotype mismatches, regions of homozygosity, and BLAST alignment
#' summary per region
#' @export
#'
#' @examples
createRingPlot.plot <- function(regionTable,
                                colorDef,
                                missAlignTable.hap1 = NULL,
                                missAlignTable.hap2 = NULL,
                                readRegionT = T,
                                readColorDef = T,
                                readMissAlignT.hap1 = T,
                                readMissAlignT.hap2 = T,
                                unalignCol = "darkgrey",
                                unanchCol = "#000000",
                                misalignCol = "#F92241",
                                writeOutput = T,
                                outfolder = NULL,
                                outpath = "snpMatch_refGenome",
                                plot.width = 1500,
                                plot.units = "px",
                                plot.dpi = 300,
                                returnOutput = F){
  if(readRegionT){
    if(!file.exists(regionTable)){
      stop("'regionTable' file not found. Please make sure 'regionTable' file is correctly defined (case sensitive & including filename extenstion (e.g. '.csv')")
    }
    regionTable <- read.csv(file = regionTable, header = T, stringsAsFactors = F)
  }
  if (!all(c("state", "xmin", "xmax", "ymin", "ymax") %in% colnames(regionTable))){
    stop("Missing columns in 'regionTable'. Please make sure columns are named correctly and include 'state', 'xmin', 'xmax', 'ymin', and 'ymax' (case sensitive)")
  }
  
  if(readColorDef){
    if(!file.exists(colorDef)){
      stop("'colorDef' file not found. Please make sure 'colorDef' file is correctly defined (case sensitive & including filename extenstion (e.g. '.csv')")
    }
    colorDef <- read.csv(file = colorDef, header = T, stringsAsFactors = F)
  }
  if (!all(c("state", "color", "legend") %in% colnames(colorDef))){
    stop("Missing columns in 'colorDef'. Please make sure columns are named correctly and include 'state', 'color', and 'legend' (case sensitive)")
  }
  colorDefPlot <- colorDef[colorDef$state %in% unique(regionTable$state),]
  
  if(!is.null(missAlignTable.hap1)){
    if(readMissAlignT.hap1){
      if(!file.exists(missAlignTable.hap1)){
        stop("'missAlignTable.hap1' file not found. Please make sure 'missAlignTable.hap1' file is correctly defined (case sensitive & including filename extenstion (e.g. '.csv')")
      }
      missAlignTable.hap1 <- read.csv(file = missAlignTable.hap1, header = T, stringsAsFactors = F)
    }
    if (!all(c("unalign.end", "unalign.start", "unanch.end", "misalign.end") %in% colnames(missAlignTable.hap1))){
      stop("Missing columns in 'missAlignTable.hap1'. Please make sure columns are named correctly and include 'unalign.end', 'unalign.start', 'unanch.end', and 'misalign.end' (case sensitive)")
    }
  }
  
  if (!is.null(missAlignTable.hap2)) {
    if(readMissAlignT.hap2){
      if(!file.exists(missAlignTable.hap2)){
        stop("'missAlignTable.hap2' file not found. Please make sure 'missAlignTable.hap2' file is correctly defined (case sensitive & including filename extenstion (e.g. '.csv')")
      }
      missAlignTable.hap2 <- read.csv(file = missAlignTable.hap2, header = T, stringsAsFactors = F)
    }
    if (!all(c("unalign.end", "unalign.start", "unanch.end", "misalign.end") %in% colnames(missAlignTable.hap2))){
      stop("Missing columns in 'missAlignTable.hap2'. Please make sure columns are named correctly and include 'unalign.end', 'unalign.start', 'unanch.end', and 'misalign.end' (case sensitive)")
    }
  }
  
  ringPlot =  ggplot2::ggplot() + 
              ggplot2::coord_polar(theta= "x") +
              ggplot2::ylim(c(min(1.5,(min(c(regionTable$xmin, missAlignTable.hap1$unalign.start, missAlignTable.hap1$misalign.end, missAlignTable.hap2$unalign.start, missAlignTable.hap2$misalign.end))-1.5)),
                max(c(regionTable$xmax, missAlignTable.hap1$unalign.start, missAlignTable.hap1$misalign.end, missAlignTable.hap2$unalign.start, missAlignTable.hap2$misalign.end)))) +
              ggplot2::theme_bw() +
              ggplot2::theme(panel.grid= ggplot2::element_blank()) +
              ggplot2::theme(axis.text= ggplot2::element_blank()) +
              ggplot2::theme(axis.ticks= ggplot2::element_blank(), panel.border = ggplot2::element_blank()) +
      
              ggplot2::geom_rect(data = regionTable, ggplot2::aes(fill=as.factor(state),
                                                ymax=xmax,
                                                ymin=xmin, 
                                                xmax=ymax, 
                                                xmin=ymin)) +
      
              ggplot2::scale_fill_manual(values = colorDefPlot$color,
                                name="Color Code",
                                breaks=colorDefPlot$state,
                                labels=colorDefPlot$legend)
  
    
  if(!is.null(missAlignTable.hap1)){
    ringPlot <- ringPlot +
        
      ggplot2::geom_ribbon(data = missAlignTable.hap1, ggplot2::aes(x = cummPos, y = unalign.end, ymin = unalign.start, ymax = unalign.end), fill=unalignCol) +
        
      ggplot2::geom_ribbon(data = missAlignTable.hap1, ggplot2::aes(x = cummPos, y = unanch.end, ymin = unalign.end, ymax = unanch.end), fill=unanchCol) +
        
      ggplot2::geom_ribbon(data = missAlignTable.hap1, ggplot2::aes(x = cummPos, y = misalign.end, ymin = unanch.end, ymax = misalign.end), fill=misalignCol)
        
  }
  
  if(!is.null(missAlignTable.hap2)){
    ringPlot <- ringPlot +
      
      ggplot2::geom_ribbon(data = missAlignTable.hap2, ggplot2::aes(x = cummPos, y = unalign.end, ymin = unalign.start, ymax = unalign.end), fill=unalignCol) +
      
      ggplot2::geom_ribbon(data = missAlignTable.hap2, ggplot2::aes(x = cummPos, y = unanch.end, ymin = unalign.end, ymax = unanch.end), fill=unanchCol) +
      
      ggplot2::geom_ribbon(data = missAlignTable.hap2, ggplot2::aes(x = cummPos, y = misalign.end, ymin = unanch.end, ymax = misalign.end), fill=misalignCol)
  }
  

  if(writeOutput){
    if (!is.null(outfolder)){
      if (!file.exists(outfolder)){
        dir.create(outfolder)
      }
      outpath <- paste0(outfolder,"/",outpath)
    } 
    
    ggplot2::ggsave(filename = paste0(outpath,"_ringplot.pdf"),
                     plot = ringPlot,
                     device = "pdf",
                     width = plot.width,
                     units = plot.units,
                     dpi = plot.dpi)
  }
  
  if(returnOutput){
    return(ringPlot)
  }  
}


###### Wrapper Functions ########

phasedWgsVsSNP <- function(parFile = "parFile.csv"){
  
  print("** main step 1: Loading parameters & checking data files **")
  
  ## create a set of default parameters in case of missing parameters
  defaultPar <- data.frame(rbind(c("analysisType", "all", "# default = all; should be 'blast', 'refInd', or 'all' - If 'blast', BLAST results are compared against a genetic map and the best position for each locus is reported, as well as any inconsistencies and the genetic profile (genotypes) for the (loci of) the genetic map. If 'refInd', the genetic profile of the WGS will be compared against a reference profile (e.g., from SNP array data) from a specific individual and visualize the outcome, as well as visualizise the outcome from the BLAST analysis. If 'all', all of the analyses will be performed"),
                                 c("wgsHap1.fasta", NA, "# required if analysisType is 'all' or 'blast'; no default - Filename of Genome sequence for first haplome, haploid or collapsed genome. Fasta file format and division into chromosomes/pseudomolecules is expected"),
                                 c("wgsHap1.blastFull", NA, "# required if analysisType is 'all' or 'blast'; no default - Filename of Blast output for (SNP) markers, usually from provided flanking or probe sequences. Blast output format 6 is expected"),
                                 c("wgsHap1.short", "NULL", "# only used if analysisType is 'all' or 'blast'; default = NULL - Filename of Blast output for (SNP) markers using shorter sequences if available. Used to avoid gaps in allignment and improve determination of marker location. Blast output format 6 is expected. Use NULL when not used"),
                                 c("chr.prefix.hap1", "chr", "# default = chr - prefix used before chromosome number in haplome 1 (case sensitive). For example, in 'Chr1A', prefix is 'Chr' and script will check for 'Chr1' and 'Chr01'. NULL if no prefix present"),                                                                                                                                                                                                             
                                 c("chr.suffix.hap1", "NULL", "# default = NULL - suffix used after aomosome number in haplome 1 (case sensitive). For example, in 'Chr1A', suffix is 'A'. NULL if no suffix present"),
                                 c("uniqPos.hap1", NA, "# required if analysisType is 'refInd'; no default - Filename of document containing the (unique) WGS position of reference loci in the first haplome. If loci do not have a unique position, they should not be included in the document. Three named columns are expected 'locusName', 'wgsChr', 'wgsPos' (case sensitive), containing the names of the loci, the chromosome they aligned to, and their nucleotide position on the chromosome. csv file format required. This file can also be created using 'all' or 'blast' as 'analysisType'"),
                                 c("wgsProf.hap1", NA, "# required if analysisType is 'refInd'; no default - Filename of document containing the DNA profile of the first haplome. First row should contain the name of the loci and the second row should contain the alleles present for these loci in the first haplome. csv file format required. This file can also be created using 'all' or 'blast' as 'analysisType'"),
                                 c("mapMismatch.hap1", NA, "# required if analysisType is 'refInd'; no default - Filename of document containing loci whose positions in the first haplome are inconsistent with the genetic map. Contains at least one column with 'locusName' as header which has the names of the loci that are inconsistent. csv file format required. This file can also be created using 'all' or 'blast' as 'analysisType'"),
                                 c("wgsHap2.fasta", "NULL", "# only used if analysisType is 'all' or 'blast'; default = NULL - Filename of Genome sequence for second haplome in case of phased whole genome sequence. Fasta file format and division into chromosomes/pseudomolecules is expected. Chromosome numbering is expected to identical between the two haplomes. NULL if not present"),
                                 c("wgsHap2.blastFull", "NULL", "# required if analysisType is 'all' or 'blast' AND wgsHap2.fasta is not NULL; default = NULL - Filename of Blast output for (SNP) markers, usually from provided flanking or probe sequences. Blast output format 6 is expected"),
                                 c("wgsHap2.short", "NULL", "# only used if analysisType is 'all' or 'blast' AND wgsHap2.fasta is not NULL; default = NULL - (only used if wgsHap2.fasta is provided) - Filename of Blast output for (SNP) markers using shorter sequences. Used to avoid gaps in allignment and improve determination of marker location. Blast output format 6 is expected. Use NULL when not used"), 
                                 c("chr.prefix.hap2", "chr", "# only used if wgsHap2.fasta or uniqPos.hap2 is not NULL; default = chr - prefix used before chromosome number in haplome 2 (case sensitive). For example, in 'Chr1B', prefix is 'Chr' and script will check for 'Chr1' and 'Chr01'. NULL if no prefix present"),
                                 c("chr.suffix.hap2", "NULL", "# only used if wgsHap2.fasta or uniqPos.hap2 is not NULL; default = NULL - suffix used after aomosome number in haplome 2 (case sensitive). For example, in 'Chr1B', suffix is 'B'. NULL if no suffix present"),
                                 c("uniqPos.hap2", "NULL", "# only used if analysisType is 'refInd'; default = NULL - Filename of document containing the (unique) WGS position of reference loci in the second haplome. If loci do not have a unique position, they should not be included in the document. Three named columns are expected 'locusName', 'wgsChr', 'wgsPos' (case sensitive), containing the names of the loci, the chromosome they aligned to, and their nucleotide position on the chromosome. csv file format required. This file can also be created using 'all' or 'blast' as 'analysisType'"),
                                 c("wgsProf.hap2", "NULL", "# required if analysisType is 'refInd' AND uniqPos.hap2 is not NULL; default = NULL - Filename of document containing the DNA profile of the second haplome (in case of phased WGS). First row should contain the name of the loci and the second row should contain the alleles present for these loci in the first haplome. csv file format required. This file can also be created using 'all' or 'blast' as 'analysisType'"),
                                 c("mapMismatch.hap2", "NULL", "# required if analysisType is 'refInd' AND uniqPos.hap2 is not NULL; default = NULL - Filename of document containing loci whose positions in the second haplome are inconsistent with the genetic map. Contains at least one column with 'locusName' as header which has the names of the loci that are inconsistent. csv file format required. This file can also be created using 'all' or 'blast' as 'analysisType'"), 
                                 c("snpInfo", NA, "# required if analysisType is 'all' or 'blast'; no default - Filename of document that provides info on position of SNPs in the used sequences for the Blast Analyses. This file requires at least two columns with 'snpName' and 'flSeqSnpPos.full' as headers (case sensitive) . If Blast results using short sequences are also provided, a column with 'flSeqSnpPos.short' as header is also required. This file list for each SNP used in the BLAST analyses (as rows), the corresonding SNP's name used in the BLAST analyses (snpName), the position of the SNP in the sequence provided for the full BLAST analysis (flSeqSnpPos.full), and, if provided, the position of the SNP in the sequence provided for the BLAST analysis with short sequences (flSeqSnpPos.short). In case of probes, the SNP's postion can be just outside the provided sequence, in that case the position can be 0 (if SNP is just upstream of provided sequence) or length of the probe + 1 (if SNP is just downstream of provided sequence). csv file format required"),
                                 c("mapFile", NA, "# required; no default - Filename of genetic map file including (some of) the (SNP) markers used in the BLAST analyses to improve the determination of the position of markers in the whole genome sequences. This file requires at least three columns with 'locusName', 'lg', and 'genPos' as headers (case sensitive) and list the SNP name (locusName), its linkage group in the genetic map as a number (lg), and its genetic position in cM in the genetic map (genPos). Only a sinlge location is expected for each SNP in the genetic map. Ideally, numbering of linkage groups matches that of the WGS, if not markers position can be accurately determined but will be listed as incorrect position. csv file format required"),
                                 c("length.hit", 15, "# only used if analysisType is 'all' or 'blast'; default = 15 - indicating minimum length of a full BLAST hit to be considered for WGS position determination"),
                                 c("dist.bp", 0, "# only used if analysisType is 'all' or 'blast'; default = 0 - distance in bp that a found hit can deviate from the genomic region determined by flanking markers. Any hit located further from this region will not be considered when multiple hits exist. Hits located further from the genomic region determined by flanking markers than the value provided will be considered as incorrect locations in the analyses"),
                                 c("dist.cM", 2.5, "# only used if analysisType is 'all' or 'blast'; default = 2.5 - distance in cM within which flanking loci with unique BLAST hit should be looked for on the genetic map"),
                                 c("nloci", "NULL", "# only used if analysisType is 'all' or 'blast'; default = NULL - NULL or numerical, number of upstream/downstream loci to consider when determining the best BLAST hit. If NULL, all loci within the provided window (dist.cM) will be used"),
                                 c("mhaplotypes", NA, "# required if analysisType is 'all' or 'refInd'; no default - Filename containing phased reference genetic data for at least one individual to compare against WGS. Follows the mhaplotypes file format of FlexQTL. One row with at least 'ID' and SNP names used in genetic map (and BLAST analyses) expected but does not need to be the first row. This row will be used as column names and is immediately followed by phased genetic data for individuals. Two rows per individual (one per homolog), each starting with the individual's name and followed by phased marker scores in corresponding columns (based on column names). Marker scores are expected to either follow the AB format from Illumina Infinium SNP Arrays ('A' for 'A' or 'T'; 'B' for 'C' or 'G'), or uncleotide format ('A', 'C', 'G', or 'T'). csv file format expected"),
                                 c("indName", NA, "# required if analysisType is 'all' or 'refInd'; no default - Individual name as provided in mhaplotypes to be compared against WGS"),
                                 c("convertABfmt", T, "# only used if analysisType is 'all' or 'blast'; default = T - T or F depending on whether AB format or Nucleotide format is used in mhaplotypes file, respectively"),
                                 c("alleleNA", "-", "# default = '-' - Character(s) used for missing alleles in the mhaplotypes file and (generated) WGS profile"),                                                                                                                                                                                                                    
                                 c("skipGap", T, "# only used if analysisType is 'all' or 'blast'; default = T - T or F for excluding alignment with gaps from the evaluation or not, respectively"),                                                                                                                                                                                                                     
                                 c("alignErr.window", 5, "# only used if analysisType is 'all' or 'refInd'; default = 5 - cM window accross which proportion of alignments inconcsistent with map, alignments to unanchored regions, and unaligned markers are summarized"),
                                 c("alignErr.steps", 5, "# only used if analysisType is 'all' or 'refInd'; default = 5 - cM steps for between consequtive windows defined under alignErr.window"),
                                 c("hom.MissVal", "skip", "# only used if analysisType is 'all' or 'refInd'; default = skip - either 'skip', 'exclude', or 'include' (case sensitive). Indicates how missing genotypic data should be handled for calculating homozygous regions. 'skip' removes missing data from the analyses (thus practically considering missing data as homozygous when it is within a homozygous region and as heterozygous when in a heterozygous region or at the end of a homozygous region (thus not further extending the homozygous region), 'exclude' considers all missing data as heterozygous, and 'include' considers all missing data as homozygous"),
                                 c("hom.threshold",5, "# only used if analysisType is 'all' or 'refInd'; default = 5 - size in cM for a continuous section of homozygous markers to be considered as a homozygous region. For homozygous regions, allele origin cannot be accurately determined"),
                                 c("exclude.GTmismatch", T, "# only used if analysisType is 'all' or 'refInd'; default = T - T or F for excluding or including genotype mismatches with reference (SNP array) genotypic data, respectively. Inclusion can be useful when only a relative is available as reference data"),
                                 c("wgsHap1.range", "c(4,5)", "# only used if analysisType is 'all' or 'refInd'; default = c(4,5) - Y-coordinates of start and end for displaying circle with allele origin of haplome 1. Determines width of circle. Provided as 'c(x,y)' with x and y the start and end coordinate, respectively"),
                                 c("misalignHap1.range", "c(3.975,3.225)", "# only used if analysisType is 'all' or 'refInd'; default = c(3.975,3.225) - Y-coordinates of start and end for displaying circle with allignment issues for haplome 1. Determines width of circle. Provided as 'c(x,y)' with x and y the start and end coordinate, respectively, in which x can be larger than y to invert the plot (i.e. peaks pointing inwards)"),
                                 c("wgsHap2.range", "c(5.5,6.5)", "# only used if analysisType is 'all' or 'refInd' AND uniqPos.hap2 is not NULL; default = c(5.5,6.5) - Y-coordinates of start and end for displaying circle with allele origin of haplome 2. Determines width of circle. Provided as 'c(x,y)' with x and y the start and end coordinate, respectively"),
                                 c("misalignHap2.range", "c(6.525,7.275)", "# only used if analysisType is 'all' or 'refInd' AND uniqPos.hap2 is not NULL; default = c(6.525,7.275) - Y-coordinates of start and end for displaying circle with allignment issues for haplome 2. Determines width of circle. Provided as 'c(x,y)' with x and y the start and end coordinate, respectively, in which x can be larger than y to invert the plot (i.e. peaks pointing inwards)"),
                                 c("mismatch.range", "c(5.25,5.5)", "# only used if analysisType is 'all' or 'refInd'; default = c(5.25,5.5) - Y-coordinates of start and end for displaying circle with genotype inconsistencies between WGS and reference marker data. Provided as 'c(x,y)' with x and y the start and end coordinate, respectively"),
                                 c("mismatch.width", 1, "# only used if analysisType is 'all' or 'refInd'; default = 1 - width in cM for diplaying markers with genotypic inconsistencies between WGS and reference data. As single (SNP) markers cover no or limited interval, they would occur as a very fine line that is not or hard to observe. This parameter enables the user to set a wider interval (e.g., 1 or 2 cM) for clearly displaying the markers with inconsitency"),
                                 c("hom.range", "c(5,5.25)", "# only used if analysisType is 'all' or 'refInd'; default = c(5,5.25) - Y-coordinates of start and end for displaying circle with homozygous regions (as allele origin cannot be assigned there). Provided as 'c(x,y)' with x and y the start and end coordinate, respectively"),
                                 c("chrSepSize", 2, "# only used if analysisType is 'all' or 'refInd'; default = 2 - width (in cM) of the separator lines between each chromosome in the plot"),
                                 c("ringSepsize", 0.05, "# only used if analysisType is 'all' or 'refInd'; default = 0.05 - width of separator lines between two rings The center of these separator lines will be positioned at the y-coordinates provided under 'wgsHap1.range', 'mismatch.range', 'hom.range', and, when used, 'wgsHap2.range'. It will cover the provided size of y-coordinates "),
                                 c("hap1.col", "gold2", "# only used if analysisType is 'all' or 'refInd'; default = gold2 - color code used for displaying alleles of the first homolog (maternal alleles in plants) of individual chosen under 'indName'. Provided as either color name recognized by R or as hex color code"),
                                 c("hap2.col", "forestgreen", "# only used if analysisType is 'all' or 'refInd'; default = forestgreen - color code used for displaying alleles of the second homolog (paternal alleles in plants) of individual chosen under 'indName'. Provided as either color name recognized by R or as hex color code"),
                                 c("hom.col", "dodgerblue", "# only used if analysisType is 'all' or 'refInd'; default = dodgerblue - color code used for displaying homozygous regions of individual chosen under 'indName'. Provided as either color name recognized by R or as hex color code"),
                                 c("misOrigin.col","white","# only used if analysisType is 'all' or 'refInd'; default = white - color used when allele in WGS is not found in reference and thus no origin can be assigned. Such cases will only occur if `exclude.GTmismatch` is set to false" ),
                                 c("mismatch.col", "red3", "# only used if analysisType is 'all' or 'refInd'; default = red3 - color code used for displaying genotypic inconsistencies between reference data and WGS. Provided as either color name recognized by R or as hex color code"),
                                 c("unalign.col", "darkgrey", "# only used if analysisType is 'all' or 'refInd'; default = darkgrey - color code used for displaying proportion of markers that were aligned to unanchored sequences (if applicable). Provided as either color name recognized by R or as hex color code"), 
                                 c("unanch.col", "black", "# only used if analysisType is 'all' or 'refInd'; default = black - color code used for displaying proportion of markers for which no unique position could be found in the WGS. Provided as either color name recognized by R or as hex color code"),
                                 c("misalign.col", "red3", "# only used if analysisType is 'all' or 'refInd'; default = red3 - color code used for displaying proportion of markers for whose positions are inconsistent with the provided genetic map. Provided as either color name recognized by R or as hex color code"),
                                 c("ringOutlineCol", "black", "# only used if analysisType is 'all' or 'refInd'; default = black - color code used for displaying separators between chromosomes and rings. Provided as either color name recognized by R or as hex color code"),
                                 c("hap1.label", "Maternal Haplotypes", "# only used if analysisType is 'all' or 'refInd'; default = Maternal Haplotypes - label provided in ringplot for alleles in hap2.col"),
                                 c("hap2.label", "Paternal Haplotypes", "# only used if analysisType is 'all' or 'refInd'; default = Paternal Haplotypes - label provided in ringplot for alleles in hap1.col"),
                                 c("plot.width", 1500, "# only used if analysisType is 'all' or 'refInd'; default = 1500 - width of the ringplot to be generated. A number is expected, the units depend on the 'plot.units' parameter"),
                                 c("plot.units", "px", "# only used if analysisType is 'all' or 'refInd'; default = px - unit for the width of the ringplot to be generated. Can be either 'in', 'cm', 'mm', or 'px'"),
                                 c("plot.dpi", 300, "# only used if analysisType is 'all' or 'refInd'; default = 300 - Determines plot resolution for the ringplot to be generated"),
                                 c("returnOutput", T, "default = T - T or F to determine whether results should be returned to be saved in an R object or not, respectively"),
                                 c("writeOutput", T, "default = T - T or F to determine whether results should be saved to external files"),
                                 c("outfolder", "wgsAnalysis", "# only used if writeOutput is T; default = wgsAnalysis - name of folder in which to put all output, will be created if folder does not exists. Use NULL when not used"),
                                 c("outpath", "wgsAnalysis", "# only used if writeOutput is T; default = wgsAnalysis - Provides the prefix that will be used for all created files")),
                           stringsAsFactors = F)
  colnames(defaultPar) = c("par", "defaultVal", "comment")
  
  #loading parameter file with all other file names and parameter values
  if(!file.exists(parFile)){
    write.csv(defaultPar, "exampleParFile.csv", row.names = F, quote = T)
    
    stop("'parFile' file not found. Please make sure 'parFile' file is correctly defined (case sensitive & including filename extenstion ('.csv'). 'exampleParFile.csv' created")
  }
  parFile <- read.csv(parFile, header = F)
  
  ## reading from text file
  # parFile <- readLines("ParFile as Text.txt")
  # while (max(lengths(regmatches(parFile, gregexpr("\t\t", parFile)))) > 0){
  #   parFile <- gsub("\t\t", "\t", parFile)
  # }
  # parFile <- read.table(text = parFile, sep="\t", header = F,  comment.char = "#", fill = T)
  
  if (!(ncol(parFile) == 2 | ncol(parFile) == 3)){
    stop("'parFile' does not have two or three columns. Either two columns are expected, the first column having parameter names and the second column with corresponding parameter values, or three columns are expected which includes an additional column with comments (each comment starting with #)")
  }
  if (ncol(parFile) == 2){
    colnames(parFile) = c("par", "val")
  } else {
    colnames(parFile) = c("par", "val", "comment")
  }
  
  analysisT <- parFile[which(parFile$par == "analysisType"),]$val
  
  if (!length(analysisT) == 0){
    if (!analysisT %in% c("all", "blast", "refInd")){
      stop("analysisType should be 'all', `blast', or 'refInd' (case sensitive)")
    }
  } else {
    analysisT <- "all"
    print("'analysisType' not defined, default of 'all' will be used.")
  }
  
  parFile <- parFile[which(!is.na(parFile$val)),]
  
  if(analysisT %in% c("all","blast")){
    parNeeded <- c("wgsHap1.fasta", "wgsHap1.blastFull",
                   "mapFile", "snpInfo")
    parCheck <- parNeeded %in% parFile$par
    if (!all(parCheck)){
      stop(paste0("The following parameters are missing from the parameter file without default option. Ensure they are included (case sensitive): ", paste(parNeeded[!parCheck], collapse = ", ")))
    }
    parOptional <- c("wgsHap1.short", "wgsHap2.fasta", "wgsHap2.blastFull", "wgsHap2.short",
                     "length.hit", "dist.bp", "dist.cM", "nloci", "convertABfmt", "skipGap") 
  } else {
    parOptional <- NULL
  } 
  
  if(analysisT %in% c("all","refInd")){
    parNeeded <- c("mhaplotypes", "indName")
    parCheck <- parNeeded %in% parFile$par
    if (!all(parCheck)){
      stop(paste0("The following parameters are missing from the parameter file without default option. Ensure they are included (case sensitive): ", paste(parNeeded[!parCheck], collapse = ", ")))
    }
    parOptional <- c(parOptional, "alleleNA", "alignErr.window", "alignErr.steps", "hom.MissVal", "hom.threshold", "exclude.GTmismatch",
                     "wgsHap1.range", "misalignHap1.range", "wgsHap2.range", "misalignHap2.range", "mismatch.range", "mismatch.width", "hom.range", "chrSepSize", "ringSepsize",
                     "hap1.col", "hap2.col", "hom.col", "misOrigin.col", "mismatch.col", "unalign.col", "unanch.col", "misalign.col","ringOutlineCol", "hap1.label", "hap2.label", "plot.width", "plot.units", "plot.dpi",
                     "returnOutput", "writeOutput", "outfolder", "outpath")
  } 
  
  if(analysisT == "refInd"){
    parNeeded <- c("mapFile", "uniqPos.hap1", "wgsProf.hap1","mapMismatch.hap1")
    parCheck <- parNeeded %in% parFile$par
    if (!all(parCheck)){
      stop(paste0("The following parameters are missing from the parameter file without default option. Ensure they are included (case sensitive): ", paste(parNeeded[!parCheck], collapse = ", ")))
    }
    parOptional <- c(parOptional, "uniqPos.hap2", "wgsProf.hap2","mapMismatch.hap2")
  }

  parOptional <- c(parOptional,"analysisType", "chr.prefix.hap1", "chr.suffix.hap1", "chr.prefix.hap2", "chr.suffix.hap2",
                   "returnOutput", "writeOutput", "outfolder", "outpath")
  
  
  parOptCheck <- parOptional %in% parFile$par
  if (!all(parOptCheck)){
    print(paste0("The following parameters are missing from the parameter file: ",paste(parOptional[!parOptCheck], collapse = ", "), ". Default values will be used. Otherwise, ensure these parameters are included in the paramter file (case sensitive)"))
    
    #set default values for parameters used during analyses
    defaultPar$providedVal <- parFile[match(defaultPar$par,parFile$par), ]$val
    defaultPar$usedVal <- mapply(function(x,y){if(!is.na(y)){return(y)} else {(return(x))}},
                                 x = defaultPar$defaultVal,
                                 y = defaultPar$providedVal)
    parFile <- subset(defaultPar, select= c(par, usedVal, comment))
    colnames(parFile) = c("par", "val", "comment")
  }
  
  parFile$valParse <- mapply(function(x){
    if(!is.na(x)){
      if(is.na(suppressWarnings(as.numeric(x))) & is.na(suppressWarnings(as.logical(x))) & !startsWith(x, "c(") & x != "NULL"){paste0("\"",x,"\"")
      } else {return(x)}
    } else {return(x)}}, x = parFile$val)
    
  for(i in 1:nrow(parFile)){
    eval(parse(text = paste(parFile$par[i]," <- ", parFile$valParse[i]) ))
  }
  
  #check all other files
  
  ## Check map File 
  if(!file.exists(mapFile)){
    stop("'mapFile' file not found. Please make sure 'mapFile' file is correctly defined (case sensitive & including filename extenstion ('.csv')")
  }
  mapFile <- read.csv(file = mapFile, header = T, stringsAsFactors = F)
  if (!all(c("locusName", "lg", "genPos") %in% colnames(mapFile))){
    stop("Missing columns in 'mapFile'. Please make sure columns are named correctly and include 'locusName', 'lg', and 'genPos' (case sensitive)")
  }
  
  if (analysisT %in% c("all","blast")){
    ## Check fasta file for haplome 1
    if(!file.exists(wgsHap1.fasta)){
      stop("'wgsHap1.fasta' file not found. Please make sure 'wgsHap1.fasta' file is correctly defined (case sensitive & including filename extenstion (e.g. '.fasta')")
    }
    ## Check Blast Output of full sequences for haplome 1
    if(!file.exists(wgsHap1.blastFull)){
      stop("'wgsHap1.blastFull' file not found. Please make sure 'wgsHap1.blastFull' file is correctly defined (case sensitive & including filename extenstion (e.g. '.out')")
    } 
    wgsHap1.blastFull <- data.frame(read.table(file = wgsHap1.blastFull), stringsAsFactors = F)
    if(ncol(wgsHap1.blastFull) != 12){
      stop("Unexpected Number of Columns in Provided 'wgsHap1.blastFull' file. Please check it was generated using the '-outfmt 6' command and contains the following columns 'querry name', 'subject name', 'proportion identitiy', 'lenght of Match', 'number mismatches', 'number gaps', 'querry Start', 'querry End', 'subject Start', 'subject gEnd', 'eValue', and 'bitScore'")
    }
    
    ## Check Blast Output of short sequences for haplome 1 if present
    if(!is.null(wgsHap1.short)){
      if(!file.exists(wgsHap1.short)){
        stop("'wgsHap1.short' file not found. Please make sure 'wgsHap1.short' file is correctly defined (case sensitive & including filename extenstion (e.g. '.out')")
      }
      wgsHap1.short <- data.frame(read.table(file = wgsHap1.short), stringsAsFactors = F)
      if(ncol(wgsHap1.short) != 12){
        stop("Unexpected Number of Columns in Provided 'wgsHap1.short' file. Please check it was generated using the '-outfmt 6' command and contains the following columns 'querry name', 'subject name', 'proportion identitiy', 'lenght of Match', 'number mismatches', 'number gaps', 'querry Start', 'querry End', 'subject Start', 'subject gEnd', 'eValue', and 'bitScore'")
      }
    }
    
    # Check files for haplome 2 if provided
    if(!is.null(wgsHap2.fasta)){
      ## Check fasta file for haplome 2
      if(!file.exists(wgsHap2.fasta)){
        stop("'wgsHap2.fasta' file not found. Please make sure 'wgsHap2.fasta' file is correctly defined (case sensitive & including filename extenstion (e.g. '.fasta')")
      }
      ## Check Blast Output of full sequences for haplome 2
      if(!file.exists(wgsHap2.blastFull)){
        stop("'wgsHap2.blastFull' file not found. Please make sure 'wgsHap2.blastFull' file is correctly defined (case sensitive & including filename extenstion (e.g. '.out')")
      }
      wgsHap2.blastFull <- data.frame(read.table(file = wgsHap2.blastFull), stringsAsFactors = F)
      if(ncol(wgsHap2.blastFull) != 12){
        stop("Unexpected Number of Columns in Provided 'wgsHap2.blastFull' file. Please check it was generated using the '-outfmt 6' command and contains the following columns 'querry name', 'subject name', 'proportion identitiy', 'lenght of Match', 'number mismatches', 'number gaps', 'querry Start', 'querry End', 'subject Start', 'subject gEnd', 'eValue', and 'bitScore'")
      }
      ## Check Blast Output of short sequences for haplome 2 if present
      if(!is.null(wgsHap2.short)){
        if(!file.exists(wgsHap2.short)){
          stop("'wgsHap2.short' file not found. Please make sure 'wgsHap2.short' file is correctly defined (case sensitive & including filename extenstion (e.g. '.out')")
        }
        wgsHap2.short <- data.frame(read.table(file = wgsHap2.short), stringsAsFactors = F)
        if(ncol(wgsHap2.short) != 12){
          stop("Unexpected Number of Columns in Provided 'wgsHap2.short' file. Please check it was generated using the '-outfmt 6' command and contains the following columns 'querry name', 'subject name', 'proportion identitiy', 'lenght of Match', 'number mismatches', 'number gaps', 'querry Start', 'querry End', 'subject Start', 'subject gEnd', 'eValue', and 'bitScore'")
        }
      }
    }
    
    ## Check SNP info File
    if(!is.null(snpInfo)){
      if(!file.exists(snpInfo)){
        stop("'snpInfo' file not found. Please make sure 'snpInfo' file is correctly defined (case sensitive & including filename extenstion ('.csv')")
      }
      snpInfo <- read.csv(file = snpInfo, header = T, stringsAsFactors = F)
      
      if(is.null(wgsHap1.short) & is.null(wgsHap2.short)){
        if(!all(c("snpName", "flSeqSnpPos.full") %in% colnames(snpInfo))){
          stop("Missing columns in 'snpInfo'. Please make sure columns are named correctly and include 'snpName', 'flSeqSnpPos.full' (case sensitive)")
        }
      } else {
        if(!all(c("snpName", "flSeqSnpPos.full","flSeqSnpPos.short") %in% colnames(snpInfo))){
          stop("Missing columns in 'snpInfo'. Please make sure columns are named correctly and include 'snpName', 'flSeqSnpPos.full', 'flSeqSnpPos.short' (case sensitive). If no BLAST was run on short sequences, ensure wgsHap1.short and wgsHap2.short are set to NULL in the parameter file")
        }
      }
    }
  }
  
  if (analysisT %in% c("all","refInd")){
    ## Check mhaplotype file & if needed, sort it according to mapFile (and introduce missing data for markers in map but not in mhaplotypes)
    if(!file.exists(mhaplotypes)){
      stop("'mhaplotypes' file not found. Please make sure 'mhaplotypes' file is correctly defined (case sensitive & including filename extenstion ('.csv')")
    }
    mhaplotypes <- read.mhaplotypes(mhaplotypes)
    if(!(indName %in% mhaplotypes$ID)){
      stop("'indName' not found in mhaplotypes. Please ensure that individual's name is identically named to mhaplotypes (case sensitive) and present in mhaplotypes")
    }
    mhaplotypes.ind <- mhaplotypes[which(mhaplotypes$ID == indName), ]
    if ("VAR" %in% colnames(mhaplotypes.ind)){
      mhaplotypes.ind <- mhaplotypes.ind[which(mhaplotypes.ind$VAR %in% c("hap1", "hap2")), ]
    }
    if(nrow(mhaplotypes.ind) > 2){
      stop("More than two haplotypes found for individual defined under 'indName'. Please ensure only 2 haplotypes are present")
    }
    mhaplotypes.ind <- data.frame(t(t(mhaplotypes.ind)[match(mapFile$locusName, colnames(mhaplotypes.ind)), 1:2]), stringsAsFactors = F)
    colnames(mhaplotypes.ind) <- mapFile$locusName
    
  }
  
  if (analysisT == "refInd"){
    if(!file.exists(uniqPos.hap1)){
      stop("'uniqPos.hap1' file not found. Please make sure 'uniqPos.hap1' file is correctly defined (case sensitive & including filename extenstion ('.csv')")
    }
    uniqPos.hap1 <- read.csv(file = uniqPos.hap1, header = T, stringsAsFactors = F) 
    if (!all(c("locusName", "wgsChr", "wgsPos") %in% colnames(uniqPos.hap1))){
      stop("Missing columns in 'uniqPos.hap1'. Please make sure columns are named correctly and include 'locusName', 'wgsChr', and 'wgsPos' (case sensitive)")
    }
    mapFile.hap1 <- mapFile
    mapFile.hap1$wgsChr <- uniqPos.hap1[match(mapFile$locusName, uniqPos.hap1$locusName),]$wgsChr
    mapFile.hap1$wgsPos <- uniqPos.hap1[match(mapFile$locusName, uniqPos.hap1$locusName),]$wgsPos
    
    if(!file.exists(wgsProf.hap1)){
      stop("'wgsProf.hap1' file not found. Please make sure 'wgsProf.hap1' file is correctly defined (case sensitive & including filename extenstion ('.csv')")
    }
    wgsHap1.snpProf <- read.csv(file = wgsProf.hap1, header = T, stringsAsFactors = F)
    wgsHap1.snpProf <- data.frame(t(t(wgsHap1.snpProf)[match(mapFile$locusName, colnames(wgsHap1.snpProf)), ]), stringsAsFactors = F)
    colnames(wgsHap1.snpProf) <- mapFile$locusName
    
    
    if(!file.exists(mapMismatch.hap1)){
      stop("'mapMismatch.hap1' file not found. Please make sure 'mapMismatch.hap1' file is correctly defined (case sensitive & including filename extenstion ('.csv')")
    }
    wgsHap1.mapMismatch <- read.csv(file = mapMismatch.hap1, header = T, stringsAsFactors = F)
    if (!("locusName" %in% colnames(wgsHap1.mapMismatch))){
      stop("Missing column in 'mapMismatch.hap1'. Please make sure columns are named correctly and include 'locusName' (case sensitive)")
    }
    
    if(!is.null(uniqPos.hap2)){
      if(!file.exists(uniqPos.hap2)){
        stop("'uniqPos.hap2' file not found. Please make sure 'uniqPos.hap2' file is correctly defined (case sensitive & including filename extenstion ('.csv')")
      }
      uniqPos.hap2 <- read.csv(file = uniqPos.hap2, header = T, stringsAsFactors = F) 
      if (!all(c("locusName", "wgsChr", "wgsPos") %in% colnames(uniqPos.hap2))){
        stop("Missing columns in 'uniqPos.hap2'. Please make sure columns are named correctly and include 'locusName', 'wgsChr', and 'wgsPos' (case sensitive)")
      }
      mapFile.hap2 <- mapFile
      mapFile.hap2$wgsChr <- uniqPos.hap2[match(mapFile$locusName, uniqPos.hap2$locusName),]$wgsChr
      mapFile.hap2$wgsPos <- uniqPos.hap2[match(mapFile$locusName, uniqPos.hap2$locusName),]$wgsPos
      
      if(!file.exists(wgsProf.hap2)){
        stop("'wgsProf.hap2' file not found. Please make sure 'wgsProf.hap2' file is correctly defined (case sensitive & including filename extenstion ('.csv')")
      }
      wgsHap2.snpProf <- read.csv(file = wgsProf.hap2, header = T, stringsAsFactors = F)
      wgsHap2.snpProf <- data.frame(t(t(wgsHap2.snpProf)[match(mapFile$locusName, colnames(wgsHap2.snpProf)), ]), stringsAsFactors = F)
      colnames(wgsHap2.snpProf) <- mapFile$locusName
      
      
      if(!file.exists(mapMismatch.hap2)){
        stop("'mapMismatch.hap1' file not found. Please make sure 'mapMismatch.hap1' file is correctly defined (case sensitive & including filename extenstion ('.csv')")
      }
      wgsHap2.mapMismatch <- read.csv(file = mapMismatch.hap2, header = T, stringsAsFactors = F)
      if (!("locusName" %in% colnames(wgsHap1.mapMismatch))){
        stop("Missing column in 'mapMismatch.hap1'. Please make sure columns are named correctly and include 'locusName' (case sensitive)")
      }
    } else {wgsHap2.snpProf <- NULL}
  }

  if(writeOutput){
    if (!is.null(outfolder)){
      if (!file.exists(outfolder)){
        dir.create(outfolder)
      }
      outpath <- paste0(outfolder,"/",outpath)
    }
    
    write.csv(subset(parFile, select= c(par, val, comment)), paste0(outpath,"_parametersUsed.csv"), row.names = F, quote = T)
  }
  
  if (analysisT %in% c("all","blast")){
    print("** main step 2: Determining best SNP locations wgs hap1 (10 steps) **")
    wgsHap1.blastResults <- blastAnalysis(blastResults.full = wgsHap1.blastFull,
                                          readBlast.full = F,
                                          blastResults.short = wgsHap1.short,
                                          readBlast.short = F,
                                          chr.prefix = chr.prefix.hap1,
                                          chr.suffix = chr.suffix.hap1,
                                          snpInfo = snpInfo,
                                          readSnpInfo = F,
                                          mapFile = mapFile,
                                          readMapFile = F,
                                          length.hit = length.hit,
                                          dist.bp = dist.bp,
                                          dist.cM = dist.cM,
                                          nloci = nloci,
                                          writeOutput = writeOutput,
                                          outfolder = NULL,
                                          outpath = paste0(outpath,"_hap1"),
                                          returnOutput = T)$uniqueHits
  
    print("** main step 3: Determining mismatches between WGS hap1 locations and genetic map **")
    mapFile.hap1 <- mapFile
    mapFile.hap1$wgsChr <- wgsHap1.blastResults[match(mapFile$locusName, wgsHap1.blastResults$locusName),]$wgsChr
    mapFile.hap1$wgsPos <- wgsHap1.blastResults[match(mapFile$locusName, wgsHap1.blastResults$locusName),]$wgsPos
    wgsHap1.mapMismatch <- determine.mapWgsMismatch(mapWgsFile = mapFile.hap1,
                                                    readMapWGS = F,
                                                    chr.prefix = chr.prefix.hap1,
                                                    chr.suffix = chr.suffix.hap1,
                                                    dist.bp = dist.bp,
                                                    dist.cM = dist.cM,
                                                    nloci = nloci,
                                                    writeOutput = writeOutput,
                                                    outfolder = NULL,
                                                    outpath = paste0(outpath,"_hap1"),
                                                    returnOutput = T)
    
    print("** main step 4: Determining alleles present in wgs hap1 (2 steps) **")
    wgsHap1.snpAlleles <- retrieveSnpAllele(wgs = wgsHap1.fasta,
                                            readWgs = T,
                                            snpPos = wgsHap1.blastResults,
                                            readSnpPos = F,
                                            skipGap = skipGap,
                                            convertABfmt = convertABfmt,
                                            writeOutput = writeOutput,
                                            outfolder = NULL,
                                            outpath = paste0(outpath,"_hap1"),
                                            returnOutput = T)
    
    print("** main step 5: Creating wgs hap1 profile **")
    wgsHap1.snpProf <- createProfile(wgsAlleles = wgsHap1.snpAlleles,
                                     mapFile = mapFile,
                                     mapMismatch = wgsHap1.mapMismatch,
                                     readAlleles = F,
                                     readMapFile = F,
                                     readMismatch = F,
                                     naString = "-",
                                     writeOutput = writeOutput,
                                     outfolder = NULL,
                                     outpath = paste0(outpath,"_hap1"),
                                     returnOutput = T)
    
    if(!is.null(wgsHap2.fasta)){
      print("** main step 6: Determining best SNP locations wgs hap2 (10 steps) **")
      wgsHap2.blastResults <- blastAnalysis(blastResults.full = wgsHap2.blastFull,
                                            readBlast.full = F,
                                            blastResults.short = wgsHap1.short,
                                            readBlast.short = F,
                                            chr.prefix = chr.prefix.hap2,
                                            chr.suffix = chr.suffix.hap2,
                                            snpInfo = snpInfo,
                                            readSnpInfo = F,
                                            mapFile = mapFile,
                                            readMapFile = F,
                                            length.hit = length.hit,
                                            dist.bp = dist.bp,
                                            dist.cM = dist.cM,
                                            nloci = nloci,
                                            writeOutput = writeOutput,
                                            outfolder = NULL,
                                            outpath = paste0(outpath,"_hap2"),
                                            returnOutput = T)$uniqueHits
  
      print("** main step 7: Determining mismatches between WGS hap2 locations and genetic map **")
      mapFile.hap2 <- mapFile
      mapFile.hap2$wgsChr <- wgsHap2.blastResults[match(mapFile$locusName, wgsHap2.blastResults$locusName),]$wgsChr
      mapFile.hap2$wgsPos <- wgsHap2.blastResults[match(mapFile$locusName, wgsHap2.blastResults$locusName),]$wgsPos
      wgsHap2.mapMismatch <- determine.mapWgsMismatch(mapWgsFile = mapFile.hap2,
                                                      readMapWGS = F,
                                                      chr.prefix = chr.prefix.hap2,
                                                      chr.suffix = chr.suffix.hap2,
                                                      dist.bp = dist.bp,
                                                      dist.cM = dist.cM,
                                                      nloci = nloci,
                                                      writeOutput = writeOutput,
                                                      outfolder = NULL,
                                                      outpath = paste0(outpath,"_hap2"),
                                                      returnOutput = T)
  
      
      print("** main step 8: Determining alleles present in wgs hap2 (2 steps) **")
      wgsHap2.snpAlleles <- retrieveSnpAllele(wgs = wgsHap2.fasta,
                                              readWgs = T,
                                              snpPos = wgsHap2.blastResults,
                                              readSnpPos = F,
                                              skipGap = skipGap,
                                              convertABfmt = convertABfmt,
                                              writeOutput = writeOutput,
                                              outfolder = NULL,
                                              outpath = paste0(outpath,"_hap2"),
                                              returnOutput = T)
      
      print("** main step 9: Creating wgs hap2 profile **")
      wgsHap2.snpProf <- createProfile(wgsAlleles = wgsHap2.snpAlleles,
                                       mapFile = mapFile,
                                       mapMismatch = wgsHap2.mapMismatch,
                                       readAlleles = F,
                                       readMapFile = F,
                                       readMismatch = F,
                                       naString = "-",
                                       writeOutput = writeOutput,
                                       outfolder = NULL,
                                       outpath = paste0(outpath,"_hap2"),
                                       returnOutput = T)
    
    } else {
      print("** main step 5-9 skipped: no second haploid WGS to analyze **")
      wgsHap2.snpProf <- NULL
    }
  }
  
  if (analysisT %in% c("all","refInd")){
    if(analysisT == "refInd"){
      print("** main step 2: Check for genotypic inconsistencies **")
    } else {
      print("** main step 10: Check for genotypic inconsistencies **")
    }
    
    wgsProf <-  rbind(mhaplotypes.ind, wgsHap1.snpProf, wgsHap2.snpProf)
    
    if(!is.null(wgsHap2.snpProf)){
      wgsVsSNPprof.mismatch <- compareIndGt(ind1.hap1 = wgsProf[1,], ind1.hap2 = wgsProf[2,], ind2.hap1 = wgsProf[3,], ind2.hap2 = wgsProf[4,],
                                            locusNames = colnames(wgsHap1.snpProf),
                                            alleleNA = alleleNA,
                                            mismatch.code = 0, match.code = 1,
                                            writeOutput = writeOutput,
                                            outpath = outpath,
                                            outfolder = NULL,
                                            returnOutput = T,
                                            reportMismatchOnly = T)
      
      if(exclude.GTmismatch){
        wgsProf[3:4, colnames(wgsProf) %in% wgsVsSNPprof.mismatch$locusmatch$locusName] <- NA
      }
    } else {
      
      wgsVsSNPprof.mismatch <- compareIndGt(ind1.hap1 = wgsProf[1,], ind1.hap2 = wgsProf[2,], ind2.hap1 = wgsProf[3,], ind2.hap2 = NULL,
                                            locusNames = colnames(wgsHap1.snpProf),
                                            alleleNA = alleleNA,
                                            mismatch.code = 0, match.code = 1,
                                            writeOutput = writeOutput,
                                            outpath = outpath,
                                            outfolder = NULL,
                                            returnOutput = T,
                                            reportMismatchOnly = T)
      
      if(exclude.GTmismatch){
        wgsProf[3:4, colnames(wgsProf) %in% wgsVsSNPprof.mismatch$locusmatch$locusName] <- NA
      }
    }
    
    if(analysisT == "refInd"){
      print("** main step 3: Creating ringplot files for hap1 **")
    } else {
      print("** main step 11: Creating ringplot files for hap1 **")
    }  

    wgsHap1.wgsHapOrigin <- determineAlleleOrig(ind1.hap1 = wgsProf[1,], ind1.hap2 = wgsProf[2,], ind2.hap = wgsProf[3,],
                                                ind1.hap1.code = 1, ind1.hap2.code = 2, misOrigin.code = -1, 
                                                alleleNA = alleleNA,
                                                mapFile = mapFile,
                                                readMapFile = F,
                                                perRegion = T,
                                                writeOutput = writeOutput,
                                                outfolder = NULL,
                                                outpath = paste0(outpath,"_hap1"),
                                                returnOutput = T)
    
    wgsHap1.alignErrSumm <- AlignProp(mapWgsFile = mapFile.hap1,
                                      mismatchLoci = wgsHap1.mapMismatch$locusName,
                                      readMapWGS = F,
                                      readMismatch = F,
                                      chr.prefix = chr.prefix.hap1,
                                      chr.suffix = chr.suffix.hap1,
                                      chrSepSize = chrSepSize,
                                      window = alignErr.window,
                                      steps = alignErr.steps,
                                      writeOutput = writeOutput,
                                      outfolder = NULL,
                                      outpath = paste0(outpath,"_hap1"),
                                      returnOutput = T)
    
    wgsHap1.alignRing <- createRingPlot.align(AlignErrFile = wgsHap1.alignErrSumm, readAlignErrFile = F, xmin = misalignHap1.range[1], xmax = misalignHap1.range[2])
    
    wgs.hom <- determineHomInd(hap1 = wgsProf[1,], hap2 = wgsProf[2,],
                               alleleNA = alleleNA,
                               MissVal = hom.MissVal,
                               hom.code = 11,
                               mapFile = mapFile,
                               readMapFile = F,
                               perRegion = T,
                               threshold = hom.threshold,
                               writeOutput = writeOutput,
                               outfolder = NULL,
                               outpath = outpath,
                               returnOutput = T)
  
    mismatchPlot <- createRingplot.locus(locusNames = wgsVsSNPprof.mismatch$locusmatch,
                                         locus.code = "gtMatch",
                                         mapFile = mapFile,
                                         readMapFile = F,
                                         readlocusNames = F,
                                         xmin = mismatch.range[1],
                                         xmax = mismatch.range[2],
                                         locusWidth = mismatch.width,
                                         chrSepSize = chrSepSize,
                                         ringSepsize = ringSepsize,
                                         sep.code = 99)
    
    
    if(!is.null(wgsHap2.snpProf)){  
      if(analysisT == "refInd"){
        print("** main step 4: Creating ringplot files for hap2 **")
      } else {
        print("** main step 12: Creating ringplot files for hap2 **")
      }
      
      wgsHap2.wgsHapOrigin <- determineAlleleOrig(ind1.hap1 = wgsProf[1,], ind1.hap2 = wgsProf[2,], ind2.hap = wgsProf[4,],
                                                  ind1.hap1.code = 1, ind1.hap2.code = 2, misOrigin.code = -1,
                                                  alleleNA = alleleNA,
                                                  mapFile = mapFile,
                                                  readMapFile = F,
                                                  perRegion = T,
                                                  writeOutput = writeOutput,
                                                  outfolder = NULL,
                                                  outpath = paste0(outpath,"_hap2"),
                                                  returnOutput = T)
      
      wgsHap2.alignErrSumm <- AlignProp(mapWgsFile = mapFile.hap2,
                                        mismatchLoci = wgsHap2.mapMismatch$locusName,
                                        readMapWGS = F,
                                        readMismatch = F,
                                        chr.prefix = chr.prefix.hap2,
                                        chr.suffix = chr.suffix.hap2,
                                        chrSepSize = chrSepSize,
                                        window = alignErr.window,
                                        steps = alignErr.steps,
                                        writeOutput = writeOutput,
                                        outfolder = NULL,
                                        outpath = paste0(outpath,"_hap2"),
                                        returnOutput = T)
  
      wgsHap2.alignRing <- createRingPlot.align(AlignErrFile = wgsHap2.alignErrSumm, readAlignErrFile = F, xmin = misalignHap2.range[1], xmax = misalignHap2.range[2])
  
      ringTable <- list(list(regionDef = wgsHap1.wgsHapOrigin, ringPos = wgsHap1.range),
                        list(regionDef = wgsHap2.wgsHapOrigin, ringPos = wgsHap2.range),
                        list(regionDef = wgs.hom, ringPos = hom.range))
      
      regionPlot <- createRingPlot.region(plotList =  ringTable,
                                          xmin = min(c(wgsHap1.range, wgsHap2.range, hom.range, misalignHap1.range, misalignHap2.range, mismatch.range)),
                                          xmax = max(c(wgsHap1.range, wgsHap2.range, hom.range, misalignHap1.range, misalignHap2.range, mismatch.range)),
                                          mapFile = mapFile,
                                          readMapFile = F,
                                          chrSepSize = chrSepSize,
                                          ringSepsize = ringSepsize,
                                          sep.code = 99,
                                          writeOutput = F,
                                          outfolder = NULL,
                                          outpath = "snpMatch_refGenome",
                                          returnOutput = T)
      
      regionPlot <- rbind(regionPlot, mismatchPlot)
      
    } else {
      ringTable <- list(list(regionDef = wgsHap1.wgsHapOrigin, ringPos = wgsHap1.range),
                        list(regionDef = wgs.hom, ringPos = hom.range))
      
      regionPlot <- createRingPlot.region(ringTable ,
                                          xmin = min(c(wgsHap1.range, hom.range, misalignHap1.range, mismatch.range)),
                                          xmax = max(c(wgsHap1.range, hom.range, misalignHap1.range, mismatch.range)),
                                          mapFile = mapFile,
                                          readMapFile = F,
                                          chrSepSize = chrSepSize,
                                          ringSepsize = ringSepsize,
                                          sep.code = 99,
                                          writeOutput = F,
                                          outfolder = NULL,
                                          outpath = "snpMatch_refGenome",
                                          returnOutput = T)
      
      regionPlot <- rbind(regionPlot, mismatchPlot)
      
      wgsHap2.alignRing <- NULL
      
      if(analysisT == "refInd"){
        print("** main step 4 skipped: no second haploid WGS to analyze **")
      } else {
        print("** main step 12 skipped: no second haploid WGS to analyze **")
      }
    }
    
    if(writeOutput){
      write.csv(regionPlot, file = paste0(outpath, "_ringPlot.csv"), row.names = F)
      write.csv(wgsHap1.alignRing, file = paste0(outpath, "_hap1_alignmentPlot.csv"), row.names = F)
      if(!is.null(wgsHap2.snpProf)){
        write.csv(wgsHap2.alignRing, file = paste0(outpath, "_hap2_alignmentPlot.csv"), row.names = F)
      }
    }
    
    if(analysisT == "refInd"){
      print("** main step 5: creating plot **")
    } else {
      print("** main step 13: creating plot **")
    }
    colorDef <- data.frame(matrix(c(c(-1,0,1,2,11,99),c(misOrigin.col, mismatch.col, hap1.col,hap2.col,hom.col, ringOutlineCol),c("Origin Mismatch","Genotype Inconsistency", hap1.label, hap2.label, paste0("Homozygosity > ",hom.threshold,"cM"), "")), ncol = 3), stringsAsFactors = F)
    colnames(colorDef) <- c("state", "color", "legend" )
    
    if(writeOutput){
      write.csv(colorDef, file = paste0(outpath, "_colorDef.csv"), row.names = F)
    }
    
    ringPlot <- createRingPlot.plot(regionTable = regionPlot,
                                    colorDef = colorDef,
                                    missAlignTable.hap1 = wgsHap1.alignRing,
                                    missAlignTable.hap2 = wgsHap2.alignRing,
                                    readRegionT = F,
                                    readColorDef = F,
                                    readMissAlignT.hap1 = F,
                                    readMissAlignT.hap2 = F,
                                    unalignCol = unalign.col,
                                    unanchCol = unanch.col,
                                    misalignCol = misalign.col,
                                    writeOutput = writeOutput,
                                    outfolder = NULL,
                                    outpath = outpath,
                                    plot.width = plot.width,
                                    plot.units = plot.units,
                                    plot.dpi = plot.dpi,
                                    returnOutput = T)
    
    if(analysisT == "all"){
      if(returnOutput){
        if(!is.null(wgsHap2.fasta)){
          return(list(wgsHap1.blastResults = wgsHap1.blastResults,
                      wgsHap2.blastResults = wgsHap2.blastResults,
                      wgsHap1.snpAlleles = wgsHap1.snpAlleles,
                      wgsHap2.snpAlleles = wgsHap2.snpAlleles,
                      wgsHap1.snpProf = wgsHap1.snpProf,
                      wgsHap2.snpProf = wgsHap2.snpProf,
                      wgsProf = wgsProf,
                      wgsVsSNPprof.mismatch = wgsVsSNPprof.mismatch,
                      wgsHap1.wgsHapOrigin = wgsHap1.wgsHapOrigin,
                      wgsHap2.wgsHapOrigin = wgsHap2.wgsHapOrigin,
                      wgs.hom = wgs.hom,
                      wgsHap1.alignErrSumm = wgsHap1.alignErrSumm,
                      wgsHap2.alignErrSumm = wgsHap2.alignErrSumm,
                      regionPlot = regionPlot,
                      wgsHap1.alignRing = wgsHap1.alignRing,
                      wgsHap2.alignRing = wgsHap2.alignRing,
                      ringplot = ringPlot))
        } else {
          return(list(wgsHap1.blastResults = wgsHap1.blastResults,
                      wgsHap1.snpAlleles = wgsHap1.snpAlleles,
                      wgsHap1.snpProf = wgsHap1.snpProf,
                      wgsProf = wgsProf,
                      wgsHap1.wgsHapOrigin = wgsHap1.wgsHapOrigin,
                      wgs.hom = wgs.hom,
                      wgsHap1.alignErrSumm = wgsHap1.alignErrSumm,
                      regionPlot = regionPlot,
                      wgsHap1.alignRing = wgsHap1.alignRing,
                      ringplot = ringPlot))
        }
      }  
    } else if (analysisT == "blast"){
      if(returnOutput){
        if(!is.null(wgsHap2.fasta)){
          return(list(wgsHap1.blastResults = wgsHap1.blastResults,
                      wgsHap2.blastResults = wgsHap2.blastResults,
                      wgsHap1.snpAlleles = wgsHap1.snpAlleles,
                      wgsHap2.snpAlleles = wgsHap2.snpAlleles,
                      wgsHap1.snpProf = wgsHap1.snpProf,
                      wgsHap2.snpProf = wgsHap2.snpProf,
                      wgsProf = wgsProf))
        } else {
          return(list(wgsHap1.blastResults = wgsHap1.blastResults,
                      wgsHap1.snpAlleles = wgsHap1.snpAlleles,
                      wgsHap1.snpProf = wgsHap1.snpProf,
                      wgsProf = wgsProf))
        }
      } 
    } else if (analysisT == "refInd") {
      if(returnOutput){
        if(!is.null(wgsHap2.snpProf)){
          return(list(wgsVsSNPprof.mismatch = wgsVsSNPprof.mismatch,
                      wgsHap1.wgsHapOrigin = wgsHap1.wgsHapOrigin,
                      wgsHap2.wgsHapOrigin = wgsHap2.wgsHapOrigin,
                      wgs.hom = wgs.hom,
                      wgsHap1.alignErrSumm = wgsHap1.alignErrSumm,
                      wgsHap2.alignErrSumm = wgsHap2.alignErrSumm,
                      regionPlot = regionPlot,
                      wgsHap1.alignRing = wgsHap1.alignRing,
                      wgsHap2.alignRing = wgsHap2.alignRing,
                      ringplot = ringPlot))
        } else {
          return(list(wgsVsSNPprof.mismatch = wgsVsSNPprof.mismatch,
                      wgsHap1.wgsHapOrigin = wgsHap1.wgsHapOrigin,
                      wgs.hom = wgs.hom,
                      wgsHap1.alignErrSumm = wgsHap1.alignErrSumm,
                      regionPlot = regionPlot,
                      wgsHap1.alignRing = wgsHap1.alignRing,
                      ringplot = ringPlot))
        }
      }  
    }
  }
}
```
