## Supplementary material for "Whole genome sequence improvement with pedigree information and reference genotypic profiles, demonstrated in outcrossing apple": Suppl File S1 - R functions v1: ReadMe R functions.txt

Supplementary material to Vanderzande et al. “Approaches for whole genome sequence improvement through the use of pedigree information and reference DNA profiles demonstrated in outcrossing apple”

Created R functions to run a WGS analysis using reference map and genotpypic data.

Two R packages are needed to run the function:
tidyverse (of which ggplot2, dplyr, stringr, and tibble are needed)
pbapply
