## Supplementary material for "Whole genome sequence improvement with pedigree information and reference genotypic profiles, demonstrated in outcrossing apple": Suppl File S2 - Demo Input Files: ReadMe Demo Input Files.txt

Supplementary material to Vanderzande et al. “Approaches for whole genome sequence improvement through the use of pedigree information and reference DNA profiles demonstrated in outcrossing apple”

Example input files provided to run with the developed R functions

Warning!: The phased Gala WGS should also be downloaded as input file (Sun et al., 2020). It is accessible, for example, under the "assembly" section of https://www.rosaceae.org/Analysis/10816131  

Demo 20K_GalaA_blast.out: Example output file from BLAST against the first haplome of Gala v1.0. Outcome of SNPs of the first three linkage groups of the iGL map are provided
Demo 20K_GalaA_blast.out: Example output file from BLAST against the second haplome of Gala v1.0. Outcome of SNPs of the first three linkage groups of the iGL map are provided
Demo 20K_mapFile iGL.csv: Example mapFile needed for the analysis. The first three linkage groups of the iGL map are provided
Demo 20K_mhaplotypes.csv: Example of phased data file needed for the analysis. Format based on the mhaplotypes.csv output from FlexQTL. Phased data of Gala provided for first three linkage groups of the iGL map
Demo 20K_snpInfoProbes.csv: Example snpInfo file needed for the analysis. Contains position of SNP within the BLASTed sequence. In case of probes here, the position of the SNP is immediately downstream of the sequence, which is also allowed. Information is provided for the first three linkage groups of the iGL map
Demo ParFile_Gala_Probes_bn.csv: Example parFile for use with the 'PhasedWgsVsSNP' wrapper R function. Contains all parameter settings to do a WGS analysis using reference genetic data
