## Supplementary material for "Whole genome sequence improvement with pedigree information and reference genotypic profiles, demonstrated in outcrossing apple": Suppl File S3 - Demo Output Files: ReadMe Demo Output Files.txt

Supplementary material to Vanderzande et al. “Approaches for whole genome sequence improvement through the use of pedigree information and reference DNA profiles demonstrated in outcrossing apple”

Supplementary Material S2: Example output files from the developed R functions. Generated with the provided input files from Suppl M1.

## Output files from the analysis of the first haplome
Demo_hap1_alignmentPlot.csv: Information used to create the alignment plot with provided ring width and location. It provides per user-defined window what proportion of markerss did not align to the first haplome of the WGS, aligned to unanchored regions, and aligned to regions inonsistent with the genetic map, adjusted for ring width and position. 
Demo_hap1_AlignSumm.csv: Provides, per user-defined window, the unaltered proportions of markers did not align to the first haplome of the WGS, aligned to unanchored regions, and aligned to regions inonsistent with the genetic map
Demo_hap1_alleleOrigSumm.csv: Provides parental origin of WGS segments for the first haplome based on the phased reference individual. "1" and "2" are used for alleles originating from the first and second parent of the reference individual, respectively.
Demo_hap1_dnaProfile.csv: Marker alleles retrieved from the first haplome of the WGS, ordered accroding to the provided genetic map
Demo_hap1_mapMismatch.csv: Provides list of markers whose determined position in the first haplome of the WGS do not correspond with the genetic map.
Demo_hap1_MulitHits_MultiChr.csv: Provides list of markers and their BLAST hits that aligned to multiple chromosomes of the first haplome (and for which no single location in the haplome could be determined)
Demo_hap1_MulitHits_SingleChr.csv: Provides list of markers and their BLAST hits that aligned to locations within a single chromosome of the first haplome (and for which no single location in the haplome could be determined)
Demo_hap1_UniqueHits.csv: Provides list of markers and their BLAST hits for which a single location in the first haplome of the WGS could be determined
Demo_hap1_wgsAlleles.csv: Provides list of alleles in the first haplome of the WGS of markers for which a single location could be determined. 

## Output files from the analysis of the second haplome (not present when only one haplome is provided)
Demo_hap2_alignmentPlot.csv: Information used to create the alignment plot with provided ring width and location. It provides per user-defined window what proportion of markers did not align to the second haplome of the WGS, aligned to unanchored regions, and aligned to regions inonsistent with the genetic map, adjusted for ring width and position.
Demo_hap2_AlignSumm.csv: Provides, per user-defined window, the unaltered proportions of markers did not align to the second haplome of the WGS, aligned to unanchored regions, and aligned to regions inonsistent with the genetic map
Demo_hap2_alleleOrigSumm.csv: Provides parental origin of WGS segments for the second haplome based on the phased reference individual. "1" and "2" are used for alleles originating from the first and second parent of the reference individual, respectively.
Demo_hap2_dnaProfile.csv: Marker alleles retrieved from the second haplome of the WGS, ordered accroding to the provided genetic map
Demo_hap2_mapMismatch.csv: Provides list of markers whose determined position in the second haplome of the WGS do not correspond with the genetic map.
Demo_hap2_MulitHits_MultiChr.csv: Provides list of markers and their BLAST hits that aligned to multiple chromosomes of the second haplome (and for which no single location in the haplome could be determined)
Demo_hap2_MulitHits_SingleChr.csv: Provides list of markers and their BLAST hits that aligned to locations within a single chromosome of the second haplome (and for which no single location in the haplome could be determined)
Demo_hap2_UniqueHits.csv: Provides list of markers and their BLAST hits for which a single location in the first haplome of the WGS could be determined
Demo_hap2_wgsAlleles.csv: Provides list of alleles in the second haplome of the WGS of markers for which a single location could be determined.

## Output files from the analysis of both haplomes (or single haplome when only one is provided)
Demo_colorDef.csv: Color definition file created from parameter file. Needed adjust colors and run the ringplot function separately without need of redoing whole analysis.
Demo_homSumm: Provides list of homozygous regions in the reference individual larger than a user-defined length.
Demo_IdentityReport.csv: Provides a proportion of markers whose WGS-based genotype matched that of the reference data. In case of a single haplome, it provides markers for which there is a conflict (i.e., 'A' allele in the WGS and 'BB' genotype in the reference individual). Also provides a list of markers whose genotype were in conflict with the reference data
Demo_parametersUsed.csv: Provides a list of all parameter values used for the analysis. (to later retrace what was done)
Demo_ringPlot.csv: Information used to generate the ringplot, based on user-defined ring locations and widths. Contains info on location of segements of continuous parental origin, homozygous regions, and SNPs whose WGS genotype conflict with the reference data. all adjusted for their respective ring location and width
Demo_ringplot.pdf: Ringplot created based on information from Demo_hap1_alignmentPlot.csv, Demo_hap2_alignmentPlot.csv (if generated), and Demo_ringPlot.csv as well as colors provided in the parameter file.
