## Supplementary figures and images for "Whole genome sequence improvement with pedigree information and reference genotypic profiles, demonstrated in outcrossing apple"

### Demo_ringplot.pdf

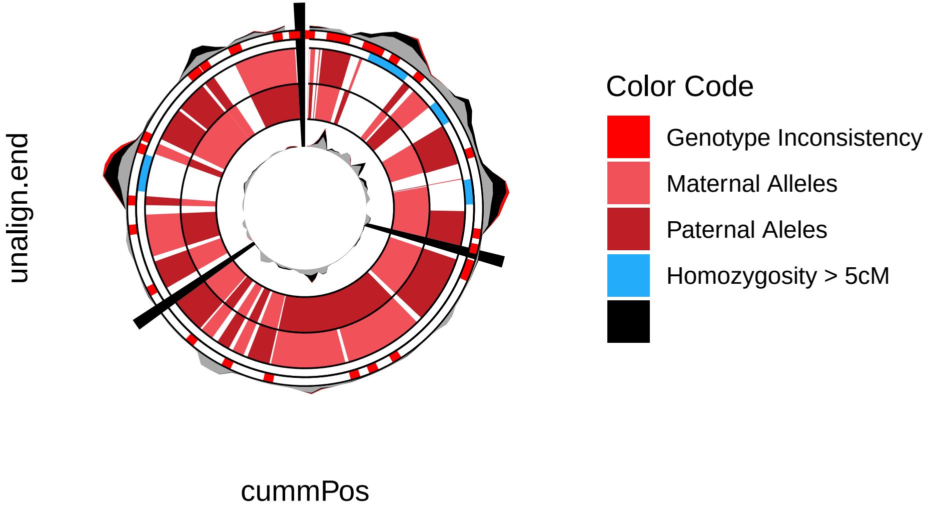
